# Temporal and Spatial Resolution of a Protein Quake that Activates Hydrogen Tunneling in Soybean Lipoxygenase

**DOI:** 10.1101/2022.03.31.486235

**Authors:** Jan Paulo T. Zaragoza, Adam R. Offenbacher, Shenshen Hu, Christine L. Gee, Zachary M. Firestein, Natalie Minnetian, Zhenyu Deng, Flora Fan, Anthony T. Iavarone, Judith P. Klinman

## Abstract

The enzyme soybean lipoxygenase (SLO) provides a prototype for deep tunneling mechanisms in hydrogen transfer catalysis. This work combines room temperature X-ray studies with extended hydrogen deuterium exchange experiments to detect a radiating cone of aliphatic side chains that extends from the iron active site of SLO to the protein-solvent interface. Employing eight variants of SLO, nanosecond fluorescence Stokes shifts have been measured using a probe appended to the identified surface loop. We report a remarkable identity of the enthalpies of activation for the Stokes shifts decay rates and the millisecond C-H bond cleavage step that is restricted to side chain mutants within the identified thermal network. While the role of dynamics in enzyme function has been predominantly attributed to a distributed protein conformational landscape, these new data implicate a thermally initiated, cooperative protein quake as the source of the activation of SLO. These findings indicate a direct coupling of distal protein motions surrounding the exposed fluorescent probe to active site motions controlling catalysis.

## Introduction

The contribution of protein scaffold motions to the modulation of reaction barrier height and shape remains a major unresolved component in our ability to describe the physical properties underlying enzyme catalysis. There is a growing acceptance of the role of protein conformational sampling in the determination of reaction rate and regulation of enzyme catalysis, formulated in the context of a Boltzmann distribution of thermodynamically equilibrated protein sub-states that are primarily populated by low energy configurations accessible near room temperature (1-3). As expected from the anisotropic topology of almost all folded proteins, site specificity in the achievement of functionally relevant protein sub-states is apparent from both computational analyses (4) and room temperature X-ray crystallographic measurements (5-7). Though more elusive, the involvement of real-time couplings of protein dynamical motions to the chemical steps of catalysis has also been addressed through the application of transition path sampling (8-10). This approach leads to the computation of local atomic motions that accompany reaction barrier crossings, but has not yet been adopted to include more widespread and remote protein motions.

A related but somewhat different context for our understanding of the role of the protein scaffold in catalysis comes from the central role of thermal activation among the vast majority of enzyme catalyzed reactions, with only a small, albeit important, handful of enzymes requiring photochemical initiation (e.g., bacterial reaction centers (11, 12), protochlorophyllide oxidoreductase (13), DNA photolyase and cryptochromes (14), and fatty acid photodecarboxylase (15)). The temperature dependence of enzyme-catalyzed reaction, in certain instances, has been analyzed in terms of changes in heat capacity (ΔC^‡^_p_), attributed to a reduction in active site flexibility at the point of barrier crossings (16, 17). Temperature dependent hydrogen-deuterium exchange coupled to mass spectrometry (TDHDX-MS) has recently emerged as a broadly accessible experimental tool to interrogate heat induced changes in local protein flexibility, which when coupled to function-altering protein mutations provides spatial resolution of activated networks that are found to lead from protein-solvent interfaces to enzyme active sites (18-22). While the thermal activation of enzymes can be rationalized as a general redistribution of equilibrated protein states toward increasingly active configurations with increasing temperature, the growing identification of reaction specific thermal networks that are distinctive from each other and independent of protein scaffold conservation points toward scaffold-embedded heat conduits as a source of dynamical excitation (23-30).

The concept of protein quakes arose initially during studies of the photodissociation of CO from myoglobin and remains an active area of research (31-33). With the development of time-resolved X-ray scattering and free-electron laser (XFEL) techniques, it has become possible to monitor the spatial and time dependencies of anisotropic heat dissipation from a protein interior to the solvent (34-36). In the present work, we turn this concept around to focus on the role of anisotropic heat movement inwards, from a protein solvent interface toward the site of substrate binding and catalysis. It has been hypothesized that the transmission of thermal energy from the solvent bath to the protein surface provides the energy required to overcome the chemical reaction barrier (3, 37, 38). Addressing this question has been greatly facilitated by the large body of emergent evidence for quantum mechanical tunneling in enzymatic hydrogen transfer reactions. The adaptation of Marcus-like models for long range electron transfer to C–H cleavage reactions provides a powerful formalism that can accommodate the multi-dimensional properties of hydrogen tunneling at room temperature (39-45). In particular, the resulting separation of the protein environmental reorganization terms that transiently create the energetic degeneracy and internuclear distances that enable effective H-tunneling from the inherently temperature-independent probability of wave function overlap refocuses structure-function-dynamics studies on the protein scaffold itself and allows its deconvolution from the primary chemical bond cleavage event.

Soybean lipoxygenase (SLO) emerged several decades ago as a paradigmatic system for exploring the properties of thermally activated tunneling in biology (46). SLO catalyzes the production of fatty acid hydroperoxides at (Z,Z)-1,4-pentadienoic moieties via a rate-limiting proton coupled electron (net hydrogen atom) transfer from substrate to an Fe^III^(OH) cofactor (Fig. 1*A*). The physiological substrate 9,12-(Z,Z)-octadecadienoic acid (linoleic acid, LA) is converted to 13-(S)-hydroper-oxy-9,11-(Z,E)-octadecadienoic acid (13-(S)-HPOD). The resulting oxygenated lipids and their byproducts are essential for signaling structural and metabolic changes in the cell, including seed growth and plant development (47). Native SLO catalysis exhibits a fairly weak temperature dependence (*E*_a_ ∼ 2.0 kcal/mol), a small Arrhenius prefactor (A_H_ < 10^5^ s^-1^), and a greatly inflated kinetic isotope effect (KIE) on *k*_cat_ (^D^*k*_cat_ = 81) (40, 48) that rises to ^D^*k*_cat_ = 500 – 700 for an enzyme variant that has been mutated at two hydrophobic side chains surrounding the reactive carbon of bound substrate (49). The aggregate structural and kinetic properties of SLO played a key role in the development of vibronically non-adiabatic analytical expressions that are now routinely applied to explain deep tunneling reactions that occur near room temperature (40, 42-44, 50, 51). SLO was also the first protein to be subjected to a combined analysis of the time-, temperature- and mutation-dependent properties of hydrogen deuterium exchange as analyzed by mass spectrometry (19). The resulting identification of a site selective thermal pathway, originating at a solvent-exposed loop and extending toward the buried iron cofactor and substrate binding site, provides a unique opportunity to explore the structural and dynamical features that control C-H activation in enzyme catalysis. Evidence is presented herein for a long-range protein quake as the source of thermal activation during the rate limiting hydrogen transfer step of SLO.

**Fig. 1.**
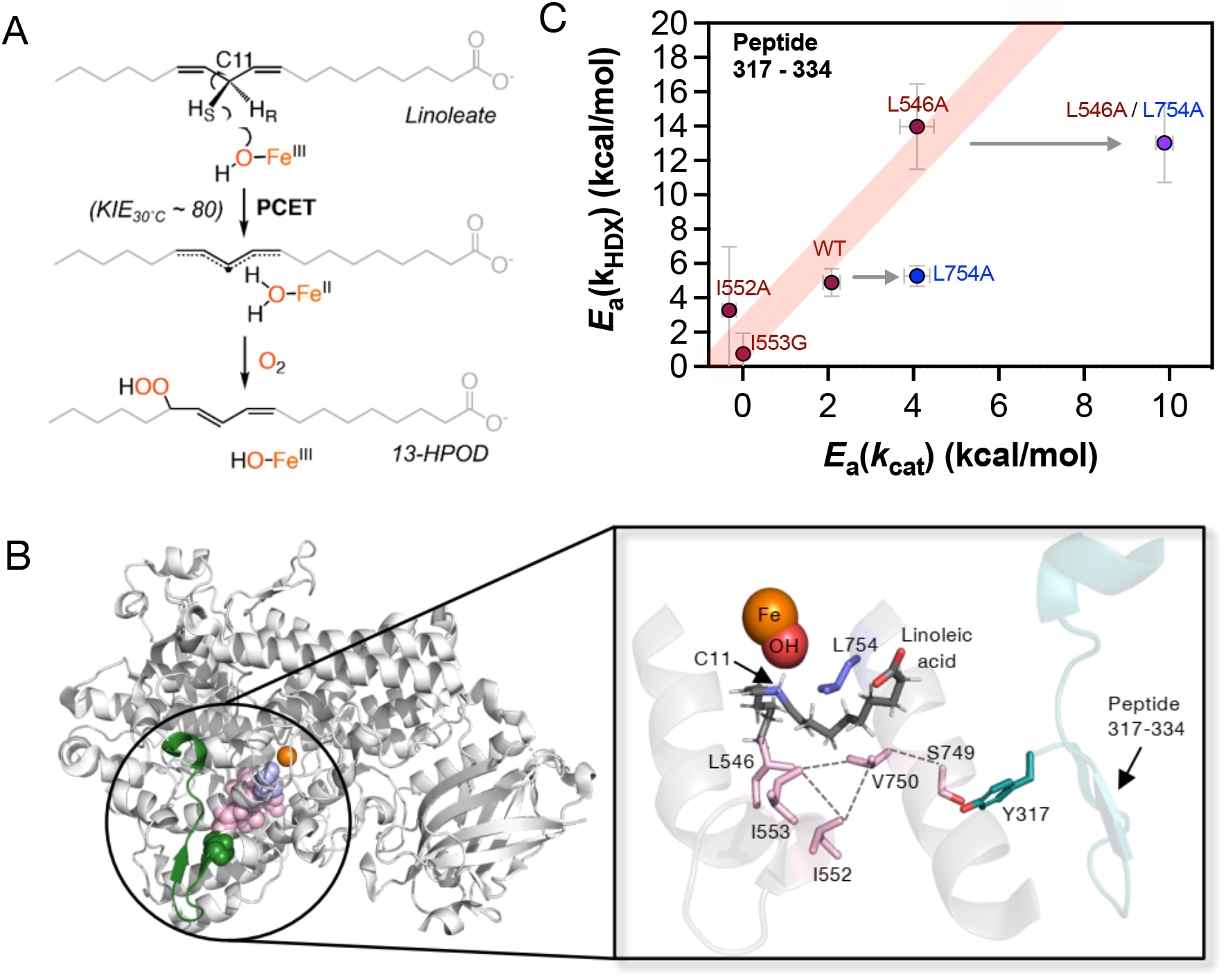
(*A*) Rate-limiting hydrogen transfer reaction from linoleic acid (LA) catalyzed by an Fe^III^(OH) cofactor in SLO (46). (*B*) X-ray structure of SLO (PDB:3PZW) depicting the active site Fe, and the network of residues (pink) and thermally activated surface loop (residues 317-334, in dark blue-green) identified from previous HDX-MS experiments (Ref. 19). Inset: The bound linoleic acid substrate (gray) is modeled from previous calculations (51) and overlaid in this structure. Leu754, which is outside the identified network, is shown in purple. The peptide 317-334 is labeled in blue-green. (*C*) Temperature-dependent HDX-MS analysis. Relationship between the HDX enthalpic barrier (E_a_ for *k*_HDX(avg)_) at peptide 317-334 and the enthalpic barrier for hydrogen tunneling (E_a_ for *k*_cat_). New data for I552A are shown and compared with data for other mutants from Ref. (52).

## Results

### The catalytically linked thermal network in SLO extends beyond the TDHDX-MS defined pathway

The originally defined thermal pathway for SLO (19) was found to be comprised of three regions: (i) the substrate binding site in proximity to the Fe^III^(OH) cofactor, (ii) an intermediate region comprised of Ile552 and Val750, and (iii) a loop that interacts directly with solvent (Fig. 1*B*). The aliphatic residues Ile552 and Val750 are positioned to act as mediators between the active site side chains Leu546 and Ile553 and the residues that extend to the surface loop. Of particular interest, neither Ile552 nor Val750 is in direct contact with the reactive carbon, raising the question of their importance and impact to catalytic proficiency. To address this issue, we examined the effect of alanine substitution at Ile552 and Val750, obtaining a suite of kinetic parameters (Table 1). While the magnitude of the difference in the enthalpy of activation for the reaction with H- vs. D-labeled LA substrate, ΔE_a_, is reduced for V750A, its value is within the experimental error of WT and the value of ΔE_a_ for I552A is unchanged. An increase in ΔE_a_ has been shown to map to a loss of precision in the local positioning of the reactive carbon (C11 of the LA substrate) for SLO (53), as well as for a wide range of H-transfer enzymes (54). In accordance with this behavior, evolutionary selection was recently applied to a primitive form of dihydrofolate reductase (DHFR), showing a progressive decrease in ΔE_a_ as the evolved enzyme approached the activity of native DHFRs (55). From this study of I552A and V750A, we conclude that ΔE_a_ is largely unaffected, with the impact of such mutations appearing primarily as a reduction in the enthalpy of activation (E_a_(H)) that is accompanied by reduction of the first-order rate constant (*k*_cat_) at 30 °C. The changes in E_a_(H) suggest a perturbation (loosening) of structure away from the configuration that is optimal for catalytic turnover in WT, with the impact of I552A exceeding that of V750A.

**Table 1.** Comparative kinetic parameters for Ala variants at positions 552 and 750 within the thermal network of SLO^*a*^

| SLO variant | $k_{cat}$, s<sup>-1</sup> | $^Dk_{cat}$ | $E_a$,<br>kcal/mol | $\Delta E_a$,<br>kcal/mol | Reference |
| --- | --- | --- | --- | --- | --- |
| WT | 297 (10) | 81 (5) | 2.1 (0.2) | 0.9 (0.2) | Ref. (40) |
| I552A | 81 (2)<sup>b</sup> | 60 (2)<sup>b</sup> | -0.3 (0.2) | 1.2 (0.5) | This work<sup>a</sup> |
| V750A | 218 (8)<sup>b</sup> | 69 (4)<sup>b</sup> | 1.0 (0.4) | 0.3 (0.4) | This work<sup>a</sup> |

<sup>a</sup>Temperature dependent kinetics data had been reported (56) but were not previously analyzed for $E_a$ and $\Delta E_a$ . The full range of additional mutants formerly characterized in this fashion is summarized in *SI Appendix*, Table S1.
<sup>b</sup>For comparison under similar protein and substrate purification procedures (19), data for WT were recollected using the enzyme preparation of the present study, generating $k_{cat} = 359$ (7) s<sup>-1</sup>, $^Dk_{cat} = 57$ (2), $E_a = 2.4$ (0.2) kcal/mol, and $\Delta E_a = 1.1$ (0.1) kcal/mol.

With the goal of more fully understanding the contribution of residues Ile552 and Val750 to the thermal pathway in SLO, additional independent experimental directions were pursued. First, TDHDX-MS was extended to I552A, resulting in the same set of non-overlapping peptides as obtained for WT and other variants (19, 52) (*SI Appendix*, Fig. S1). The majority of the I552A-derived peptides display nearly identical exchange behavior to WT across the 5 temperatures (10, 20, 25, 30, and 40 °C) analyzed (*SI Appendix*, Data S1). The peptide of particular interest is 317-334 that represents the surface loop that was previously shown (19) to undergo correlated alterations in the activation energy for HDX E_a_(*k*_HDX_) and E_a_(*k*_cat_) when interrogating the variants I553G and L546A in relation to WT. The new data for I552A indicate an E_a_(*k*_HDX_) = 3.2(3.7) kcal/mol, within experimental error of zero, analogous to its E_a_(*k*_cat_) = -0.3(0.2) kcal/mol) (*SI Appendix*, Table S2). As shown in Fig. 1*C* (red line) these data for I552A fall on the original correlation of E_a_(*k*_HDX_) and E_a_(*k*_cat_), in support of the inclusion of Ile552 within the defined thermal pathway. We note that mutations at Leu754 fall off the line correlating Ea(*k*_*HDX*_) to Ea(*k*_*cat*_) (Fig. 1*C*), and this feature serves as a valuable control for time and temperature dependent Stokes Shift decay analyses presented below. We also noted an unusually large perturbation from I552A on the TDHDX-MS of peptides 541-554 and 555-565 residing near the inferred substrate portal, with peptide 541-554 of I552A indicating a decrease in E_a_(*k*_HDX_) from 23.7(2.0) kcal/mol for WT to 5.64(0.5) kcal/mol (*SI Appendix*, Fig. S2); these data are consistent with an earlier report of an increase in *k*_off_ values for substrate release in I552A, attributed to a loosening of the substrate portal that is adjacent to the active site (56).

Room temperature (RT, 277 – 300 K) X-ray studies have been shown to reveal multiple, weakly populated sub-states within a protein conformational ensemble (5). The multi-conformer models obtained from qFit (57) uncover local anisotropic protein dynamics information that is supported by NMR measurements, and reflects multiscale motions of the protein backbone and side chains (7). Close inspection of the RT electron densities for WT SLO published earlier indicated a number of alternate side chain conformers for key active site residues (19). Herein we first applied RT X-ray studies (T = 300 K) to the activity altering variants, I553G and L546A (52), finding that the I553G variant decreases the conformational space accessed by its proximal Leu546 side chain (*SI Appendix*, Fig. S3). The L546A mutation, on the other hand, leaves Ile553 unperturbed while decreasing the electron density of an alternate conformer for a second proximal residue Ile552. With this observation, we proceeded to examine the impact of both I552A and V750A mutations on nearby residues via RT X-ray structural analysis (*SI Appendix*, Table S3). It is expected that changes in side chain conformations following site specific mutagenesis may radiate throughout the protein, however the impacts of I552A and V750A are found to be concentrated within a region that neighbors the TDHDX-MS detected thermal pathway (Fig. 2). The I552A and V750A mutations show reciprocal differences in alternate conformers at the two positions. Specifically, V750A leads to a loss of alternate conformer in Ile552, while I552A results in a different configuration of the Val750 side chain from that of WT. Additionally, I552A and V750A mutations result in a loss of alternate conformers for the adjacent Leu546 and the more remote Ile746 and Leu262 side chains (Fig. 2*A*).

**Fig. 2.**
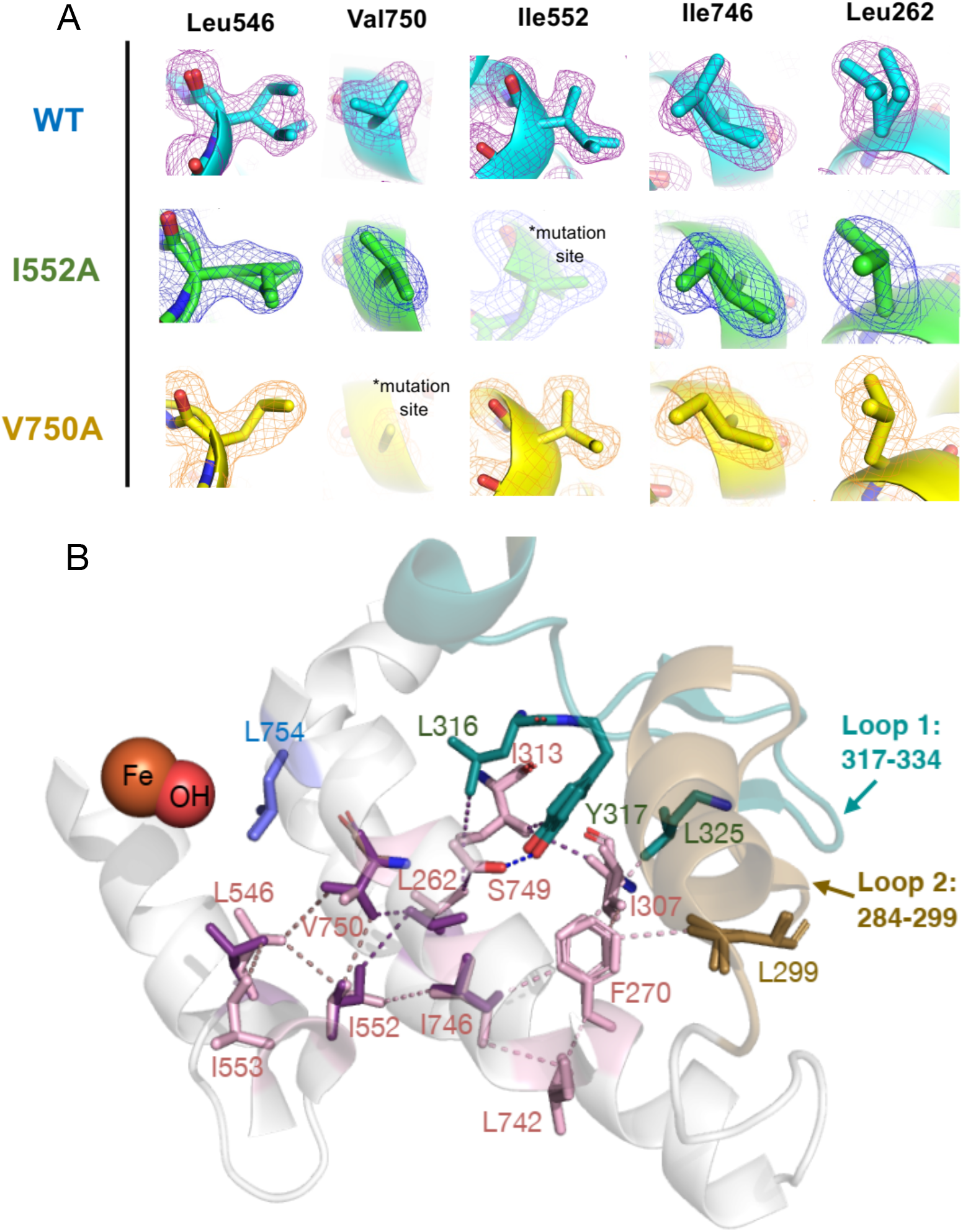
Room temperature X-ray structure analysis. (*A*) Impact of I552A and V750A mutation on the 300 K X-ray structure electron densities of their respective side chains as well as those of Leu546, Ile746 and Leu262. (*B*) Connectivity of residues from the Fe active site to the surface loops (loop 1: 317-334 (blue-green), loop 2: 284-299 (brown)) in the 300 K X-ray structure of WT SLO (PDB:5T5V). Multiple side chain conformers along the thermal network are represented in pink and purple.

The above analyses provide strong support for the role of both Ile552 and Val750 as mediators between the active site side chains of SLO and an extended thermal network that terminates at the protein-water interface. As a guide for visualizing the entire network, side chain interactions have been traced from Leu546 to the surface loops comprised primarily by the region of protein represented by amino acids 317-334 and to a lesser extent the region 284-299, a second peptide shown to undergo correlated motions in the E_a_ for HDX-MS and catalysis (19) (Fig. 2*B*). Using a cut-off of 4 Å for van der Waals interactions between adjacent side chains, an extended network of amino acids is detected beyond the TDHDX-MS derived pathway. In particular, Ile552 is seen to act as a central lynchpin, exhibiting steric interactions with Leu546, Val750, Leu262, and Ile746. As shown in Fig. 2*B*, a clearly defined, cone-shaped region of protein with activity-related alternate conformers connects the active site to the solvent exposed loop. The abundance of hydrophobic residues in this extended network led us to consider other possible clustering of hydrophobic residues in SLO. It has been proposed that the sidechains of Ile, Leu, and Val often form hydrophobic clusters (58, 59) which serve as cores of stability and prevent intrusion of water molecules. The hydrophobic clusters in SLO (*SI Appendix*, Fig. S4) were computed using a CSU algorithm (60). Of considerable interest for how thermal networks may be constructed, the largest calculated hydrophobic cluster in SLO is comprised of 32 residues and includes the Ile839 ligand to the Fe center, the active site residues that are in contact of the linoleic acid substrate (Ile553, Leu546, Leu754) and all of the residues that are experimentally assigned to a thermal activation pathway.

### Demonstration of a dynamically activated thermal network in SLO that stretches from the protein/solvent interface to the active site

To examine dynamical motions within the thermal loop in SLO, the fluorescent probe 6-bromoacetyl-2-dimethylaminonaphthalene (BADAN, BD) was incorporated at position 322 at the tip of the identified thermal network (Fig. 3*A*) and at a remote control position 596 via thiol modification (61). Acetyl-2-dimethylaminonaphthalene-based dyes such as BADAN are used as sensors to probe changes in local solvation dynamics in proteins following photo-excitation (62). The main mechanism for fluorescence modulation is through interactions of the surrounding solvent with the electronically excited state of the probe (63), which in the case of BADAN leads to polarization of an N,N-dimethylamino group and a carbonyl substituent. Nearby Trp residues in proteins have been shown to quench BADAN fluorescence via electron transfer (64). In Q322C-BD SLO, the nearest Trp residue, Trp340, to the BADAN probe at position 322 is 30-40 Å away. Molecular dynamics simulations of Q322C-BD show that this distance does not change significantly (*SI Appendix*, Fig. S5), making artifactual quenching of the BADAN probe at the 322 position an unlikely complication. In an earlier preliminary work (61), the temperature dependence of the BADAN Stokes shift decay was observed to be negligible at a control loop at position 596. When the probe was appended to the end of the identified thermal loop at position 322, the enthalpies of activation for Stokes shifts matched that for catalytic C-H activation E_a_(*k*_cat_). The striking observation of an exact correspondence between the enthalpic barrier for a millisecond catalytic step and for a nanosecond Stokes shift decay process suggested a direct coupling of distal protein motions surrounding the BADAN probe to active site motions controlling catalysis in the WT enzyme.

**Fig. 3.**
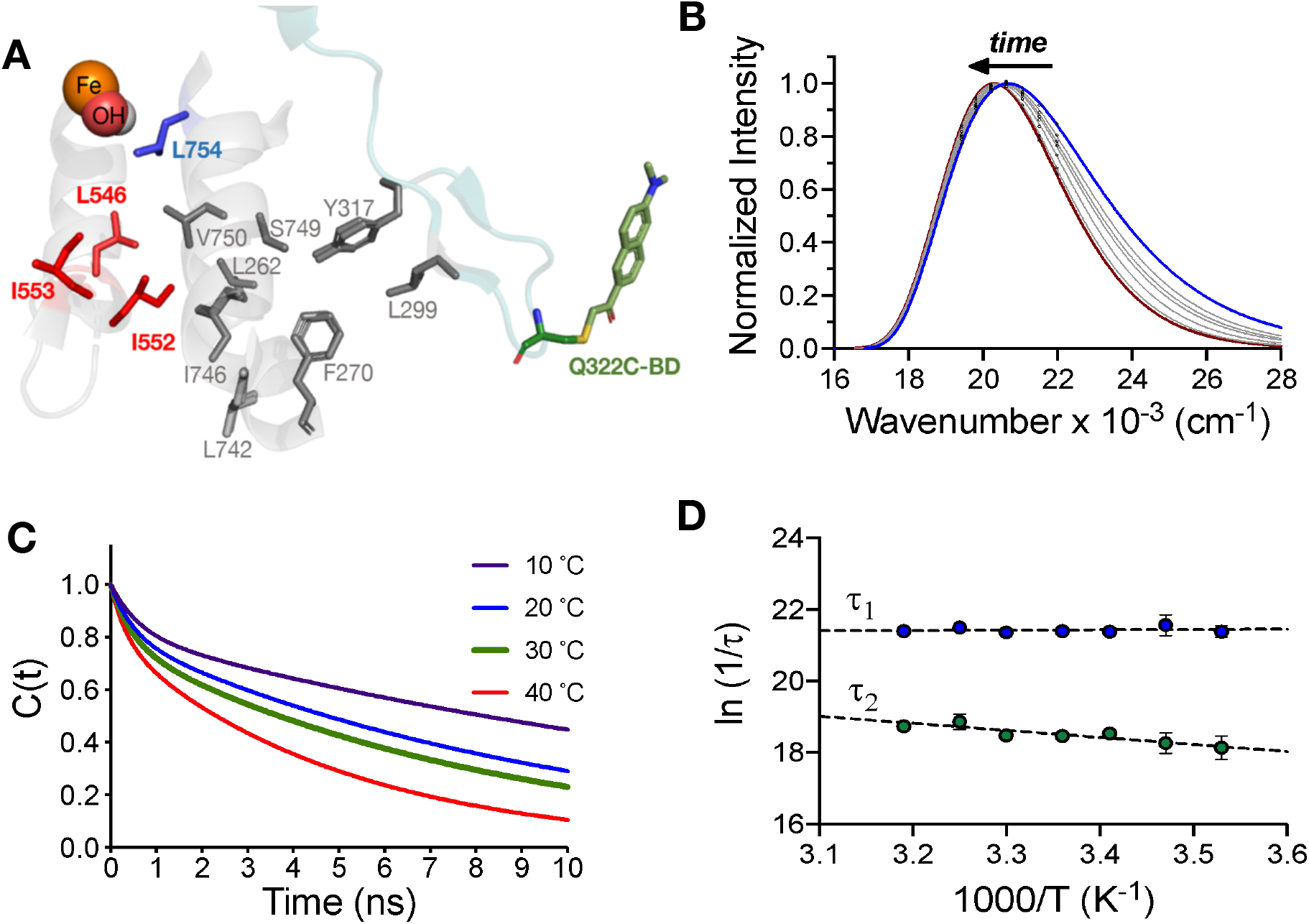
Temperature-dependent nanosecond fluorescence Stokes shift analysis of a fluorescent probe on a remote surface loop in SLO. (*A*) Location of the mutated active site residues Ile553, Leu546, Ile552, and Leu754 in relation to the thermally-activated loop (317-334, blue-green), and the BADAN-labeled Q322C residue (green). The BADAN orientation is based on the optimized structure from Ref. (61). (*B*) Representative time-resolved emission spectra (TRES) at 30 °C, pH 9.0 for the L546A SLO mutant. (*C*) Representative time and temperature dependence of the Stokes shift decay rates of L546A. (*D*) Arrhenius plots for the nanosecond Stokes shift decays (1/τ_1_, 1/τ_2_) of L546A.

In this work we pursue Stokes shift measurements with SLO using a wide range of enzyme variants with distinctive trends in E_a_(*k*_cat_) and E_a_(*k*_HDX_). Four of the variants chosen for study are either in contact with bound substrate and/or with each other: I553G, I553A, L546A, and I552A (Fig. 3*A*). Substitution of residues at positions 553 and 546 in the substrate binding pocket leads to perturbations that either increase (L546A), decrease (I553G), or retain (I553A) the same E_a_(*k*_*cat*_) as WT (40, 65) (*SI Appendix*, Table S1). Ile552, which is located proximal to Leu546, but is not in direct contact with substrate, produces a decreased E_a_(*k*_*cat*_) relative to WT (Table 1). The L754A variant was also investigated as internal control. While Leu754 is positioned on an opposite face of the reactive carbon of substrate from Leu546, and shows an elevated enthalpic barrier for catalysis, TDHDX-MS measurements (52) indicated little or no change relative to WT on E_a_(*k*_HDX_), in support of Leu754 residing outside the pathway for thermal activation (Fig. 1*C*). The impacts of L754A on rate and catalytic activation energy, in turn, have been attributed to an expanded active site that incorporates additional water molecules, while altering the active site microenvironment and Fe active site geometry (52). Stokes shift controls involving double mutations with L754A as the parent were also interrogated, involving I553A/L754A and L546A/L754A that exhibit some of the largest observed increases in activation enthalpy for catalysis relative to WT (66) (*SI Appendix*, Table S1). The full set of eight SLO variants interrogated display E_a_(*k*_cat_) values from *ca*. 0 to 10 kcal/mol, providing a broad range of catalytic properties (*SI Appendix*, Table S1) from which to test the correspondence between the magnitude of activation energies for Stokes shifts decay rates and for the rates of catalysis.

Each of the above-described variants was mutated at Gln322 to cysteine and reacted with the thiol-reactive alkyl bromide group of BADAN (BD) according to protocols established with WT SLO (61) (Fig. 3*A*). The resulting bioconjugates were characterized by electrospray ionization mass spectrometry (ESI-MS) to confirm 1:1 adduct formation (*SI Appendix*, Table S4). To verify the fidelity of the probe attachment to Q322C, liquid chromatography-tandem mass spectrometry (LC-MS/MS) was performed on the pepsin digests of each SLO-BD conjugates (*SI Appendix*, Fig. S6), corroborating sole modification at position 322C, with a mass addition (+211 Da) corresponding to the 6-acetyl-2-dimethylaminonaphthalene group. In all instances, the Cys mutation and attachment of BADAN at Gln322 position yields proteins with almost identical kinetic parameters (*k*_cat_, E_a_) to the parent SLO (*SI Appendix*, Table S5), making these constructs well-suited for probing protein dynamics.

Time-resolved emission spectra (TRES) were constructed from the steady state (*SI Appendix*, Fig. S8) and time-resolved fluorescence decay data upon excitation of BADAN at 373 nm (See Materials and Methods). The time-dependent Stokes shift is attributed to solvent/environmental reorganization around the excited state probe, and is manifest as continuous red shifts in emission spectra until the steady state emission wavelength is approximated (67-69). The TRES for the Q322C-BD labeled SLO mutants at 30 °C are shown in fig. S8, with total Stokes shifts varying from ∼700-1800 cm^-1^ (*SI Appendix*, Table S6). The Stokes shift decay curves (*SI Appendix*, Fig. S9) were fit to a biexponential function, indicating a fast (τ_1_ ∼ 0.5 ns, ∼60%) and a slower (τ_2_ ∼ 5-8 ns, ∼40%) component (*SI Appendix*, Table S6). This bimodal behavior has been shown to be nearly universal and is typically referred to as the bimodality of the reorientational response of biological water (70). The shorter lifetime (τ_1_) is assigned to the unconstrained (free) water in the biological water layer and τ_2_ is ascribed to the nanosecond reorganization of protein associated with constrained (bound) water (70). Nanosecond Stokes shift decay experiments have been reported for a number of other enzyme systems that include the analysis of a fluorescent probe attached to the surface of a protein tunnel in haloalkane dehalogenase (71) and to the active site of glutaminyl-tRNA synthetase (72). Molecular dynamics simulations support the view that water molecules hydrating the protein surface will undergo restricted translational and rotational motions (73, 74).

Arrhenius plots of the Stokes shift decay rates in SLO variants (*SI Appendix*, Fig. S10) reveal distinctive temperature dependences in the slower component (τ_2_), but not the fast component (τ_1_) (Fig. 4 and *SI Appendix*, Fig. S11). In WT SLO, both τ_1_ and τ_2_ were seen to be temperature dependent (61), however temperature independent values for τ_1_ appear to be the more general case. The previously reported viscosity independence of *k*_cat_ in SLO (66, 75) is consistent with a primary role for the hydration shell (related to τ_2_) rather than the bulk solvent (related to τ_1_) in the enzymatic activation barrier (33). The activation energies (E_a_) for τ_2_ in SLO range from ∼0-4 kcal/mol (*SI Appendix*, Table S8) and are similar to reported E_a_ values for the nanosecond dynamics of a solvent exposed tryptophan in a thermophilic ADH (76, 77) and the picosecond hydration dynamics around solvent-exposed tryptophan in Dpo4 (78), ApoMb (79), and SNase (80). The comparison of the experimental activation energies for Stokes shift decay rates (1/−_2_) and for *k*_cat_ as a function of site-specific mutagenesis is presented in Fig. 4. The data indicate a remarkable 1:1 correspondence for WT SLO and four variants within the thermally activated network, I553G, I552A, I553A, and L546A (Fig. 4, red line), generating a line with slope of 1.0(0.1), r^2^ = 0.99. The L754A mutant that lies outside the thermal network (52) (see Fig. 1C, blue data point) acts as an internal control; in this case, the correlation between the temperature dependence of Stokes shifts and catalysis is displaced to the right (Fig. 4, blue line). Note that for the double mutants that contain both on-network and off-network mutations, I553A/L754A and L546A/L754A, the temperature dependence of the Stokes shift decay arises solely from the on-network mutation (I553A, L546A). With 3 data points, this second line of L754A-containing variants displays a slope of 0.7(0.1) and a correlation coefficient of r^2^ = 0.99. The results with L754A mutation indicate that the equivalence of E_a_ values for catalysis and Stokes shift decay is restricted to residues that have been assigned to the thermal network.

**Fig. 4.**
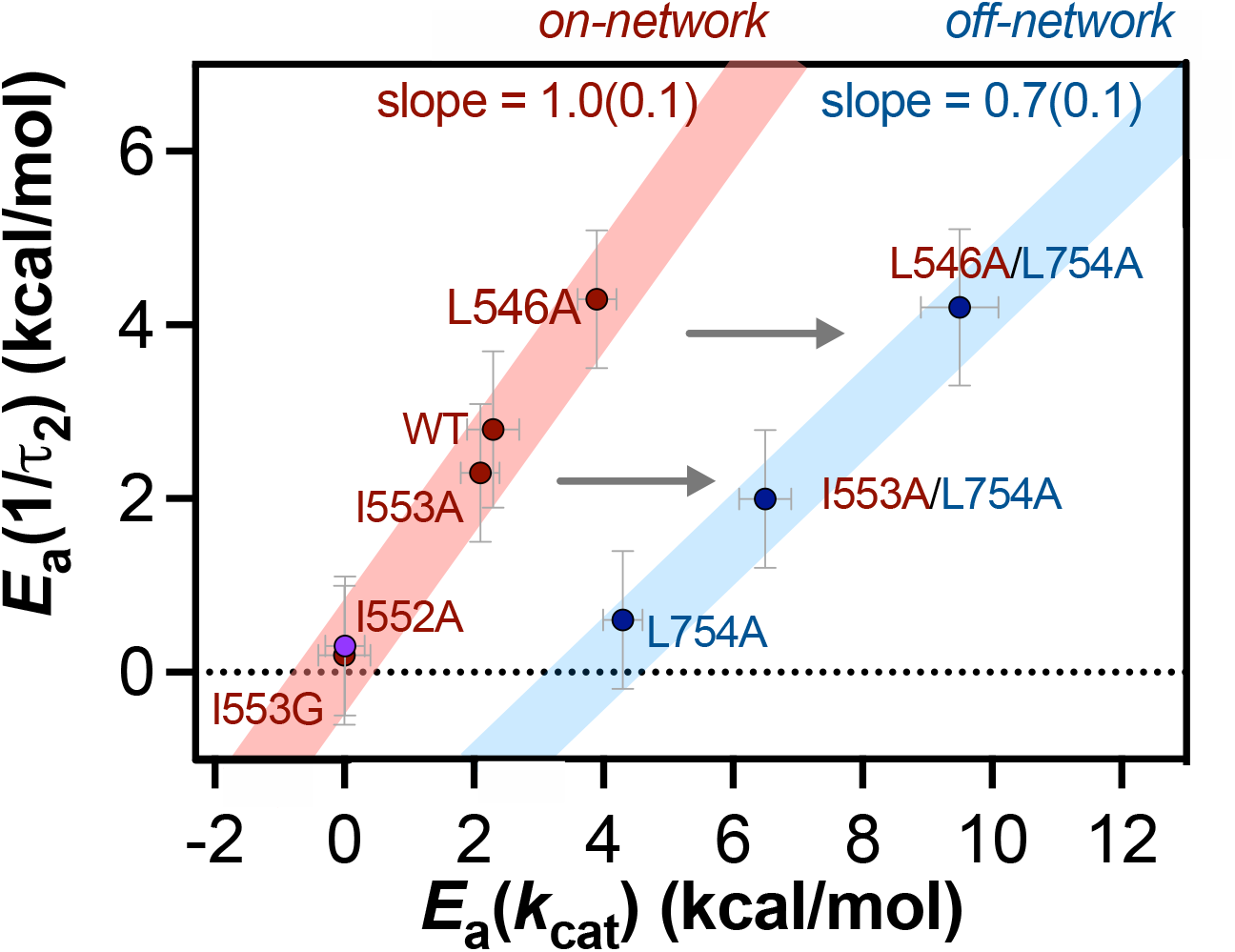
Comparative activation energies for the Stokes shift decay rates 1/τ_2_ (E_a_(1/τ_2_)) and for catalysis (E_a_(*k*_cat_)). The red line indicates residues that are along the thermal network. Given the very close overlap between the datapoints for I552A and I553G, the datapoint for I552A is shown in magenta to allow distinction from I553G. The blue line represents the single mutant L754A together with two double mutants, I553A/L754A and L546A/L754A. In this case, the temperature dependence of the observed Stokes shift decays arises solely from the on-network mutations (I553A or L546A, labeled red).

In light of the unprecedented correspondence between the E_a_ values for Stokes shift decay (τ_2_) and for *k*_*cat*_, we considered it important to rule out unanticipated impacts from thermally induced protein destabilization and possible global unfolding in the SLO samples interrogated herein. Early cryogenic X-ray studies of hydrophobic side chain variants of apo-SLO had indicated virtually no change in protein structure relative to WT (49, 65), while subsequent advanced ENDOR studies of the SLO-substrate complex revealed a progressive elongation of hydrogen donor-acceptor distances as the volume of hydrophobic side chains was reduced (53, 81). The cryogenic X-ray behavior of SLO is consistent with similar studies of other enzyme systems that show little or no change to overall protein structure following a reduction in the size of interior hydrophobic side chains, with an incursion of water molecules into enlarged interior cavities seen only rarely (82-85). Differential scanning calorimetry (DSC) has been used to provide quantitative estimates of the impact of changes in hydrophobic side chain volume on protein unfolding thermodynamics (ΔH°) relative to native enzyme (ΔΔH° = ΔH°(mutant) – ΔH°(WT)) (82, 86, 87). As summarized in *SI Appendix*, Table S9 and Fig. S12, DSC analysis of all 5 SLO variants located within the protein’s thermal network yield values for ΔΔH° in the range of those found for hydrophobic mutations in the prototypic and much smaller enzyme lysozyme (82, 86, 87) and are within experimental error of each other (*SI Appendix*, Table S9). The magnitude of the melting point values is also informative, with little difference seen among the values of WT, L546A and L546A/L754A (DM) together with a decrease by *ca*. 4 °C in the case of I553G and I552A. The high overall T_m_ values of SLO are similar to what was measured for a related isoform (88) consistent with the large size and compact global structure of this protein, and are 20 – 25 °C above the temperature ranges of the kinetic experiments reported herein. Of note, the decreases in T_m_ for I553G and I552A track with the decrease in their E_a_(*k*_cat_) ∼ 0 kcal/mol (*SI Appendix*, Table S1 and Fig. 4). The TDHDX-MS for the variants I553G and I552A indicated enhanced disorder within peptide 317-324 (Fig. 1C), suggesting a thermally induced destabilization within this region of protein. The unusual observation of net experimental E_a_(*k*_cat_) values within error of zero for I553G and I552A may then be a consequence of compensating temperature dependences, whereby the productive thermal activation of tunneling (ΔH^‡^ > 0) has been offset by a need to first restructure disordered ground state configurations (ΔH° < 0). The pattern seen for I553G and I552A in Fig. 4 is notable and in support of site-specific motions within the protein scaffold as the origin of corresponding E_a_ values for Stokes shift decay and catalytic rate constants.

## Discussion

Room temperature X-ray crystallography of proteins has gained attention as a powerful tool in detecting amino acid side chain disorders that can be related to enzyme function (5, 6). In the present study we analyzed RT X-ray structures for single site mutants of SLO with demonstrated impacts on the properties of hydrogen tunneling from C11 of the linoleic acid substrate to the active site Fe^III^(OH) cofactor (19, 52) (Fig. 1*A*). Electron density analyses (*SI Appendix*, Fig. S3 and Fig. 2*A*) extend the thermal transfer pathway identified by TDHDX-MS (19) to an island of primarily hydrophobic side chains that reach from the protein/solvent interface to the active site (Fig. 2*B*). Both methods report on thermally-averaged protein sub-states that undergo functionally relevant responses to either mutation (RT X-ray studies) and/or to changes in temperature (TDHDX-MS). In the case of TDHDX-MS, the linear relationship between E_a_(*k*_HDX_) and E_a_(*k*_cat_) (52) (Fig. 1*C*, red line) provided initial insights into the location and composition of a thermal network for enzyme catalysis in SLO. An important caveat of the TDHDX-MS derived correlation is the significant difference in the magnitudes of E_a_(*k*_HDX_) in relation to E_a_(*k*_cat_), as expected for inherently different processes that reflect local and fully reversible protein unfolding events (E_a_(*k*_HDX_)) vs. the tuning of active site electrostatics and distances that promote hydrogenic wave function overlap between the donor and acceptor atoms (E_a_(*k*_cat_)).

The presented nanosecond Stokes shift analyses differ in a fundamental way from the TDHDX-MS experiments, uncovering activation energies for a photo-induced environmental reorganization around a surface attached probe in SLO that are *identical* to the measured activation energies for cleavage of the C-H bond of substrate (Fig. 4). A limited number of previous studies have shown corresponding enthalpies for distinct kinetic processes within a single system, that include a comparison of Stokes shifts and fluorescence anisotropy using the method of tryptophan scanning in DNA polymerase IV (78) and a comparison of the rate of solvent dielectric relaxation to changes in Mössbauer spectra following the photo-excitation of the carbon monoxide-heme complex in myoglobin (33). However, neither of these studies was able to address the physical origins and site specificity of heat activation in the context of an enzyme catalyzed bond cleavage reaction. The observation of the same enthalpy of activation among variants of SLO for two distinctive chemical processes that take place with rate constants that differ by *ca*. 10^6^-fold and at a distance of at least 30 Å apart, provides compelling evidence for a shared protein quake that connects thermal activation at the protein/solvent interface to the primary bond cleavage step in SLO. The regional specificity of this heat flow, as first revealed from in depth studies of both TDHDX-MS and RT X-ray, illustrates the importance of anisotropic thermal conduits as a means of generating productive activated enzyme-substrate (ES) complexes that also avoid interference from disruptive protein motions that could slow down or eliminate catalysis. While the detection of outward directed protein quakes during the photoexcitation of protein-bound chromophores has been previously described (34), the data presented here implicate a functional and inward-directed protein quake in SLO that reaches from a localized protein surface toward a deeply buried active site.

Discussions of the role of protein dynamics in enzyme catalysis have been primarily focused on the role of conformational landscapes that are comprised of large numbers of protein substates that can be readily accessed at temperatures relevant to catalysis (89-93). One key question from the present work is the inter-relationship between a thermal quake from the solvent bath to an enzyme active site and the distributed conformational landscape. A revised working model of protein dynamics, Fig. 5, begins with the growing body of experimental evidence for protein embedded networks (18-22) that provide pathways for heat activation in thermally initiated enzyme reactions. The principle of conformational selection in protein function, as conceptualized by Frauenfelder for ligand binding dynamics in myoglobin (94), and developed by Kern and co-workers for catalyzed reactions (91, 92) can be readily integrated with this feature, via substrate induced shifts in the conformational ensemble of enzyme-sub-state complexes toward configurations that place the reactive components of an enzyme active site in close proximity to the region of protein capable of efficient heat transmission. The model in Fig. 5 provides a context from which to rationalize the large and growing body of evidence from directed evolution (95-98) and high throughput screening (99) that regions distal to the active site play essential roles in enzyme catalysis (100).

**Fig. 5.**
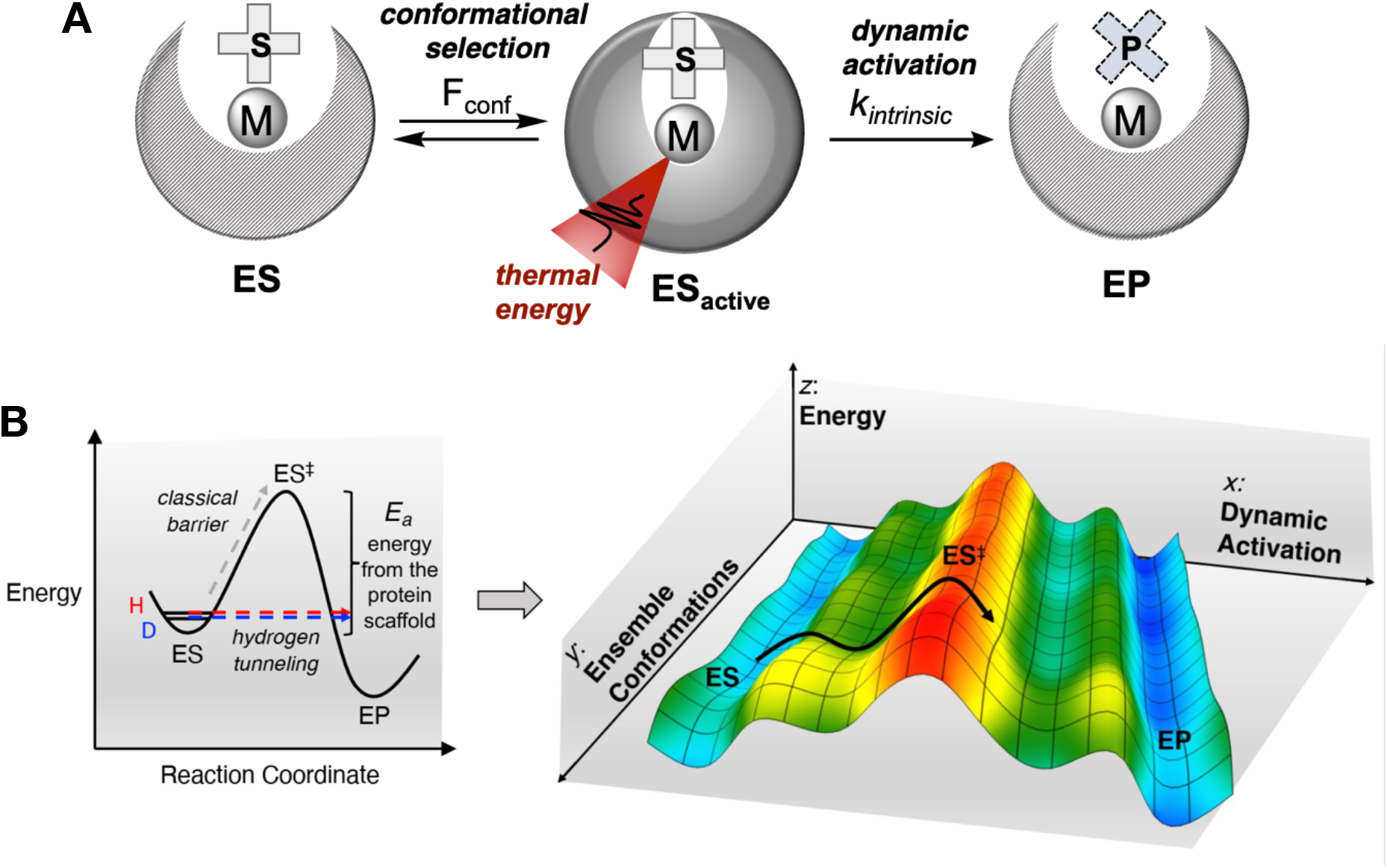
A revised working model of protein dynamics. (*A*) Two distinct types of protein motions contribute to enzyme catalysis. An initially formed ES complex first undergoes conformational selection to achieve refinement in the position of bound substrate relative to active site cofactor and surrounding protein side chains, F_conf_ = [E_active_]/[E_total_]. The resulting family of ES_active_ states is positioned to take maximal advantage of an embedded site-specific network that provides transient activation via thermally activated protein quakes. The term F_conf_ has been introduced into formal rate expressions for non-adiabatic hydrogen tunneling reactions that describe the deep tunneling reaction of SLO, in order to reconcile experimental rate constants that are significantly smaller than the computed tunneling rates (50, 101) (*B*) Left: Two-dimensional reaction coordinate illustrating contribution of protein scaffold dynamics to thermal activation of non-adiabatic hydrogen tunneling. Right: Three-dimensional representation of the free energy landscape for catalysis, adapted from Ref. (1). As illustrated, the initial ES complex (blue) undergoes conformational selection among a large number of possible ground state configurations (green and yellow) that are in rapid equilibrium near RT (y coordinate). Productive barrier crossings (red) occur when the transient alignment of substrate, active site side chains and/or cofactor coincides with a protein embedded thermal conduit (x coordinate).

A key question from this work concerns the different time scales for measured fluorescence Stokes shifts (nanoseconds) vs substrate turnover (milliseconds) that differ by *ca*. 10^6^-fold. Protein motions are expected to vary greatly in rate and to be dependent on the forces and locality of the motions being studied and to the initiating events. With regard to thermal transfer through a protein, XFEL-based studies of heat evolution in a light activated protein illustrate structural changes that radiate out from the active site to solvent on a time scale of picoseconds (34). This time scale is consistent with experimental and computational studies of a number of well-studied model systems, e.g., myoglobin and albumin, that consistently indicate lifetimes in the range of 1 – 50 ps for heat flow in proteins over distances of tens of angstroms (102-107). None of these studies directly address the question of whether rapid, long distance and cooperative heat flow through a protein may deviate from a fully equilibrated process. In a recent study of IR-detected protein motions following photochemical excitation of a surface probe in bovine serum albumin, an out-of-equilibrium protein quake was proposed with properties of a Frochlich-like condensate comprised of coupled low frequency vibrations (108).

A simple but comprehensive formulation for the observation of a range of rate constants that arise from a shared thermal activation of the protein scaffold is shown in equation 1:

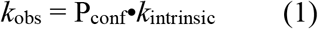

where the measured rate constant *k*_obs_ depends on both the primary thermal quake of the protein scaffold, *k*_intrinsic_, and the probability that the resulting structural rearrangement is sufficient for the process undergoing measurement, according to P_conf_. In the case of a fully optimized enzyme catalyzed reaction, P_conf_ may be very small, a result of the dependence of productive barrier crossings on a multitude of transiently optimized electrostatic interactions and precise internuclear distances. Conformational sampling may play a key role in increasing the magnitude of P_conf_ into a range compatible with the typical millisecond turnover numbers of enzymes, however, in any case, the achievement of the barrier crossing states in enzymes is a comparatively rare event. The magnitude of P_conf_ can reasonably be expected to become much larger in the case of the Stokes shifts behavior of a surface-appended fluorophore, given the availability of a wide range of both protein and solvent interactions for the stabilization of excited state dipoles, as reflected in the observation of Stokes shifts within the picosecond to nanosecond regimes (61, 76-79).

A pictorial representation of Eq (1) is given in Fig. 6, where a red cone is used to represent a shared site for rapid and recurring heat transfer that originates at the protein-solvent interface and terminates at the active site (labeled ‘thermal input’). The magnitude of the observed rate constants resulting from such scaffold activation is represented by the green cone at the top of the figure (labeled ‘productive output’). In a separate enzyme system catalyzing hydride transfer from an alcohol substrate to a nicotinamide cofactor, microsecond FRET between an interior Trp residue and the nicotinamide ring of cofactor was reported to occur with an enthalpy of activation equivalent to that of millisecond catalysis (109), and this behavior has been integrated into Fig. 6 as arising from a value for P_conf_ that is greater than the catalytic hydride transfer, yet smaller than P_conf_ for the protein surface Stokes shifts measurements described herein. To summarize, the origin of the different timescales for each kinetic measurement arises from a common, thermally activated protein quake that is accompanied by a different probability for achieving the set of protein configurations capable of supporting the physical or chemical event under measurement, analogous to the entropic component of the familiar rate expressions describing chemical processes (110, 111). A similar formalism has been suggested in the context of solvent-slaved motions that modulate the internal movement of carbon monoxide within the interior of myoglobin (112).

**Fig. 6.**
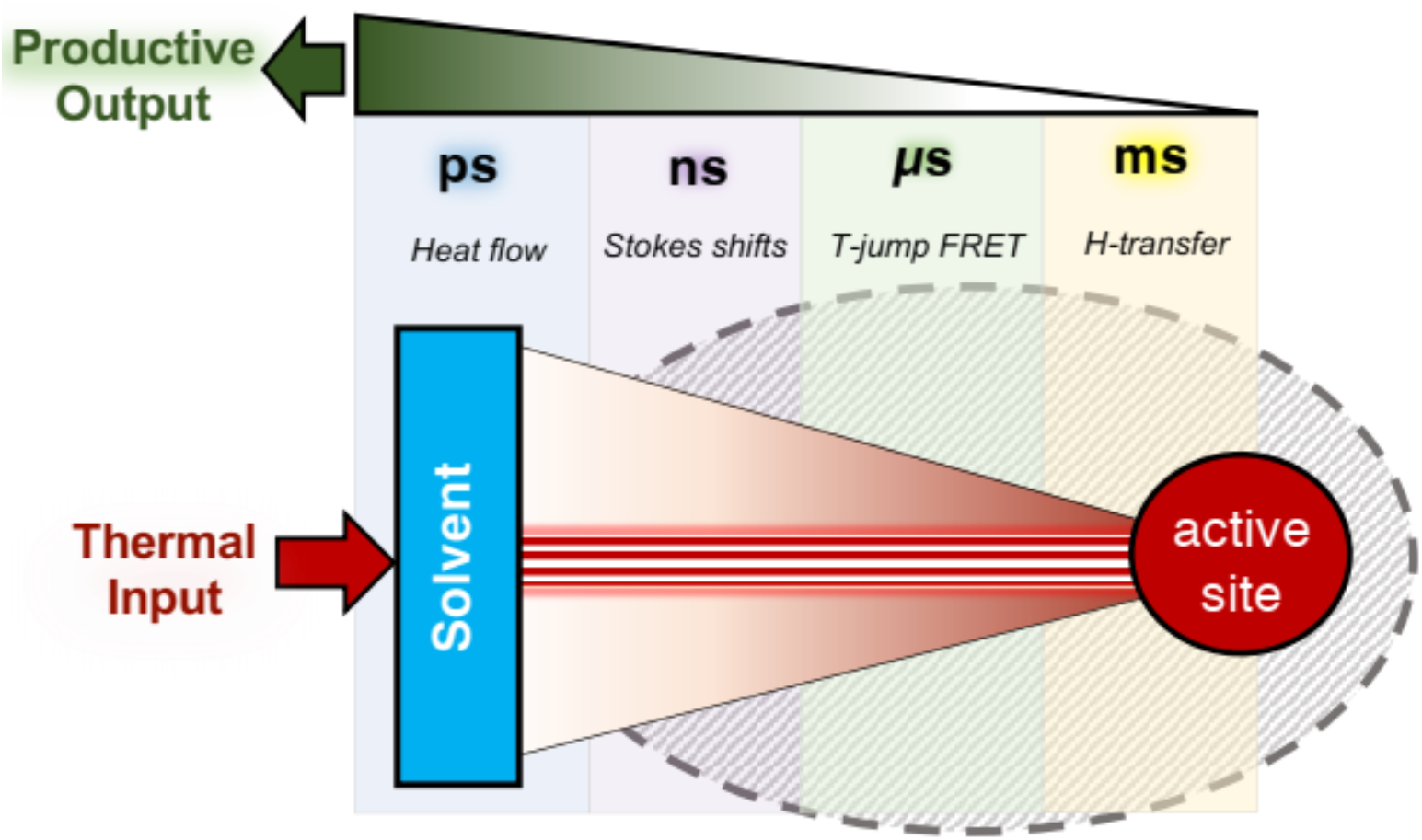
Time scales for productive responses to a long range protein quake. A model is presented that illustrates how different physical events with widely different rate constants can share a common activation energy. According to Eq (1) and the discussion in the text, a shared protein quake leads to a site-specific transfer of thermal activation from the protein/solvent interface to the active site on a picosecond time scale (indicated by the red cone in the above figure). The response (green cone, labeled productive output) is seen to vary from nanoseconds for Stokes shifts in SLO (present study) to microseconds for FRET behavior in the case of an alcohol dehydrogenase (109); for both processes, the frame of reference is enzymatic turnover on a millisecond time scale.

There has been a recent resurgence and growing interest in understanding and harnessing the role of quantum mechanics, including spin coherence, entanglement and tunneling, in biological and other macroscopic processes (113, 114). Among the unresolved questions is the quest to determine and quantify the link, and possible synergy, between classical and quantum behavior, and the degree to which noise at the warm temperatures of biology influences the efficiency of quantum effects. The defining feature from this work is the primacy of the extended protein scaffold in overcoming the energetic barriers of tunneling; this necessitates a simultaneous reduction in the donor-acceptor distance (DAD) and the creation of transient degeneracy between reactant and product states (42, 44). An atomistic understanding of how classical motions within the scaffold of SLO may control these quantum properties becomes apparent when the substrate linoleic acid is modeled into the available X-ray structure, Fig. 7A. The fluctuations in this localized region of aliphatic residues are influenced by the remote, solvent-exposed loop. These motions culminate at Leu546, a key residue in direct contact with the reactive carbon of substrate (C11). Thermally activated changes in this region are expected to moderate the distance between the substrate hydrogen donor and its acceptor, the iron bound hydroxide ion, leading to a transient reduction in DAD from an initial van der Waals interaction (Fig. 7B and Ref. (53)) to a tunneling ready distance of *ca*. 2.7 Å (42, 44, 115). Thermally activated motions that alter the substrate cofactor distance may reasonably be expected to propagate along the substrate chain, producing a concomitant reduction in distance between a remote hydrophobic carbon of substrate (C14) and Ile839. The residue Ile839 is directly ligated to the active site iron of SLO and has been previously proposed to facilitate the tunneling process by moderating hydrogen bonding interactions between the iron bound hydroxide ion and its C-terminal carboxylate (50, 116). Accordingly, a shared protein quake becomes capable of both a reduction in the H-donor-acceptor distance and an alteration of local electrostatics that enables transient production of matched energy states for reactant and product.

**Fig. 7.**
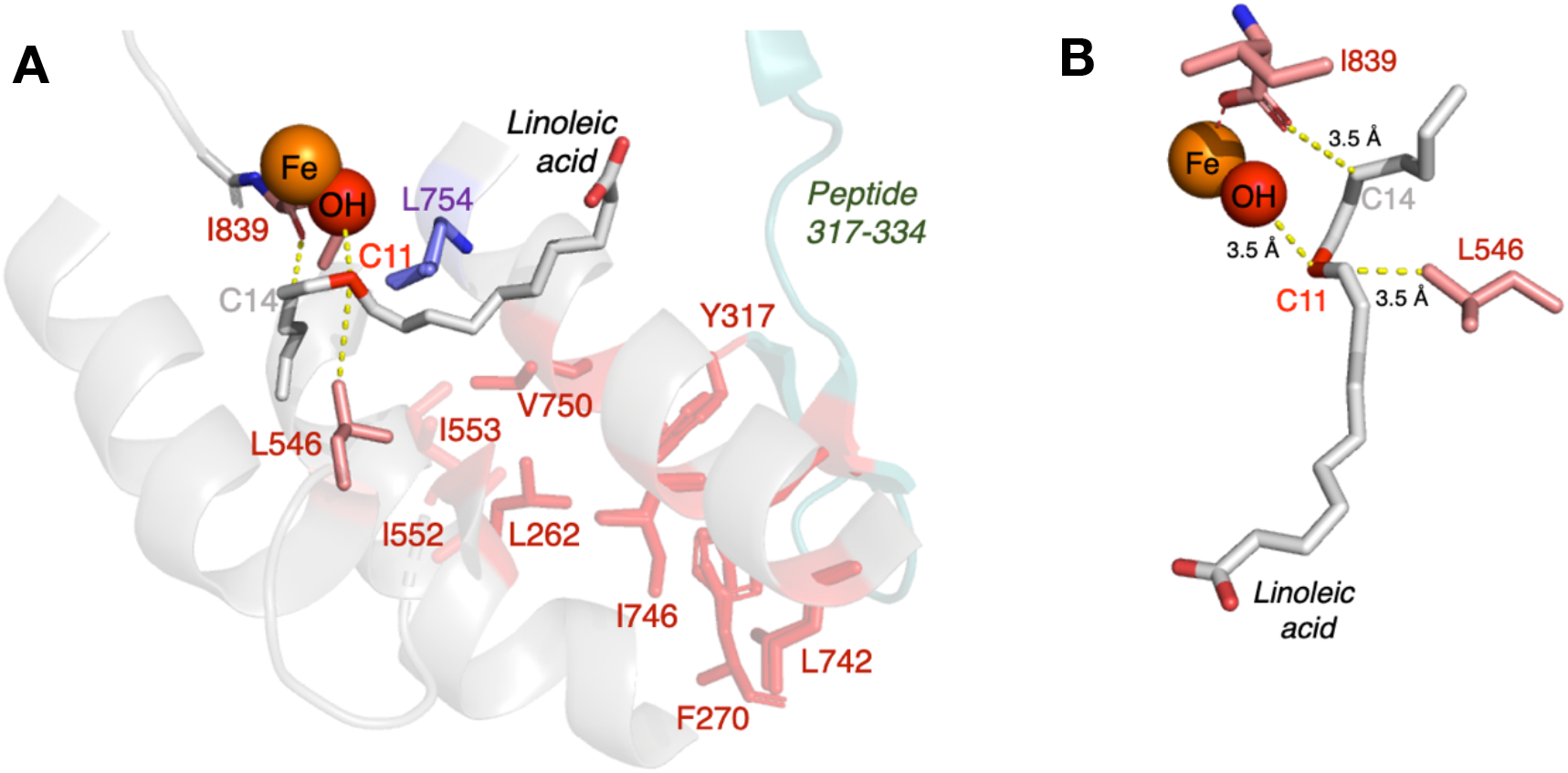
Graphical representation of the proposed role of thermally-activated protein scaffold motions in modulating the donor-acceptor distance (DAD) and reactant/product energetics in SLO catalyzed hydrogen tunneling. (*A*) The bound linoleic acid substrate (gray) is modeled from previous QM/MM calculations (51) and overlaid in the WT RT X-ray structure (PDB: 5T5V). The experimentally identified solvent-exposed loop (peptide 317-334) is shown in blue-green. Connecting this loop to the active site is a network of mainly aliphatic residues (in red) that mediates the heat transfer, and culminates at Leu546 in van der Waals contact with the reactive carbon C11. (*B*) Interactions between LA and the Fe(OH) cofactor are dependent on fluctuations of L546 and I839 that arise from within the defined thermal network.. The side chain L546 influences the positioning of the carbon backbone of bound LA, while I839 mediates the electrostatic environment.

The presented model (Fig. 5) differs from previous textbook descriptions of enzyme catalysis that are primarily focused on static enzyme structures and are rooted in the principle of “enhanced transition state binding” (117, 118). Though this particular study is focused on an enzyme reaction that proceeds via a deep tunneling hydrogen transfer (Fig. 5B, left) a combination of distributed protein conformational substates and embedded thermal networks within a protein scaffold is expected to play a key physical role in enzymatic rate enhancements, independent of the reaction catalyzed. The dominant role of the protein scaffold can be visualized through a three-dimensional energy plot (Fig. 5B, right). As illustrated, the primary coordinates for catalysis are an x-coordinate that represents site specific distribution of heat into the protein scaffold to reach the high energy states capable of barrier crossings (ES^‡^, labeled red) that acts in concert with an orthogonal distributed conformational landscape (y coordinate). The latter provides the high specificity with which a small sub-set of the conformational landscape can respond to local anisotropic heat transfer. This model incorporates a functionally linked thermal activation of embedded protein modes into the contributions of protein conformational selection induced by binding of either substrate (in the case of enzyme catalysis) or allosteric effector (in the case of enzyme regulation). The newly presented data and interpretation both emphasize the unique properties of enzyme reactivity relative to the reaction of small molecules in condensed phase and serve as a launching point to answering the long-sought goal of integrating the protein matrix into our understanding of the origins of naturally evolved enzymatic rate accelerations and our ability to carry over this knowledge into rational *de novo* design.

## Materials and Methods

### General

All reagents were purchased from commercial sources at the highest grade available. Water was purified to a resistivity of 18.2 MΩ·cm (at 25 °C) using a Milli-Q Gradient ultrapure water purification system (Millipore, Billerica, MA). DNA sequencing was performed at the UC Berkeley DNA Sequencing Facility. Mass spectrometric analysis of intact and pepsin-digested proteins was performed at the UC Berkeley QB3/Chemistry Mass Spectrometry Facility.

### Site-Directed Mutagenesis of SLO-1

The SLO-1 mutants I552A and V750A were generated as previously described (56). In addition to these mutants, site-directed mutation of the tip of the dynamic loop (Q322) to cysteine (Q322C) was performed following the Q5® site-directed mutagenesis protocol starting from SLO-1 genes (WT, I553G, I553A, I552A, L546A, L754A, I553A/L754A, L546A/L754A) contained in a pT7-7 *E. coli* expression vector(119). The forward and reverse primers were designed using NEBaseChanger®, and obtained from Eurofins Genomics (Louisville, KY). The mutant plasmids were generated and amplified using a PTC-2000 Peltier Thermal Cycler (MJ Research). Plasmid purification for gene sequencing was performed using the QIAprep spin Miniprep kit (QIAGEN). Mutations were confirmed by sequencing with three different primers that targeted the beginning, middle, and end regions of the gene.

### Protein Expression and Purification

SLO-1 was expressed using the pT7-7 vector in *E. coli* BL21(DE3) Codon Plus RIL cells (Agilent) as described previously with minor modifications (119). Purification of the harvested cell pellets proceeded following a previously described procedure(61). For SLO-1 variants bearing the surface cysteine Q322C mutation, the purification procedure was performed in the presence of 0.1 mM TCEP to avoid cysteine oxidation. The purified SLO-1 was buffer-exchanged in either 0.1 M pH 9.0 borate buffer, or 0.5 mM TCEP, 50 mM pH 7.0 HEPES buffer for the Q322C variants. The purity of the enzymes was greater than 95% as assessed by SDS-PAGE. Protein aliquots were stored at -80°C. The iron content for SLO-1 variants was determined using ferrozine assay(120), and typically found to be in the range of 0.8 ± 0.1 Fe atoms per SLO-1 molecule.

### Labeling of Surface Cys-modified SLO with BADAN

Prior to labeling, TCEP was removed from solution by buffer exchange with deoxygenated 50 mM pH 7.0 HEPES. A solution of SLO (200 µM) was incubated in the presence of 5-fold excess of BADAN (in DMSO) at 23 °C in 50 mM HEPES (pH 7.0) for 3 h. The volume of DMSO was kept to <10% of the total buffer volume. Excess unreacted BADAN was removed by gel filtration by passing the reaction solution through a PD-10 column (GE Healthcare). Labeling efficiency was measured by UV-visible spectroscopy based on extinction coefficients of SLO-1 (ε_280_ = 132 mM^-1^ cm^-1^) (19) and BADAN (ε_387_ = 21,000 M^-1^ cm^-1^)(121). BADAN labeling efficiency is typically found in the range of 0.9 ± 0.1 BADAN per SLO-1 molecule. BADAN-labeled SLO-1 was stored at 50-150 µM, 50 µL aliquots at -80 °C until further use.

### Characterization of BADAN-SLO conjugates using Mass Spectrometry

BADAN-labeled SLO was analyzed by mass spectrometry to confirm site-specific cysteine bioconjugation with BADAN. Intact Mass Spectrometry was performed for the full protein to confirm BADAN conjugation, and LC-MS was performed on pepsin-digested SLO to obtain sequence coverage maps confirming site-specificity of BADAN conjugation at Q322C. A summary of all mass spectrometry data, including the sequence coverage maps obtained from the pepsin digests is shown in Table S4 and Figure S6.

### Enzyme Kinetics

Steady-state kinetics for both unlabeled and BD-labeled Q322C SLO variants was performed on a Cary 50 UV/vis spectrophotometer in the single-wavelength mode (λ = 234 nm). The reaction progress was monitored by following the production of 13-(S)-hydroperoxy-9,11-(Z,E)-octadecadienoic acid [13-(S)-HPOD] (ε_234_ = 23,600 M^-1^ cm^-1^). All assays were performed in 100 mM borate (pH 9.0) under an ambient atmosphere at 10 – 40°C, regulated by a water-jacketed cuvette holder attached to a NESLAB RTE-111 circulating water bath (temperature stability is ± 0.1 °C). The substrate concentrations for the kinetics vary in the range of 1-220 μM, based on the *K*_M_ values at different temperatures. The kinetic parameters *k*_cat_ and *K*_M_ were determined from the Michaelis-Menten non-linear fits in GraphPad Prism v8.4.2 and represent the mean ± standard error from the mean of three independent measurements for each protein variant. All initial rates were normalized to the fraction of bound iron in the enzyme samples. The ^D^*k*_cat_ was determined as the ratio of the first-order rate constant for protium (for H-LA) versus deuterium (for d_31_-LA) transfer (i.e. ^D^*k*_cat_ = *k*_cat_(H)/*k*_cat_(D)). The perdeuterated linoleic acid (d_31_-LA) was isolated and purified as previously described (56). The activation energies for protio substrate, E_a_(H), were determined by linear fits to Arrhenius plots and the data represent the mean ± s.e.m. from at least seven temperatures for each variant reported. The ΔE_a_ was determined from the difference in activation energies for protio substrate (Ea(H)), and deuterated substrate (Ea(D)). In general, the *k*_cat_, E_a_ and ΔE_a_ parameters for the Q322C-BD labeled samples are similar to the unlabeled (Q322) samples within experimental error. A summary of all enzyme kinetics data (*k*_cat_ at 30 °C and *E*_a_) is shown in Table 1, Table S1, and Table S5.

### I552A SLO HDX Sample Preparation

HDX samples of I552A SLO-1 were exchanged as a function of time by a 10-fold dilution into D_2_O (99% D; Cambridge Isotope) buffer: 10 mM HEPES pD 7.4 (corrected pD; pD = pHread + 0.4), as previously described (19, 52). The samples were maintained at a desired temperature using a water bath (temperatures studied: 10, 20, 25, 30, and 40°C). At a specified time (10 s, 30 s, 45 s, 1 min, 3 min, 10 min, 20 min, 30 min, 45 min, 1 h, 2 h, 3 h, an 4 h), samples were rapidly cooled (−20 °C bath for 5-6 s) and acid quenched (to pH 2.4 with 90 mM (final concentration) sodium citrate buffer) and immediately digested at 4°C (off-line) using immobilized pepsin (2.5 min). Prior to pepsin digestion, 0.5 M (final concentration) solution of guanidinium-HCl at pH 2.4 was added to help increase sequence coverage during cleavage. Pepsin was removed by brief (8 s) centrifugation and the samples were immediately frozen with liquid N_2_ and stored at −80 °C until data collection. The conditions for these experiments are summarized in Table S10.

### Liquid Chromatography-Mass Spectrometry and Analysis of Hydrogen/Deuterium Exchange Measurements

Peptide fragments were identified using LC-MS/MS with a LTQ Orbitrap XL (Thermo Scientific). Full-scan mass spectra were obtained in positive mode (m/z = 350 to 1800) with a mass resolution setting of 60,000 (at m/z = 400). Data acquisition was controlled by Xcalibur software (version 2.0.7, Thermo). Raw data were searched against the amino acid sequence of full-length SLO-1 using Proteome Discoverer software (version 1.3, SEQUEST, Thermo) for peptide identity from MS/MS spectra. More details of the LC and MS system as well as data collection can be found in Ref. (19). Deuterated, pepsin-digested samples of SLO-1 from HDX experiments were analyzed using an Agilent 1200 LC (Santa Clara, CA) that was connected in-line with the LTQ Orbitrap XL mass spectrometer (Thermo Scientific). Data acquisition was controlled using Xcalibur software (version 2.0.7, Thermo). Mass spectral data acquired for HDX measurements were analyzed using the software, HDX WorkBench(122). The percent deuterium incorporation for each peptide has been normalized for 100% D_2_O and corrected for peptide-specific back-exchange, as previously described(19, 52) (see Table S11). Energies of activation representing the temperature dependence of rates of deuterium incorporated into individual peptides were calculated as previously described(19) and are summarized in Table S12.

### Differential Scanning Calorimetry (DSC)

DSC experiments were conducted on a TA-instruments Nano-DSC microcalorimeter. The SLO samples were prepared at 25-30 μM concentration in 50 mM borate (pH 9). The DSC experiments were carried out in heat-only mode, from 30 – 90 °C at a rate of 1 °C min^-1^. The pressure was kept at 3 atm during the course of the experiment. These experiments were performed in triplicate for WT and duplicate for other SLO variants.

### Steady-State Fluorescence

Fluorescence emission spectra of the BADAN-labeled mutants were collected on a custom-built Fluorolog-3 spectrofluorometer (Horiba Jobin-Yvon). Excitation was achieved with a 450 W Xenon lamp. The light was focused using a double Czerny-Turner excitation monochromator (1 nm bandpass) with 1200 grooves/mm blazed at 330 nm. Photons from sample emission were focused using a single Czerny-Turner monochromator (10 nm bandpass) with 1200 grooves/mm blazed at 500 nm. The excitation and emission optics were calibrated using the lamp spectral maximum at 467 nm and the water Raman scattering band at 397 nm, respectively, using HPLC-grade water in a quartz cuvette.

For BADAN-labeled SLO emission spectra, the excitation wavelength was set at 373 nm, and the emission spectra were collected in 0.1 M pH 9.0 borate buffer, from wavelengths spanning 400-650 nm at 0.5 nm increments. Spectra were collected over temperatures ranging from 10 to 40 °C at 5 °C intervals with a water bath (NESLAB RTE-111) controlling the cell temperature. Samples were equilibrated for 5 min at each respective temperature in a quartz cuvette (Starna Cells) before collecting emission spectra. Peak emission wavelengths were determined by fitting the corrected spectra to a Gaussian curve using GraphPad Prism v8.4.2.

### Picosecond-Resolved Fluorescence Spectroscopy

The time-correlated single photon counting technique (TCSPC) was used to obtain fluorescence decays. Selective excitation of BADAN in each mutant was achieved using a Nano-LED N370 source (λ_exc_ = 373 nm) having a typical full-width at half-maximum pulse of <1.2 ns with a 1-MHz repetition rate. The Nano-LED was powered by a FluoroHub photon-counting controller. Single-photon signals were detected by a TBX-04 photomultiplier tube detection module. The photon-counting hub window contained 1024 channels with a time-to-amplitude conversion range of 56 ps/channel. Lifetimes were measured using reverse mode counting with a 75-ns coaxial delay and 0 ns sync delay. Magic angle (55°) conditions were employed to eliminate lifetime distortions resulting from rotational motion. Fluorescence decays of the BADAN-labeled SLO variants (0.1 µM) were obtained from 450-520 nm in 10 nm intervals with a 10 nm emission bandpass. Instrument response functions were collected once at each temperature using only 0.1 M borate (pH 9.0), with the emission monochromator set at 373 nm. All optical settings and delays were kept constant when acquiring the instrument response function. All decays were collected with a peak preset of 10,000 counts. The time resolution of the instrument was calculated to be ∼120 ps. Procedures for sample incubation during data acquisition were performed in the same manner as previously described and executed at the temperatures ranging from 10 to 40 °C in 5 °C intervals(61).

### Construction of Time-Dependent Stokes Shift Spectra

Time-resolved emission spectra (TRES) were constructed from fluorescence decay data by methods previously described in more thorough detail(67-69). Briefly, the function describing the fluorescence decay was determined by deconvoluting the instrument response function (IRF) from a sum of exponential decays at each respective wavelength, λ using Decay Analysis Software v6.8 (DAS6) as represented by equation 2:

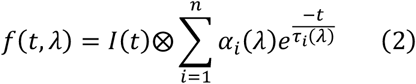

where I(*t*) is the IRF, *n* represents the number of exponential components needed for fitting the decay data, *α*_*i*_ represents the weighted amplitudes such that ∑_i_α_i_(λ) = 1, and τ_i_ represents the fluorescence decay constant for the *i*th component. One exponential was always used to begin fitting the decays, and additional exponentials were iteratively added until a reduced X^2^ < 1.25 was achieved for the residuals. After obtaining satisfactory fitting parameters, an H(λ) that is linearly proportional to the steady-state emission spectrum is calculated using the steady-state emission fluorescence intensity, F(λ) at a particular temperature (equation 3):

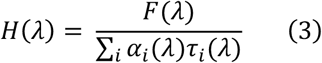

Equation 4 represents the points Γ(λ,*t*) that comprise the TRES, which are calculated at arbitrary time points *t* (0 to 10 ns, in 0.01 increments) by multiplying H(λ) by the deconvoluted decay parameters extracted from the raw data (τ_*i*_, α_*i*_, at a certain λ) represented by equation 2.

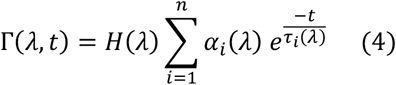

Equations 4 and 5 are used to fit the experimental time points at arbitrary times (*t*) to a log-normal line shape(123, 124) as a function of frequency (ν) in cm^-1^:

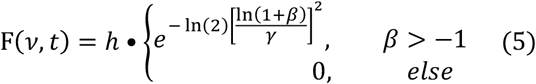

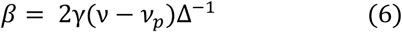

where ν_p_ is peak frequency that is time-dependent, *h* and Δ represent the respective peak heights and width, and γ is the asymmetry parameter. These parameters were allowed to float freely until a best fit was obtained. Fitting of Γ(λ,t) vs ν to a log-normal line shape was performed from *t* = 0 to 10 ns in 0.01 increments (1001 datasets) using the software MATLAB. At a time point *t*, a single value of ν_p_, *h*, Δ, and γ was obtained. For illustration purposes, the intensity Γ(λ,*t*) of the time-resolved emission spectra (TRES) was normalized by dividing Γ(λ,*t*) by the calculated peak height *h* at time *t* from the log-normal line fits.

### Analysis of Stokes Shift Decay Rates

The Stokes shift decay rates were calculated by plotting the calculated peak frequency ν_p_ as a function of time *t*. The data for BADAN-labeled SLO generally follow a bi-exponential decay behavior. The values for the frequencies at time 0 (ν_0_) and time ∞ (ν_∞_), the decay lifetimes τ_*i*_, and the relative amplitudes α_*i*_ were obtained from fitting the data to equation 7.

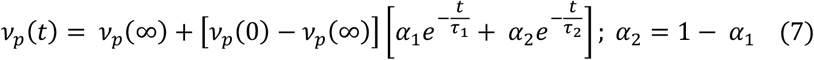

The Stokes shift was calculated as the difference between ν(∞) and ν(0). As the instrument resolution is limited (∼120 ps), only solvation events that are longer than this time are accurately detected. For illustration purposes, the solvation correlation function C(*t*) (eq. 8) was plotted vs. time *t*. As shown in equation 8, C(*t*) is related to ν_p_(*t*) where ν_p_(*t*) is normalized with respect to the total Stokes shift.

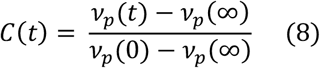

### Analysis of Temperature Dependence of Fluorescence Relaxation Lifetimes and Stokes Shift Decay Rates

The Stokes shift decay rates (τ) (in seconds) was related to temperature T (in K) using the Arrhenius equation (eq. 9). The activation enthalpy (*E*_a_) was calculated and is related to the *E*_a_ values from active site hydrogen transfer rates.

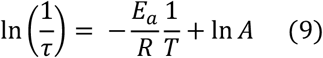

### Room-Temperature Crystallography

An additional protein purification step was performed prior to crystallization of I552A and V750A SLO-1, as previously described(49, 65). Just prior to data collection, crystals were transferred to a solution of 30 % PEG-3350, 100 mM sodium malonate, and 100 mM sodium acetate, pH 5.5(19). Diffraction data were collected at room temperature at Beamline 8.3.1 of the Advanced Light Source at Lawrence Berkeley National Laboratory. Data were processed by the ELVES program(125), with integration performed in MOSFLM(126) and scaling and merging in SCALA, POINTLESS and TRUNCATE(127, 128) or using XDS(129) with scaling and merging using AIMLESS(130), POINTLESS and TRUNCATE in the CCP4 suite(131) provided through SBGRID(132). An initial solution was found by molecular replacement using Phaser(133) with PDB 3PZW as the search model. Manual refinement was performed in Coot(134) and automated refinement using the PHENIX suite(135). Alternate conformers were modeled using qFit(57, 136). We manually deleted waters that did not fit into the F_o_-F_c_ map density and edited alternate side chain conformations as indicated in the positive density of the F_o_-F_c_ map. See Table S3 for data processing and refinement statistics.

### Molecular Dynamics Simulation of Q322C-BD

The initial structure was taken from the RCSB Protein Data Bank (PDB: 3PZW). Where alternate amino acid orientations were available, the position with the higher occupancy was chosen. Missing heavy atoms were filled in with Prime, and hydrogens were added with Maestro’s Protein Preparation Wizard. Waters beyond 3Å of added heteroatom groups were deleted, as were acetate and ethylene glycol molecules which were artifacts of crystallization. The active site iron is in its ferrous state with a water as a coordination oxygen. The Q322 position was mutated to cysteine, and BADAN was added to the thiol group to generate the labeled mutants. A hydrogen-bond optimization was run to flip waters and polar groups to maximize hydrogen bonding, followed by a restrained minimization for hydrogens only. A further minimization was performed with Macromodel. A PRCG algorithm was performed for 500 iterations. An OPLS4 force field was used in water, and the convergence threshold was 0.05. A model system was built using the energy-minimized structure, using the SPC solvent model and an orthorhombic box shape with Buffer and distances of 0 Å. After relaxation of the model system using Maestro’s default protocol, a molecular dynamics simulation was run for 100 ns with a recording interval of 100 ps and ensemble class NPT. The distance from the center of the naphthalene ring in BADAN to the center of the indole ring in Trp340 was measured and observed over the course of the simulation.

## Acknowledgments

We thank Prof. Tom Spiro for the helpful and stimulating discussions. We also thank the members of the Klinman lab for the insightful discussions about the manuscript, specifically Dr. E. Thompson for assistance with programming and data analysis. This research is supported by funding from the National Institutes of Health grants to J.P.K. (GM118117), A.R.O. (GM113432 (F32)), and A.T.I. (1S10OD020062-01). J.P.T.Z. thanks the University of California President’s Postdoctoral Program for a fellowship. Data collection at BL 8.3.1 at the Advanced Light Source is supported by the Director, Office of Science, Office of Basic Energy Sciences, of the U.S. Department of Energy under contract no. DE-AC02-05CH11231, UC Office of the President, Multicampus Research Programs, and Initiatives grant MR 15 32859, the Program Breakthrough Biomedical Research, which is partially funded by the Sandler Foundation, and the NIH P30GM124169.

## Supplementary information

is available for this paper at https://doi.org/

## Data and materials availability

Summary tables of all kinetic and thermodynamic parameters, as well as fluorescence and crystallography data measured for each mutant are included in the supplementary materials. The atomic models have been deposited at the Protein Data Bank with PDB IDs 7SOI (I552A SLO) and 7SOJ (V750A SLO).

## Author contributions

J.P.T.Z., A.R.O. and J.P.K. designed the research. J.P.T.Z., A.R.O., S.H., C.L.G., Z.M.F., N.M., F.F., Z.D., and A.T.I. performed the research. J.P.T.Z., and A.R.O. analyzed data. J.P.K. conceived and supervised the project and acquired funding. J.P.T.Z., A.R.O., and J.P.K. wrote the paper.

