## Supplementary Information for "Temporal and Spatial Resolution of a Protein Quake that Activates Hydrogen Tunneling in Soybean Lipoxygenase"

**This PDF file includes:**

Figures S1 to S12
Tables S1 to S12
SI References

**Other supplementary materials for this manuscript include the following:**

Data S1. HDX-MS deuterium uptake plots for all peptides of I552A and WT SLO
Data S2. Summary of HDX-MS data for I552A SLO at 10, 20, 25, 30, and 40 ˚C

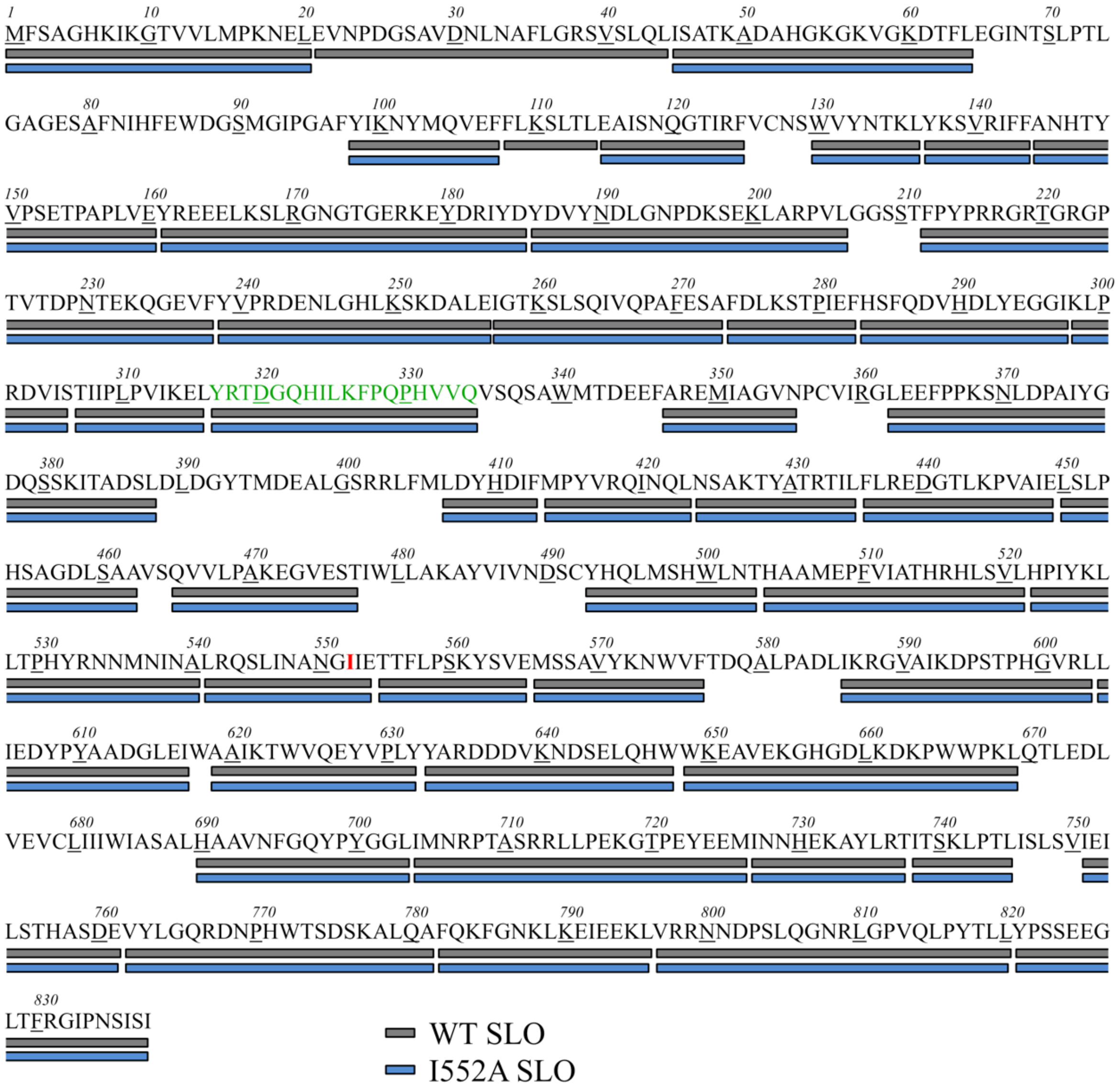


**Fig. S1.** Map of the defined non-overlapping TDHDX-MS peptide analysis of I552A, with the primary sequence of I552A (in blue) vs WT (in gray) SLO. The catalytically non-essential N-terminal β-barrel is represented by residues 1-144. The catalytic domain comprises residues 145-839. To simplify analysis, only the "catalytic" peptides (145-839) were analyzed. The I552 mutation site is in red, and the thermally activated loop (peptide 317-334) is in green.

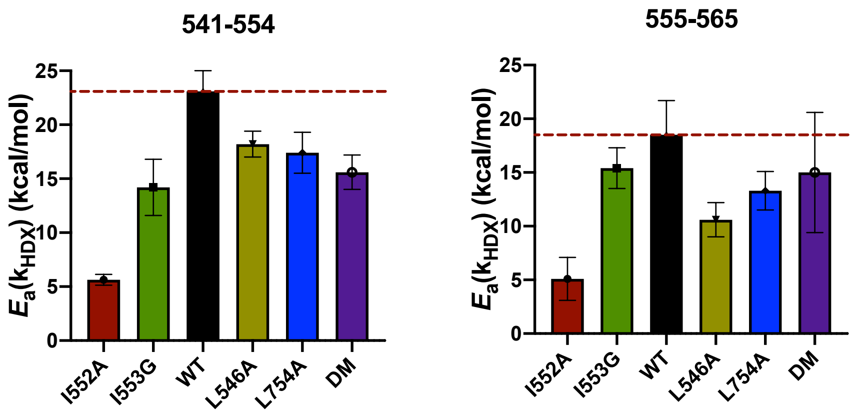


**Fig. S2.** HDX Activation energies ($E_a(k_{HDX})$) of peptide 541-554 (left) and peptide 555-565 (right) from TDHDX-MS of I552A (maroon), in comparison to other SLO mutants[1]. A large decrease in $E_a(k_{HDX})$ is observed for the I552A mutant in comparison to WT and other SLO variants.s

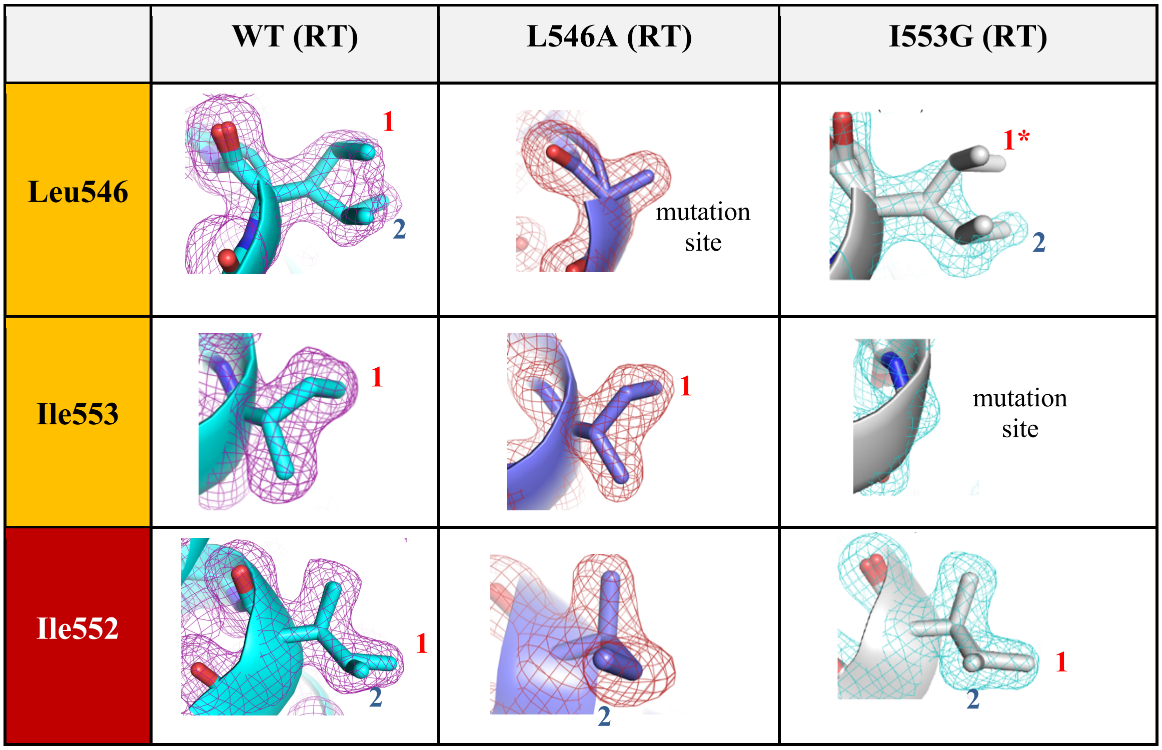


**Fig. S3.** Comparison of electron densities of the thermal network residues in the RT X-ray structures for WT (PDB: 5T5V), I553G (PDB: 5TQP), and L546A (PDB:5TQN) SLO. For I553G, conformer analysis shows the same two conformations of Leu546 as in WT, however a decrease in electron density of conformer 1 was observed.

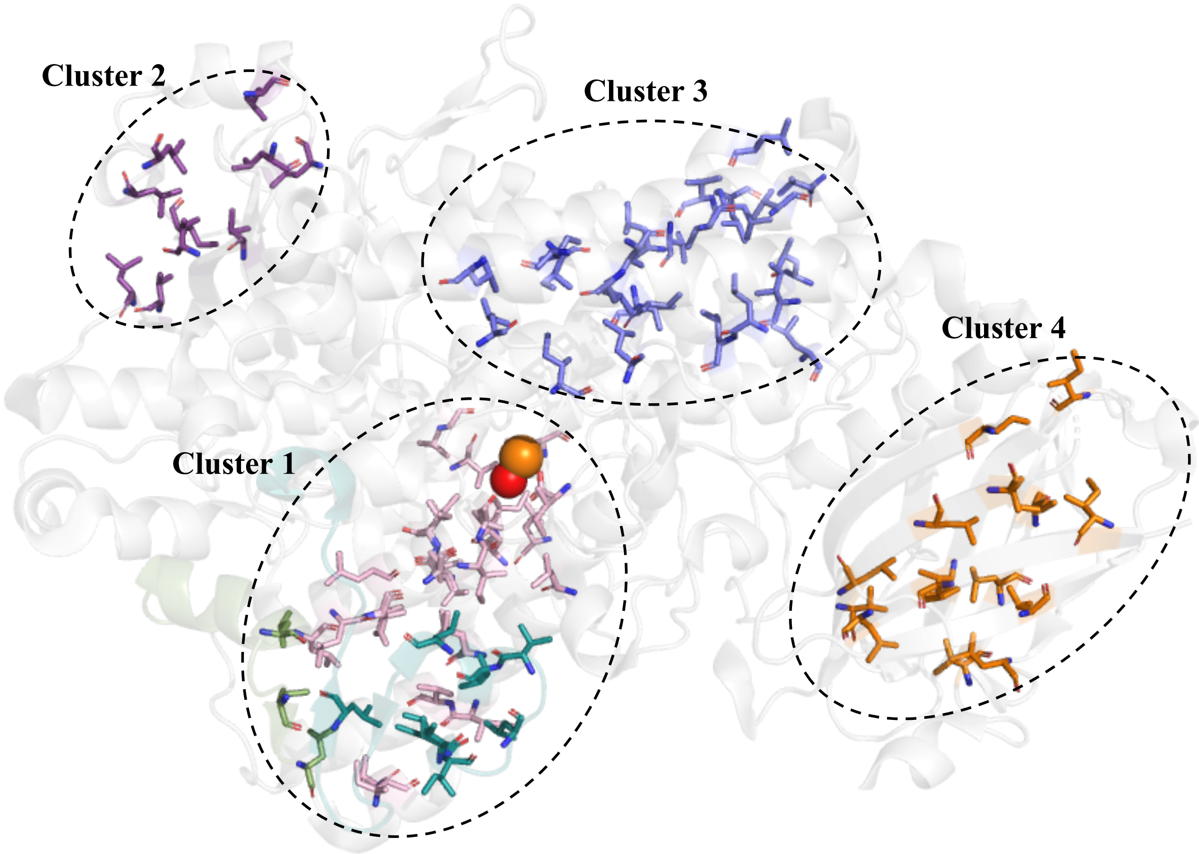


**Fig. S4.** The 4 largest clusters of hydrophobic Ile, Leu, Val (ILV) residues in SLO. Cluster 1, the largest hydrophobic cluster, contains the catalytic Fe center, substrate portal, substrate binding residues I553, L546, L754 subject to mutations, the thermal network of residues, and residues that belong in the thermally activated loop 317-334. Data obtained from ProteinTools[2] (https://proteintools.uni-bayreuth.de/clusters/) using PDB: 3PZW as input.

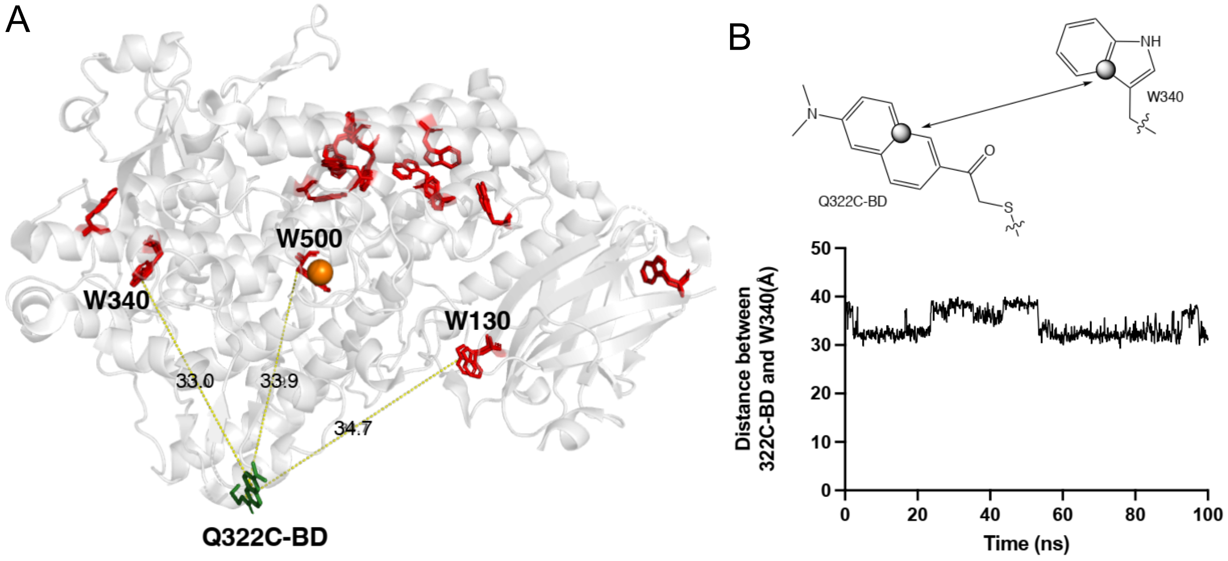


**Fig. S5.** (A) Location of the 14 tryptophan residues (red) in SLO in relation to the Badan probe (green) at the 322 position. The three nearest Trp residues, W340, W500, and W130 are shown. (B) 100 ns molecular dynamics simulation of Q322C-BD showing time-dependent changes in distance (Å) between the Badan ring covalently attached at position 322 and the Trp side chain at Trp340. Reference atoms in both rings are highlighted in gray circles.

**Fig. S6.** Protein sequence coverage maps from tandem mass spectrometry (MS/MS) measurements. For WT SLO data, see Ref. 3. Uppercase amino acids in purple were detected, lowercase amino acids in gray were below limits of detection. Uppercase amino acids in red are hydrophobic mutation sites, and the Badan-modified Q322C mutation is highlighted in green. Protein sequence coverage maps were prepared using the Protein Coverage Summarizer software (Pacific Northwest National Laboratory, https://omics.pnl.gov/).

1. **I553G-Q322C-BD**

   coverage: 774/839 residues, 92.25%

```
MFSAGHKIKGTVVLMPKNELEVNPDGSAVDNLnaFLGRSVSLQLISATKADAHGKGKVGK     60
DTFLEGINTSLPTLGAGESAFNIHFEWDGSMGIPGAFYIKNYMQVEFFLKSLTLEAISNQ     120
GTIRFvcnsWVYNTKLYKSVRIFFANHTYVPSETPAPLVEYREEELKSLRGNGTGERKEY     180
DRIYDYDVYNDLGNPDKSEKLARPVLGGSSTFPYPRRGRTGRGPTVTDPNTEKQGEVFYV     240
PRDENLGHLKSKDALEIGTKSLSQIVQPAFEsaFDLKSTPIEFHSFQDVHDLYEGGIKLP     300
RDVISTIIPLPVIKELYRTDGCHILKFPQPHVVQVSQSAWMTDEEfareMIAGVNPCVIR     360
GLEEFPPKSNLDPAIYGDQSSKITADSLDLdgytmDEALGSRRLFMLDYHDIFMPYVRQI     420
NQLnsaktYATRTILFLREDGTLKPVAIELSLPHSAGDLSAavsqVVLPAKEGVESTiwl     480
lakayVIVNDSCYHQLMSHWLNTHAAMEPFVIATHRHLSVLHPIYKLLTPHYRNNMNINA     540
LARQSLINANGIGETTFLPSKYSVEmssaVYKNWVFTDQALPADLikrgvaIKDPSTPHG     600
VRLLIEDYPYAADGLEIWaaiktwvqeYVPLYYARDDDVKNDSELqhwWKEAVEKGHGDL     660
KDKPWWPKLQTLEDlvevclIIIWIASALHAAVNFGQYPYGGLIMNRPTASRRLLPEKGT     720
PEYEEMINNHEKAYLRTITSKLptlISLSVIEILSTHASDEVYLGQRDNPHWTSDSKALQ     780
AFQKFGNKLKEIEEKLVRRNNDPSLQGNRLGPVQLPYTLLYPSSEEGLTFRGIPNSISI      839
```

**Fig. S6, continued.**

2. **I553A-Q322C-BD**

coverage: 821/839 residues, 97.86%

MFSAGHKIKGTVVLMPKNELEVNPDGSAVDNLNAFLGRSVSLQLISATKADAHGKGKVGK 60

DTFLEGINTSLPTLGAGESAFNIHFEWDGSMGIPGAFYIKNYMQVEFFLKSLTLEAISNQ 120

GTIRFVCNSWVYNTKLYKSVRIFFANHTYVPSETPAPLVEYREEELKSLRGNGTGERKEY 180

DRIYDYDVYNDLGNPDKSEKLARPVLGGSSTFPYPRRGRTGRGPTVTDPNTEKQGEVFYV 240

PRDENLGHLKSKDALEIGTKSLSQIVQPAFESAFDLKSTPIEFHSFQDVHDLYEGGIKLP 300

RDVISTIIPLPVIKELYRTDGCHILKFPQPHVVQVSQSAWMTDEEFAREMIAGVNpcvir 360

gLEEFPPKSNLDPAIYGDQSSKITADSLDLDGYTMDEALGSRRLFMLDYHDIFMPYVRQI 420

NQLNSAKTYATRTILFLREDGTLKPVAIELSLPHSAGDLSAAVSQVVLPAKEGVESTIWL 480

LAKAYVIVNDSCYHQLMSHWLNTHAAMEPFVIATHRHLSVLHPIYKLLTPHYRNNMNINA 540

LARQSLINANGIAETTFLPSKYSVEMSSAVYKNWVFTDQALPADLIKRGVAIKDPSTPHG 600

VRLLIEDYPYAADGLEIWAAIKTWVQEYVPLYYARDDDVKNDSELQHWWKEAVEKGHGDL 660

KDKPWWPKLQTLEDlvevclIIIWIASALHAAVNFGQYPYGGLIMNRPTASRRLLPEKGT 720

PEYEEMINNHEKAYLRTITSKLptlISLSVIEILSTHASDEVYLGQRDNPHWTSDSKALQ 780

AFQKFGNKLKEIEEKLVRRNNDPSLQGNRLGPVQLPYTLLYPSSEEGLTFRGIPNSisi 839

3. **I552A-Q322C-BD**

coverage: 815/839 residues, 97.1%

MFSAGHKIKGTVVLMPKNELEVNPDGSAVDNLNAFLGRSVSLQLISATKADAHGKGKVGK 60

DTFLEGINTSLPTLGAGESAFNIHFEWDGSMGIPGAFYIKNYMQVEFFLKSLTLEAISNQ 120

GTIRFVCNSWVYNTKLYKSVRIFFANHTYVPSETPAPLVEYREEELKSLRGNGTGERKEY 180

DRIYDYDVYNDLGNPDKSEKLARPVLGGSSTFPYPRRGRTGRGPTVTDPNTEKQGEVFYV 240

PRDENLGHLKSKDALEIGTKSLSQIVQPAFESAFDLKSTPIEFHSFQDVHDLYEGGIKLP 300

RDVISTIIPLPVIKELYRTDGCHILKFPQPHVVQVSQSAWMTDEEFAREMIAGVNpcvir 360

gLEEFPPKSNLDPAIYGDQSSKITADSLDLDGYTMDEALGSRRLFMLDYHDIFMPYVRQI 420

NQLNSAKTYATRTILFLREDGTLKPVAIELSLPHSAGDLSAAVSQVVLPAKEGVESTIWL 480

LAKAYVIVNDSCYHQLMSHWLNTHAAMEPFVIATHRHLSVLHPIYKLLTPHYRNNMNINA 540

LARQSLINANGAIETTFLPSKYSVEMSSAVYKNWVFTDQALPADLikrgvaIKDPSTPHG 600

VRLLIEDYPYAADGLEIWAAIKTWVQEYVPLYYARDDDVKNDSELQHWWKEAVEKGHGDL 660

KDKPWWPKLQTLEDlvevclIIIWIASALHAAVNFGQYPYGGLIMNRPTASRRLLPEKGT 720

PEYEEMINNHEKAYLRTITSKLptlISLSVIEILSTHASDEVYLGQRDNPHWTSDSKALQ 780

AFQKFGNKLKEIEEKLVRRNNDPSLQGNRLGPVQLPYTLLYPSSEEGLTFRGIPNSisi 839

**Fig. S6, continued.**

4. **L546A-Q322C-BD**
   coverage: 785/839 residues, 93.56%

```
mFSAGHKIKGTVVLMPKNELEVNPDGSAVDNLnaFLGRSVSLQLISATKADAHGKGKVGK    60
DTFLEGINTSLPTLGAGESAFnihfEWDGSMGIPGAFYIKNYMQVEFFLKSLTLEAISNQ   120
GTIRFvcnsWVYNTKLYKSVRIFFANHTYVPSETPAPLVEYREEELKSLRGNGTGERKEY   180
DRIYDYDVYNDLGNPDKSEKLARPVLGGSSTFPYPRRGRTGRGPTVTDPNTEKQGEVFYV   240
PRDENLGHLKSKDALEIGTKSLSQIVQPAFESAFDLKSTPIEFHSFQDVHDLYEGGIKLP   300
RDVISTIIPLPVIKELYRTDGCHILKFPQPHVVQvsqsaWMTDEEFareMIAGVNPCVIR   360
GLEEFPPKSNLDPAIYGDQSSKITADSLDLdgytmDEALGSRRLFMLDYHDIFMPYVRQI   420
NQLnsaktYATRTILFLREDGTLKPVAIELSLPHSAGDLSAavsqVVLPAKEGVESTIWL   480
LAKAYVIVNDSCYHQLMSHWLNTHAAMEPFVIATHRHLSVLHPIYKLLTPHYRNNMNINA   540
LARQSAINANGIIETTFLPSKYSVEMSSAVYKNWVFTDQALPADLIKRGVAIKDPSTPHG   600
VRLLIEDYPYAADGLEIWAAIKTwvqeYVPLYYARDDDVKNDSELqhwWKEAVEKGHGDL   660
KDKPWWPKLQTLEDlvevclIIIWIASALHAAVNFGQYPYGGLIMNRPTASRRLLPEKGT   720
PEYEEMINNHEKAYLRTITSKLptlISLSVIEILSTHASDEVYLGQRDNPHWTSDSKALQ   780
AFQKFGNKLKEIEEKLVRRNNDPSLQGNRLGPVQLPYTLLYPSSEEGLTFRGIPNSISI    839
```

5. **L754A-Q322C-BD**
   coverage: 816/839 residues, 97.26%

```
MFSAGHKIKGTVVLMPKNELEVNPDGSAVDNLNAFLGRSVSLQLISATKADAHGKGKVGK    60
DTFLEGINTSLPTLGAGESAFNIHFEWDGSMGIPGAFYIKNYMQVEFFLKSLTLEAISNQ   120
GTIRFVCNSWVYNTKLYKSVRIFFANHTYVPSETPAPLVEYREEELKSLRGNGTGERKEY   180
DRIYDYDVYNDLGNPDKSEKLARPVLGGSSTFPYPRRGRTGRGPTVTDPNTEKQGEVFYV   240
PRDENLGHLKSKDALEIGTKSLSQIVQPAFESAFDLKSTPIEFHSFQDVHDLYEGGIKLP   300
RDVISTIIPLPVIKELYRTDGCHILKFPQPHVVQVSQSAWMTDEEFAREMIAGVNpcvir   360
gLEEFPPKSNLDPAIYGDQSSKITADSLDLdgytmDEALGSRRLFMLDYHDIFMPYVRQI   420
NQLNSAKTYATRTILFLREDGTLKPVAIELSLPHSAGDLSAAVSQVVLPAKEGVESTIWL   480
LAKAYVIVNDSCYHQLMSHWLNTHAAMEPFVIATHRHLSVLHPIYKLLTPHYRNNMNINA   540
LARQSLINANGIIETTFLPSKYSVEMSSAVYKNWVFTDQALPADLIKRGVAIKDPSTPHG   600
VRLLIEDYPYAADGLEIWAAIKTWVQEYVPLYYARDDDVKNDSELQHWWKEAVEKGHGDL   660
KDKPWWPKLQTLEDlvevclIIIWIASALHAAVNFGQYPYGGLIMNRPTASRRLLPEKGT   720
PEYEEMINNHEKAYLRTITSKLptlISLSVIEIASTHASDEVYLGQRDNPHWTSDSKALQ   780
AFQKFGNKLKEIEEKLVRRNNDPSLQGNRLGPVQLPYTLLYPSSEEGLTFRGIPNSISI    839
```

**Fig. S6, continued.**

6. **I553A/L754A-Q322C-BD**
   coverage: 812/839 residues, 96.78%

```
MFSAGHKIKGTVVLMPKNELEVNPDGSAVDNLNAFLGRSVSLQLISATKADAHGKGKVGK    60
DTFLEGINTSLPTLGAGESAFNIHFEWDGSMGIPGAFYIKNYMQVEFFLKSLTLEAISNQ   120
GTIRFVCNSWVYNTKLYKSVRIFFANHTYVPSETPAPLVEYREEELKSLRGNGTGERKEY   180
DRIYDYDVYNDLGNPDKSEKLARPVLGGSSTFPYPRRGRTGRGPTVTDPNTEKQGEVFYV   240
PRDENLGHLKSKDALEIGTKSLSQIVQPAFESAFDLKSTPIEFHSFQDVHDLYEGGIKLP   300
RDVISTIIPLPVIKELYRTDGCHILKFPQPHVVQvsqsaWMTDEEfaremIAGVNpCVIR   360
GLEEFPPKSNLDPAIYGDQSSKITADSLDLdgytmdEALGSRRLFMLDYHDIFMPYVRQI   420
NQLNSAKTYATRTILFLREDGTLKPVAIELSLPHSAGDLSAAVSQVVLPAKEGVESTIWL   480
LAKAYVIVNDSCYHQLMSHWLNTHAAMEPFVIATHRHLSVLHPIYKLLTPHYRNNMNINA   540
LARQSLINANGIAETTFLPSKYSVEMSSAVYKNWVFTDQALPADLIKRGVAIKDPSTPHG   600
VRLLIEDYPYAADGLEIWAAIKTWVQEYVPLYYARDDDVKNDSELQHwWKEAVEKGHGDL   660
KDKPWWPKLQTLEDLVEvclIIIWIASALHAAVNFGQYPYGGLIMNRPTASRRLLPEKGT   720
PEYEEMINNHEKAYLRTITSKLptlISLSVIEIASTHASDEVYLGQRDNPHWTSDSKALQ   780
AFQKFGNKLKEIEEKLVRRNNDPSLQGNRLGPVQLPYTLLYPSSEEGLTFRGIPNSisi    839
```

7. **L546A/L754A-Q322C-BD**
   coverage: 792/839 residues, 94.4%

```
MFSAGHKIKGTVVLMPKNELEVNPDGSAVDNLNAFLGRSVSLQLISATKADAHGKGKVGK    60
DTFLEGINTSLPTLGAGESAFnihfEWDGSMGIPGAFYIKNYMQVEFFLKSLTLEAISNQ   120
GTIRFvcnsWVYNTKLYKSVRIFFANHTYVPSETPAPLVEYREEELKSLRGNGTGERKEY   180
DRIYDYDVYNDLGNPDKSEKLARPVLGGSSTFPYPRRGRTGRGPTVTDPNTEKQGEVFYV   240
PRDENLGHLKSKDALEIGTKSLSQIVQPAFESAFDLKSTPIEFHSFQDVHDLYEGGIKLP   300
RDVISTIIPLPVIKELYRTDGCHILKFPQPHVVQvsqsaWMTDEEfareMIAGVNPCVIR   360
GLEEFPPKSNLDPAIYGDQSSKITADSLDLdgytmdEALGSRRLFMLDYHDIFMPYVRQI   420
NQLNSAKTYATRTILFLREDGTLKPVAIELSLPHSAGDLSAavsqVVLPAKEGVESTIWL   480
lakaYVIVNDSCYHQLMSHWLNTHAAMEPFVIATHRHLSVLHPIYKLLTPHYRNNMNINA   540
LARQSAINANGIIETTFLPSKYSVEMSSAVYKNWVFTDQALPADLIKRGVAIKDPSTPHG   600
VRLLIEDYPYAADGLEIWAAIKTwvqeYVPLYYARDDDVKNDSELqhwWKEAVEKGHGDL   660
KDKPWWPKLQTLEDlvevclIIIWIASALHAAVNFGQYPYGGLIMNRPTASRRLLPEKGT   720
PEYEEMINNHEKAYLRTITSKLptlISLSVIEIASTHASDEVYLGQRDNPHWTSDSKALQ   780
AFQKFGNKLKEIEEKLVRRNNDPSLQGNRLGPVQLPYTLLYPSSEEGLTFRGIPNSISI    839
```

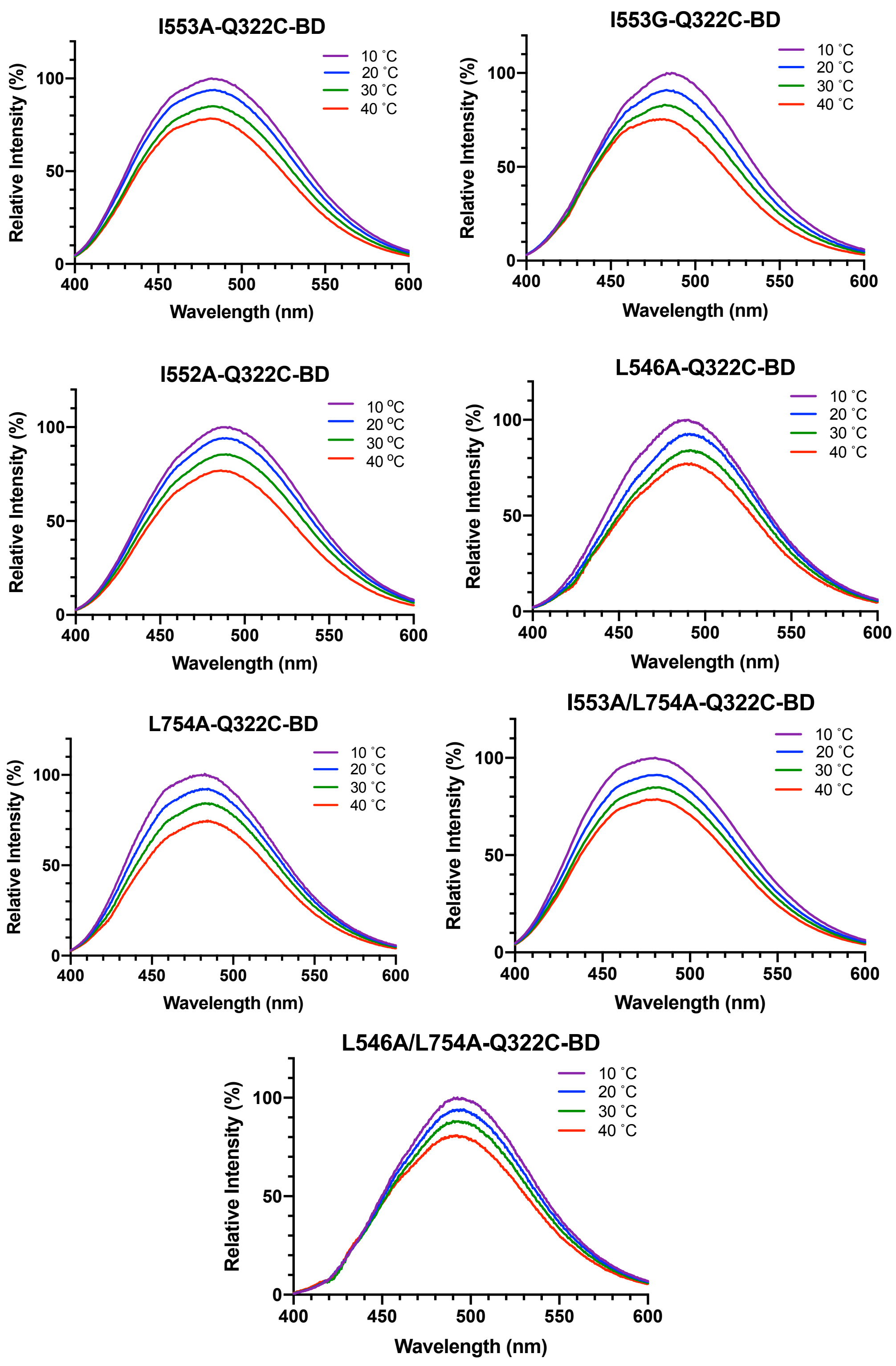


**Fig. S7.** Intensity-normalized steady-state fluorescence emission spectra ($\lambda_{exc}$ = 373 nm) collected as a function of temperature (10 – 40 ˚C) for the BD-labeled SLO series in 0.1 M pH 9.0 borate.

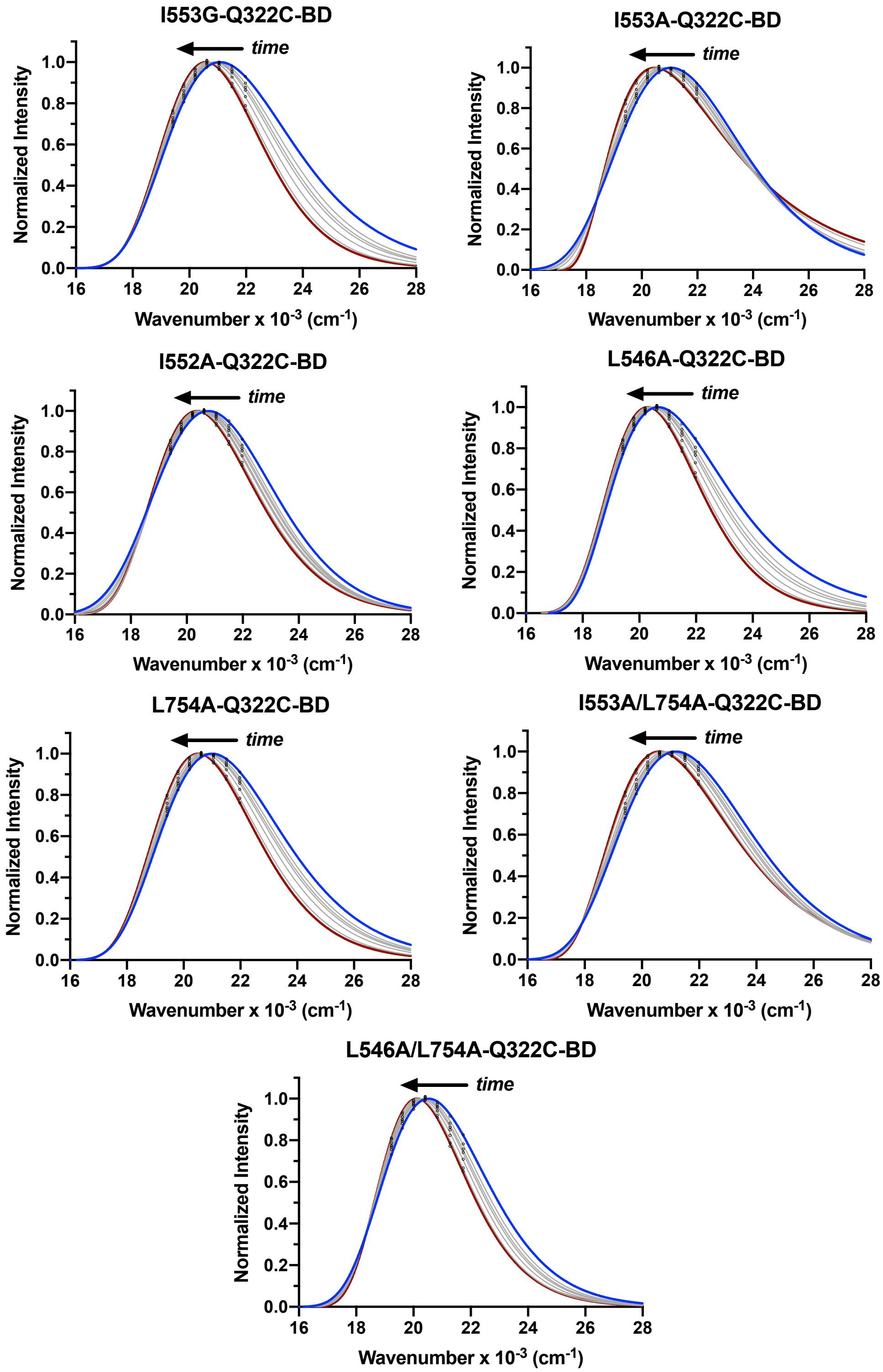


**Fig. S8.** Selected time-resolved emission spectra (TRES) for BD-labeled SLO from 0 ns (blue) to 10 ns (red) in 0.1 M pH 9.0 borate buffer at 30 ˚C.

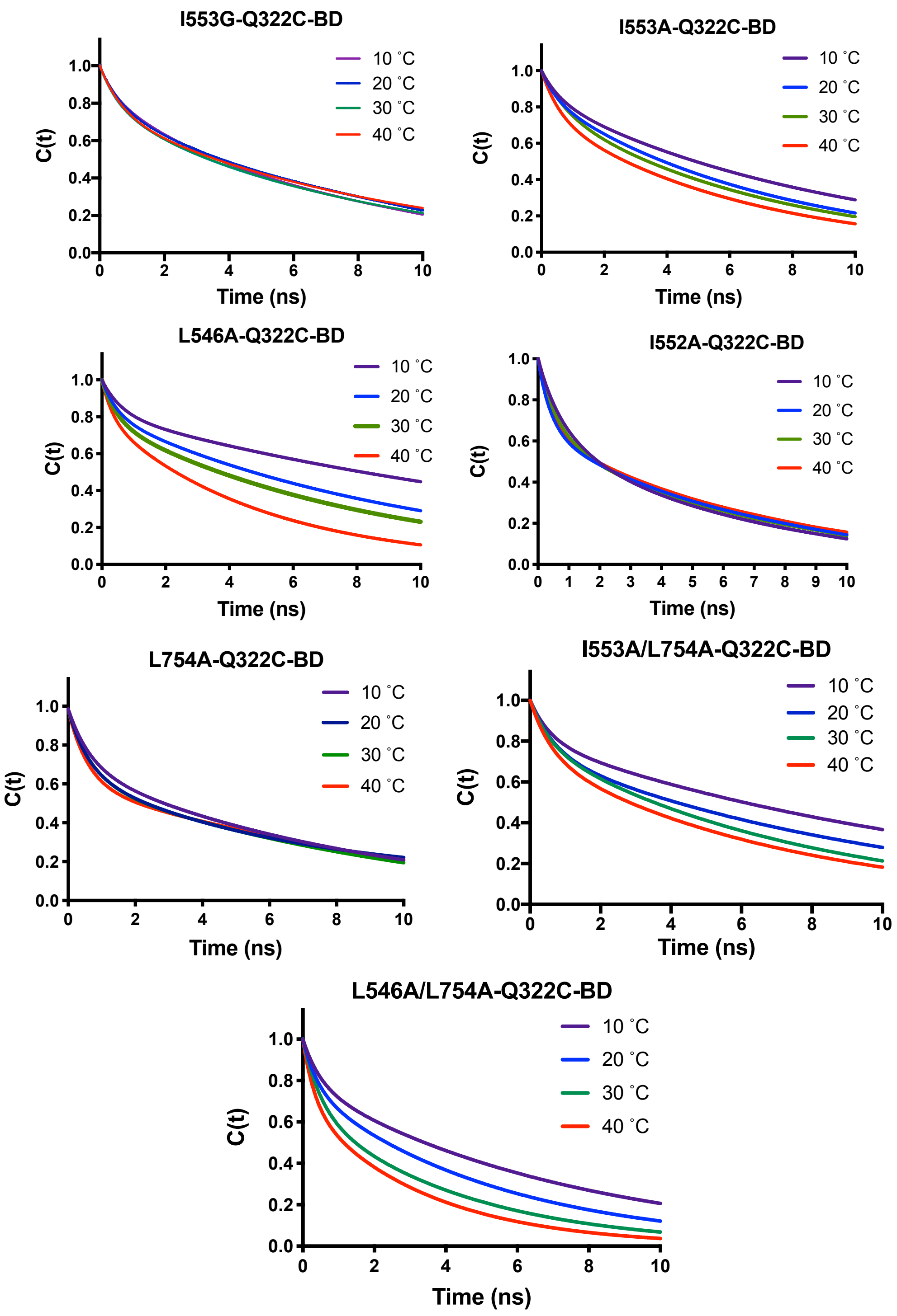


**Fig. S9.** Stokes Shift decay curves collected as a function of temperature (10 – 40 ˚C) for BD-labeled SLO, normalized in terms of the solvation correlation function C(t). Decays were fit to a biexponential function.

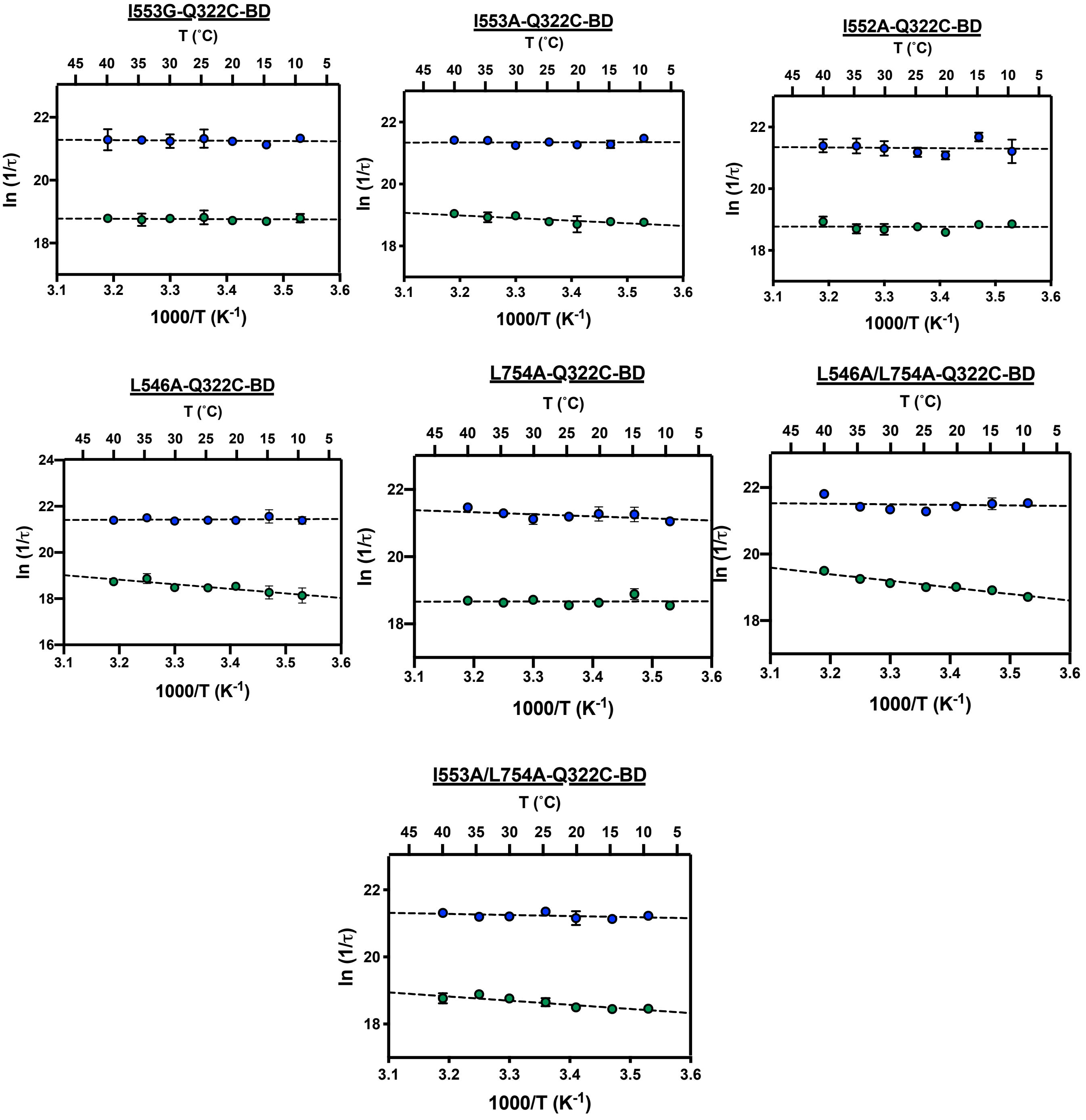


**Fig. S10.** Arrhenius plots of the temperature-dependent Stokes shift relaxation rates ($1/\tau_1$ in blue and $1/\tau_2$ in green) extracted from Figure S8.

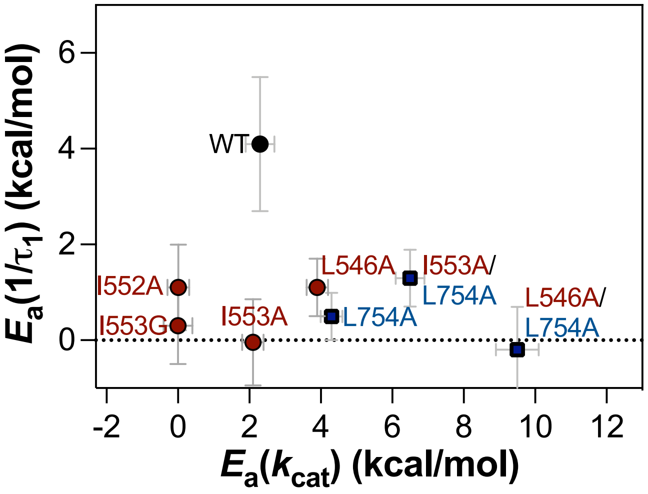


**Fig. S11.** Plot of activation energies of the Stokes shift decay rates for the fast $1/\tau_1$ component in the BD-labeled SLO variants.

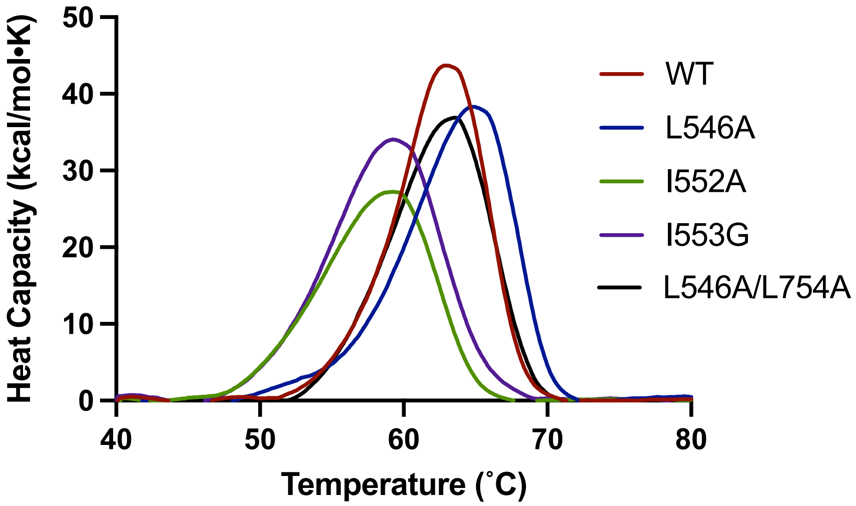


**Fig. S12**. Baseline corrected DSC thermograms for WT SLO and other variants in 50 mM borate pH 9 buffer SLO. The scan rate was 1 °C/min.

**Table S1**. Full tabulation of data from kinetic analyses of WT SLO and single and double mutants[a]

| SLO variant | $k_{cat}$ [b], $s^{-1}$ | ${}^{D}k_{cat}$ [b] | $E_a$, kcal/mol | $\Delta E_a$, kcal/mol | Reference |
|---|---|---|---|---|---|
| WT | 297 (10) | 81 (5) | 2.1 (0.2) | 0.9 (0.2) | Ref. 4 |
| *Active site single mutants* | | | | | |
| I553A | 280 (10) | 93 (4) | 1.9 (0.2) | 4.0 (0.3) | Ref. 4 |
| I553G | 58 (4) | 178 (16) | 0.03 (0.04) | 5.3 (0.7) | Ref. 5 |
| L546A | 4.8 (0.6) | 93 (9) | 4.1 (0.4) | 1.9 (0.6) | Ref. 4 |
| L754A | 0.31 (0.02) | 112 (11) | 4.1 (0.3) | 2.0 (0.5) | Ref. 4 |
| *Active site double mutants* | | | | | |
| I553A/L754A | 0.56 (0.03) | 85 (7) | 6.9(0.2) | 3.9(0.5) | Ref. 6 |
| L546A/L754A | 0.025 (0.01) | 692 (43) | 9.9(0.2) | 0.3(0.7) | Ref. 7 |
| *Connecting residue mutants* | | | | | |
| I552A | 81 (2)[c] | 60 (2)[c] | -0.3 (0.2) | 1.2 (0.5) | *This work* |
| V750A | 218 (8)[c] | 69 (4)[c] | 1.0 (0.4) | 0.3 (0.4) | *This work* |

[a]Kinetics were performed in 0.1 M borate, pH 9.0. [b]First-order rate constants ($k_{cat}$) and the corresponding deuterium isotope effects (${}^{D}k_{cat}$) are reported for 30°C [c]For comparison, under similar protein and substrate purification procedure, data for WT is $k_{cat}$ = 359 (7) $s^{-1}$, ${}^{D}k_{cat}$ = 57 (2), $E_a$ = 2.4 (0.2) kcal/mol, and $\Delta E_a$ = 1.1 (0.1) kcal/mol.

**Table S2.** Comparison of Activation Energies ($E_a$) for C-H activation ($E_a(k_{cat})$) and weighted average HDX exchange rates ($E_a(k_{HDX})$) at peptide 317-334 (thermally activated loop) for the I552A mutant (red). Data for other mutants shown from Refs. 1 and 8.

| SLO variant | $E_a$ ($k_{cat}$) (kcal/mol) | $E_a$ ($k_{HDX}$) (kcal/mol) |
|---|---|---|
| I552A | -0.3(0.2) | 3.2(3.7) |
| I553G | 0.03(0.04) | 0.68(1.2) |
| WT | 2.1(0.2) | 4.83(0.8) |
| L546A | 4.1(0.4) | 13.9(2.5) |
| L754A | 4.1(0.3) | 5.2(0.6) |
| L546A/L754A | 9.9(0.2) | 12.95(2.3) |

**Table S3.** X-ray Crystallography Data collection and refinement statistics

| | **I552A** | **V750A** |
|---|---|---|
| PDB ID: | 7SOI | 7SOJ |
| Wavelength (Å) | 0.88557 | 0.88557 |
| Temperature (K) | 300 | 300 |
| Space group | $P\ 2_1$ | $P\ 2_1$ |
| Cell Parameters | | |
| a b c (Å) | 91.531 92.759 99.836 | 91.537 92.693 100.270 |
| $\alpha\ \beta\ \gamma$ (˚) | 90.00 93.56 90.00 | 90.00 93.75 90.00 |
| Copies per a.s.u. | 2 | 2 |
| Resolution (Å) | 46.37-2.00 (2.03-2.00) | 46.35-1.85 (1.88-1.85) |
| No. Reflections | 590381 (15943) | 966364 (45698) |
| No. Unique | 109110 (4240) | 141110 (6896) |
| $R_{merge}$ | 0.254 (4.782) | 0.211 (2.462) |
| $R_{pim}$ | 0.183 (3.982) | 0.133 (1.596) |
| $I/\sigma I$ | 4.2 (0.3) | 5.7 (0.9) |
| Completeness (%) | 97.0 (75.7) | 99.1 (98.4) |
| Redundancy | 5.4 (3.8) | 6.8 (6.6) |
| $CC_{1/2}$ | 0.991 (0.105) | 0.992 (0.469) |
| **Refinement** | | |
| Resolution (Å) | 46.38 - 2.0 (2.072 - 2.0) | 45.67-1.85 (1.916-1.85) |
| Reflections | | |
| Total | 109082 (8818) | 141085 (13938) |
| Test | 5185 (337) | 6993 (699) |
| $R_{work}/R_{free}$ (%) | 19.70/25.75 (44.25/50.14) | 15.88/19.98 (31.37/35.15) |
| Number of non H atoms | 13966 | 14489 |
| Protein | 13445 | 13553 |
| Ligand | 3 | 2 |
| Solvent | 518 | 934 |
| Wilson B-factor | 33.21 | 24.42 |
| Average B-factor | 43.76 | 29.36 |
| Protein | 43.96 | 29.01 |
| Ligand | 60.91 | 29.05 |
| Solvent | 38.43 | 34.53 |
| Rmsd | | |
| Rmsd bond lengths | 0.012 | 0.010 |
| Rmsd bond angles | 1.18 | 1.05 |
| Ramachandran plot | | |
| Favored (%) | 96.06 | 96.62 |
| Allowed (%) | 3.94 | 3.32 |
| Outliers (%) | 0.00 | 0.8 |
| Clashscore | 3.77 | 2.14 |

Statistics for the highest-resolution shell are shown in parentheses.

**Table S4.** Calculated and measured molecular mass for labeled and unlabeled SLO constructs from mass spectrometry.

| SLO variant | (–) Badan | | (+) Badan | |
|---|---|---|---|---|
| | Calcd. Mass (Da) | Measured Mass (Da)[a] | Calcd. Mass (Da) | Measured Mass (Da)[a] |
| I553G – Q322C | 94,330 | 94,330 | 94,541 | 94,542 |
| I552A – Q322C | 94,344 | 94,345 | 94,555 | 94,556 |
| I553A – Q322C | 94,344 | 94,345 | 94,555 | 94,557 |
| L546A – Q322C | 94,344 | 94,344 | 94,555 | 94,556 |
| L754A – Q322C | 94,344 | 94,345 | 94,555 | 94,557 |
| I553A/L754A – Q322C | 94,302 | 94,303 | 94,513 | 94,513 |
| L546A/L754A – Q322C | 94,302 | 94,302 | 94,513 | 94,514 |

[a] Experimental uncertainty is ± 2 Da.

**Table S5.** Comparison of kinetic parameters for unlabeled and Badan-labeled SLO at Q322C.[a]

| SLO variant | Unlabeled | | Q322C-BD | |
|---|---|---|---|---|
| | $k_{cat}$ ($s^{-1}$) | $E_a$ (kcal/mol) | $k_{cat}$ ($s^{-1}$)[b] | $E_a$ (kcal/mol) |
| I552A | 81(2) | -0.3(0.2) | 83(4) | 0.01(0.3) |
| I553G | 58(4) | 0.03(0.04) | 51(1) | 0.003(0.4) |
| WT[b] | 297(12) | 2.1(0.2) | 252(13) | 2.3(0.4) |
| I553A | 280(10) | 1.9(0.2) | 235(10) | 2.1(0.3) |
| L546A | 4.8(0.6) | 4.1(0.4) | 4.6(0.2) | 3.9(0.3) |
| L754A | 0.31(0.02) | 4.1(0.3) | 0.35(0.01) | 4.3(0.3) |
| I553A/L754A | 0.56(0.03) | 6.9(0.2) | 0.45(0.07) | 6.5(0.4) |
| L546A/L754A | 0.021(0.001) | 9.9(0.2) | 0.021(0.0004) | 9.5(0.6) |

[a] 30 ˚C, 0.1 M borate buffer pH 9.0, Corrected for Fe content.
[b] WT data shown for comparison, data from Ref 3.

**Table S6.** Total Stokes shifts (Δν) and Stokes shift decay lifetimes ($\tau_1$, $\tau_2$) of SLO constructs at 30 ˚C, pH 9.0.

| SLO variant | $\nu_0$ (cm$^{-1}$) | Δν (cm$^{-1}$) | $\tau_1$ (ns) | $\alpha_1$ | $\tau_2$ (ns) | $\alpha_2$ |
|---|---|---|---|---|---|---|
| I552A – Q322C-BD | 21,121 | 944 | 0.50 | 0.54 | 8.21 | 0.46 |
| I553G – Q322C-BD | 21,631 | 1230 | 0.47 | 0.62 | 5.87 | 0.38 |
| Q322C-BD | 21,450 | 1173 | 0.41 | 0.57 | 4.70 | 0.43 |
| I553A – Q322C-BD | 21,395 | 1827 | 0.41 | 0.65 | 4.91 | 0.35 |
| L546A – Q322C-BD | 20,943 | 778 | 0.42 | 0.60 | 6.98 | 0.40 |
| L754A – Q322C-BD | 21,424 | 1205 | 0.48 | 0.60 | 6.75 | 0.40 |
| I553A/L754A – Q322C-BD | 21,530 | 1429 | 0.62 | 0.49 | 7.42 | 0.51 |
| L546A/L754A – Q322C-BD | 20,921 | 982 | 0.60 | 0.60 | 5.18 | 0.40 |

[a] WT data shown for comparison, data from Ref 3.

**Table S7.** Comparison of Fluorescence Decay Lifetimes at $\lambda_{max}$ emission ($\lambda_{max}$ excitation = 373 nm) of SLO constructs at 30 ˚C, pH 9.0[a,b,c]

| SLO variant | $\lambda_{max, em}$ (nm) | $\tau_1$ (ns) | $\alpha_1$ | $\tau_2$ (ns) | $\alpha_2$ | $\tau_3$ (ns) | $\alpha_3$ | $\tau_{ave}$ (ns) |
|---|---|---|---|---|---|---|---|---|
| I552A – Q322C-BD | 485(2) | 0.49 | 0.30 | 1.99 | 0.38 | 3.84 | 0.32 | 2.13 |
| I553G – Q322C-BD | 484(2) | 0.51 | 0.26 | 2.04 | 0.37 | 3.88 | 0.37 | 2.34 |
| Q322C-BD | 485(2) | 0.54 | 0.29 | 2.03 | 0.37 | 3.95 | 0.34 | 2.27 |
| I553A – Q322C-BD | 486(2) | 0.59 | 0.29 | 2.07 | 0.38 | 3.91 | 0.33 | 2.26 |
| L546A – Q322C-BD | 491(3) | 0.44 | 0.30 | 1.93 | 0.35 | 3.91 | 0.35 | 2.18 |
| L754A – Q322C-BD | 485(2) | 0.53 | 0.25 | 2.03 | 0.36 | 3.89 | 0.39 | 2.37 |
| I553A/L754A – Q322C-BD | 487(2) | 0.56 | 0.25 | 2.17 | 0.41 | 3.99 | 0.34 | 2.37 |
| L546A/L754A – Q322C-BD | 494(3) | 0.44 | 0.39 | 1.92 | 0.35 | 4.12 | 0.26 | 1.90 |

[a] Lifetimes were obtained at the respective $\lambda_{max}$(emission) for each sample.

[b] The amplitudes are reported such that $\alpha_1 + \alpha_2 + \alpha_3 = 1$.

[c] Lifetime errors are ±10% for $\tau_1$, and ±5% for $\tau_2$ and $\tau_3$.

**Table S8.** Comparison of Activation Energies ($E_a$, in kcal/mol) of Stokes Shift decays and C-H Activation for SLO constructs.

| **SLO variant** | $E_a(1/\tau_1)$ (kcal/mol) | $E_a(1/\tau_2)$ (kcal/mol) | $E_a(k_{cat})$ (kcal/mol) |
|---|---|---|---|
| I552A – Q322C-BD | 1.1(0.9) | 0.3(0.8) | 0.01(0.3) |
| I553G – Q322C-BD | 0.3(0.8) | 0.2(0.8) | 0.003(0.4) |
| Q322C-BD | 4.1(1.4) | 2.8(0.9) | 2.3(0.4) |
| I553A – Q322C-BD | -0.04(0.9) | 2.3(0.8) | 2.1(0.3) |
| L546A – Q322C-BD | 1.1(0.6) | 4.3(0.8) | 3.9(0.3) |
| L754A – Q322C-BD | 0.5(0.5) | 0.6(0.8) | 4.3(0.3) |
| I553A/L754A – Q322C-BD | 1.3(0.6) | 2.0(0.8) | 6.5(0.4) |
| L546A/L754A – Q322C-BD | -0.2(0.9) | 4.2(0.9) | 9.5(0.6) |

**Table S9**. DSC parameters of SLO variants

| **SLO variant** | **$T_m$ (˚C)** | **$\Delta T_m$ (˚C)[a]** | **ΔH˚** (kcal/mol) | **ΔΔH˚ (kcal/mol)[b]** |
|---|---|---|---|---|
| WT | 64.2 (0.1) | - | 311 (9) | - |
| L546A | 65.2 (0.2) | +1.0 (0.2) | 288 (10) | -23 (13) |
| I552A | 59.5 (0.1) | -4.7 (0.1) | 282 (3) | -29 (9) |
| I553G | 59.7 (0.7) | -4.5 (0.7) | 281 (14) | -30 (17) |
| L546A/L754A | 64.3 (0.6) | +0.1 (0.6) | 290 (2) | -21 (9) |

[a] $\Delta T_m = T_m(\text{mutant}) - T_m(\text{WT})$
[b] ΔΔH˚ = ΔH˚(mutant) – ΔH˚(WT)

**Table S10.** HDX data summary for I552A SLO

| Data sets | I552A SLO mutant |
|---|---|
| **HDX Reaction Details** | Labeling conditions: 10 μM protein, 90% $D_2O$, 10 mM HEPES pD = 7.4; corrected pD = $pH_{read}$ +0.4(65) to a pL of 8.0 (confirmed with pH electrode).<br>0s timepoint was collected with 10 μM protein, 10 mM HEPES pH = 7.0.<br>HDX was conducted at five different temperatures: 10, 20, 25, 30, 40 ˚C |
| **HDX Time Course** | 14 time points (0, 10, 30, 45, 60, 180, 600, 1200, 1800, 2700, 3600, 7200, 10800, 14400 s) for each temperature |
| **HDX Controls** | Maximally-labeled control (WT and I552A)<br>MS was validated in previous studies[1,8] |
| **Back Exchange: Average, Interquartile** | 29%, 16.5% |
| **Number of peptides** | 43 chosen for analysis |
| **Sequence coverage** | 87% of peptides were measurable (within the catalytic domain) |
| **Average Peptide Length** | 15 amides (6-26) |
| **Replicates** | 1 (biological) for each mutant and temperature<br>Each time point for each temperature for each protein was collected once.<br>Due to the scale of the experiment, it was impractical to collect multiple replicates at all points. To mitigate systematic errors, each temperature set (14 time points of one protein) was collected over three days in a non-sequential order.<br>The temperature dependence provides a check for outlying datasets relative to the WT protein. |
| **Repeatability** | There are no replicates, however, in majority of the peptides, the data points for WT and I552A are nearly identical. |
| **Significant Differences in HDX** | In this study, we are interested in the changes in $E_a$(HDX) for a specific peptide 317-334 for the I552A mutant in relation to WT and other active site mutants as defined previously[1,8]. $E_a$(HDX) values that were larger than their error were considered a significant difference between protein states. |

**Table S11.** Back exchange values for on-overlapping peptide set for I552A. The mutation site (I552A) is marked in red. The peptide representing the thermally-activated surface loop is highlighted in bold.

| Peptide | Sequence | Charge | Back exchange % |
|---|---|---|---|
| 144-160 | FANHTYVPSETPAPLVE | 2 | 33 |
| 161-185 | YREEELKSLRGNGTGERKEYDRIYD | 3 | 48 |
| 186-206 | YDVYNDLGNPDKSEKLARPVL | 3 | 26 |
| 212-238 | FPYPRRGRTGRGPTVTDPNTEKQGEVF | 3 | 41 |
| 239-256 | YVPRDENLGHLKSKDALE | 2 | 49 |
| 257-273 | IGTKSLSQIVQPAFESA | 2 | 21 |
| 274-283 | FDLKSTPIEF | 2 | 17 |
| 284-299 | HSFQDVHDLYEGGIKL | 2 | 28 |
| 297-305 | IKLPRDVIS | 2 | 16 |
| 306-316 | TIIPLPVIKEL | 2 | 9 |
| **317-334** | **YRTDGQHILKFPQPHVVQ** | **3** | **36** |
| 347-355 | AREMIAGVN | 2 | 18 |
| 362-388 | LEEFPPKSNLDPAIYGDQSSKITADSL | 2 | 25 |
| 407-413 | LDYHDIF | 2 | 36 |
| 414-423 | MPYVRQINQL | 2 | 8 |
| 424-435 | NSAKTYATRT | 2 | 16 |
| 436-449 | FLREDGTLKPVAIE | 2 | 30 |
| 450-462 | LSLPHSAGDLSAA | 2 | 33 |
| 465-477 | QVVLPAKEGVEST | 2 | 31 |
| 493-503 | YHQLMSHWLNT | 2 | 43 |
| 504-521 | HAAMEPFVIATHRHLSVL | 3 | 42 |
| 522-540 | HPIYKLLTPHYRNNMNINA | 2 | 32 |
| 541-554 | LARQSLINANG**A**IE | 2 | 20 |
| 555-565 | TTFLPSKYSVE | 2 | 21 |
| 566-576 | MSSAVYKNWVF | 2 | 20 |
| 586-603 | IKRGVAIKDPSTPHGVRL | 3 | 37 |
| 604-617 | LIEDYPYAADGLEI | 2 | 20 |
| 619-632 | AAIKTWVQEYVPLY | 2 | 6 |
| 633-648 | YARDDDVKNDSELQHW | 2 | 42 |
| 649-669 | WKEAVEKGHGDLKDKPWWPKL | 3 | 48 |
| 690-703 | HAAVNFGQYPYGGL | 2 | 18 |
| 704-726 | IMNRPTASRRLLPEKGTPEYEEM | 3 | 45 |
| 727-737 | INNHEKAYLRT | 2 | 30 |
| 738-745 | ITSKLPTL | 2 | 13 |
| 751-761 | IEILSTHASDE | 2 | 32 |

| | | | |
|---|---|---|---|
| 762-781 | VYLGQRDNPHWTSDSKALQA | 3 | 45 |
| 782-796 | FQKFGNKLKEIEEKL | 2 | 35 |
| 797-820 | VRRNNDPSLQGNRLGPVQLPYTLL | 2 | 30 |
| 821-839 | YPSSEEGLTFRGIPNSISI | 2 | 25 |

**Table S12:** Comparison of apparent activation energies for HDX within the analyzed peptides from WT and I552A SLO.[a]

| Peptide | $E_{aHDX(avg)}$ WT | $E_{aHDX(avg)}$ I552A |
|---|---|---|
| 144-160 | | 8.0 (1.5) |
| 161-185 | 1.5 (3.0) | -2.8 (5.2) |
| 186-206 | 7.3 (9.6) | 0.2 (3.3) |
| 212-238 | 15.0 (3.2) | 12.5 (5.2) |
| 239-256 | N.D.[c] | N.D.[c] |
| 257-273 | N.D.[c] | N.D.[c] |
| 274-283 | N.D.[c] | N.D.[c] |
| 284-299[d] | 11.6 (2.1) | 10.8 (3.5) |
| 297-305[e] | 13.9 (4.1) | 13.0 (4.7) |
| 306-316[e] | 8.9 (4.6) | 8.7 (4.7) |
| **317-334[d]** | **4.8 (0.8)** | **3.2 (3.7)** |
| 347-355 | N.D.[c] | N.D.[c] |
| 362-388 | 16.6 (3.3) | 17.0 (5.1) |
| 407-413 | 15.7 (2.2) | 17.8 (0.3) |
| 414-423 | -7.8 (7.2) | -8.3 (3.7) |
| 424-435 | N.D.[c] | N.D.[c] |
| 436-449 | N.D.[c] | N.D.[c] |
| 450-462 | N.D.[c] | N.D.[c] |
| 465-477 | -11.6 (21.5) | -14.8 (8.3) |
| 485-503 | N.D.[c] | N.D.[c] |
| 493-503 | N.D.[c] | N.D.[c] |
| 504-521 | N.D.[c] | N.D.[c] |
| 522-540 | 13.1 (1.9) | 12.1 (0.6) |
| **541-554[e]** | **23.7 (2.0)** | **5.64 (0.5)** |
| **555-565[e]** | **17.1 (2.8)** | **5.1 (2.0)** |
| 566-576 | 18.1 (1.4) | 11.7 (0.7) |
| 586-603 | 12.1 (0.9) | 13.9 (1.0) |
| 604-618 | N.D.[c] | N.D.[c] |
| 619-632 | N.D.[c] | N.D.[c] |
| 633-648 | 6.8 (1.5) | 5.7 (2.2) |
| 649-669 | 14.8 (0.7) | 15.9 (1.0) |
| 690-703 | N.D.[c] | N.D.[c] |
| 704-726 | 0.8 (1.8) | 1.3 (1.8) |

| | | |
|---|---|---|
| 727-737 | 17.3 (1.0) | 13.7 (2.1) |
| 738-745 | 19.6 (2.0) | 19.7 (1.3) |
| 751-761[e] | 7.3 (2.1) | 15.1 (5.9) |
| 762-781 | 15.0 (1.2) | 16.3 (1.1) |
| 782-796 | 8.9 (3.7) | 9.4 (2.2) |
| 797-820 | 1.5 (2.1) | -2.5 (9.1) |
| 821-839 | 2.8 (1.7) | 0.2 (4.1) |

[a]*Apparent* activation energies, in kcal/mol, are determined from slopes of Arrhenius-like plots of the weighted average rate of HDX across five temperatures. The "()" represents the standard error calculated from the linear fits. Because the dynamic range does not capture the full exchange information, the enthalpies only reflect apparent enthalpies.
[b]N.C.: Peptide not covered by this mutation.
[c]N.D.: Not determined. There is not enough information within the dynamic range to provide an accurate determination of the weighted average rates of exchange.
[d]Altered $E_{a\mathrm{HDX(avg)}}$ in the mutants, statistically different based on reported error.
[e]Altered rates of exchange in mutants, but there is no discernable trend or difference in $E_{a\mathrm{HDX(avg)}}$.
[f]Altered $E_{a\mathrm{HDX(avg)}}$ in the mutants; determined from three temperatures, 10-25˚C, only.
