## Supplementary material for "Temporal and Spatial Resolution of a Protein Quake that Activates Hydrogen Tunneling in Soybean Lipoxygenase": Data S1

**Data S1:** HDX-MS deuterium uptake plots for all peptides of I552A SLO, in comparison to WT SLO, from Offenbacher, et al., *ACS Cent. Sci.* **2017**, *3*, 570. Each peptide has been corrected for back exchange individually. HDX is plotted as % deuterium uptake vs. time in minutes. Temperatures are indicated by colored lines and points (40 ˚C red, 30˚C yellow, 25 ˚C gray, 20 ˚C green, and 10 ˚C blue).

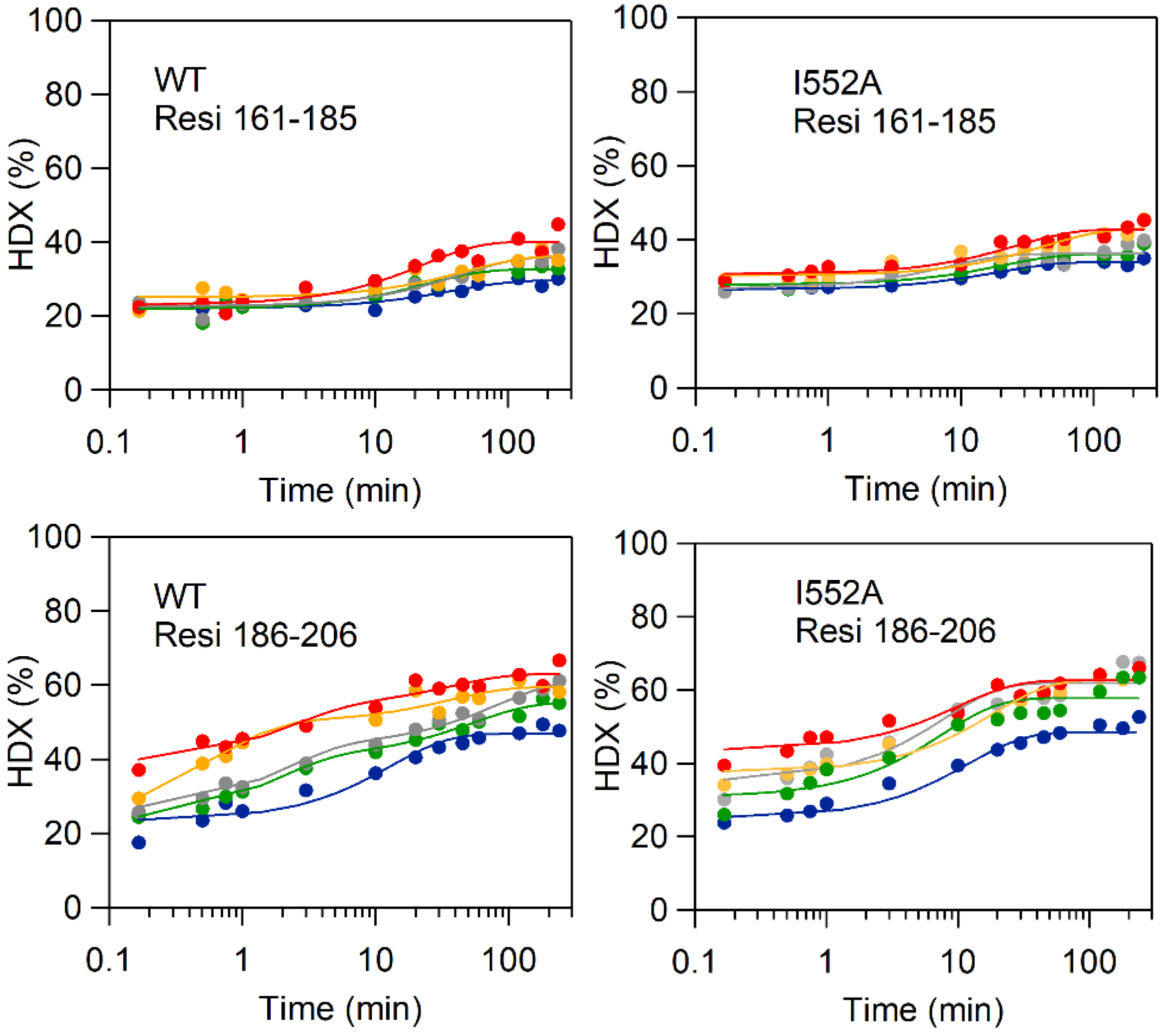

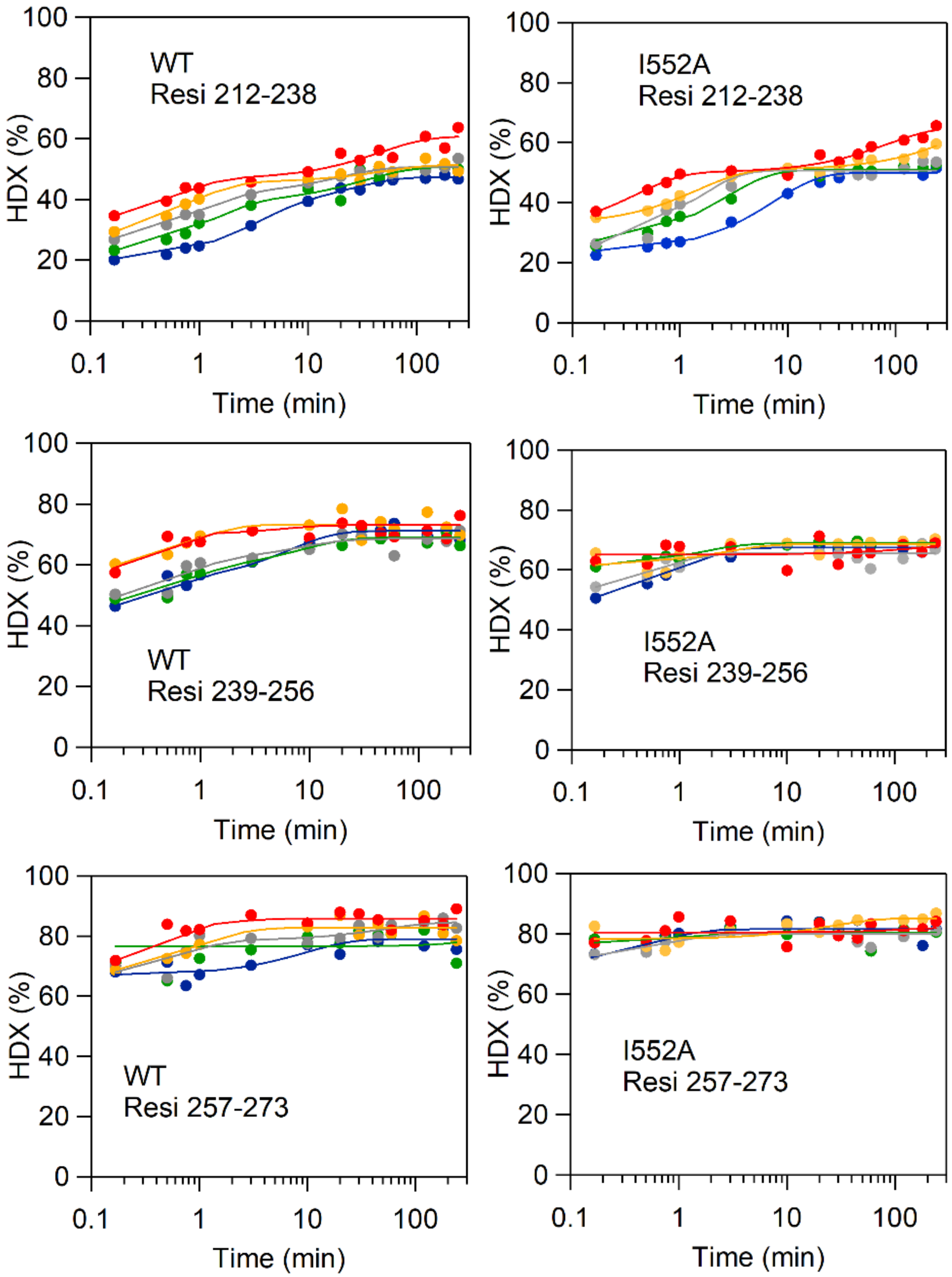
WT
Resi 212-238
I552A
Resi 212-238
WT
Resi 239-256
I552A
Resi 239-256
WT
Resi 257-273
I552A
Resi 257-273
HDX (%)
Time (min)
0
20
40
60
80
100
0.1
1
10
100

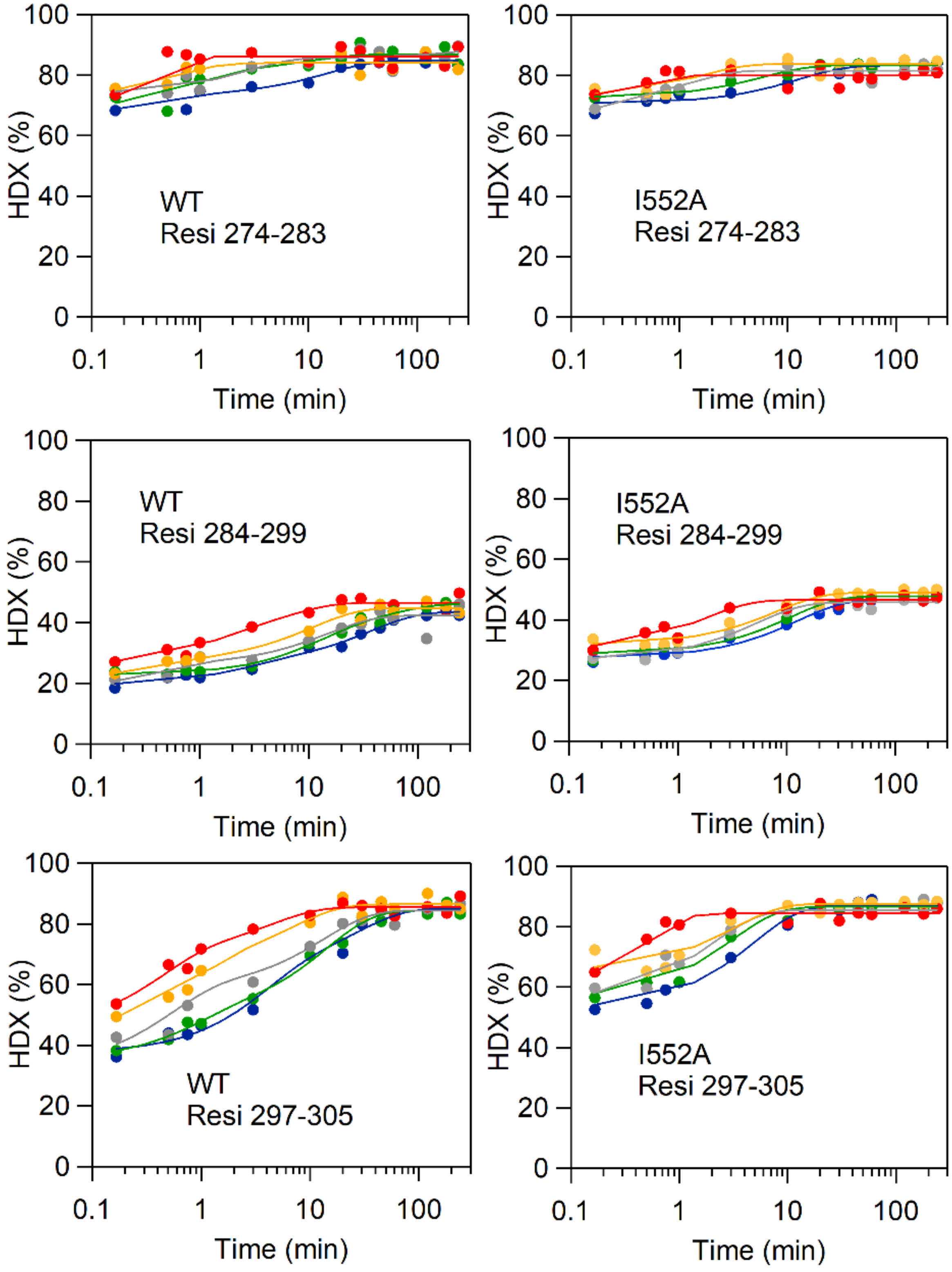
WT
Resi 274-283
I552A
Resi 274-283
WT
Resi 284-299
I552A
Resi 284-299
WT
Resi 297-305
I552A
Resi 297-305
HDX (%)
Time (min)
0
20
40
60
80
100
0.1
1
10
100

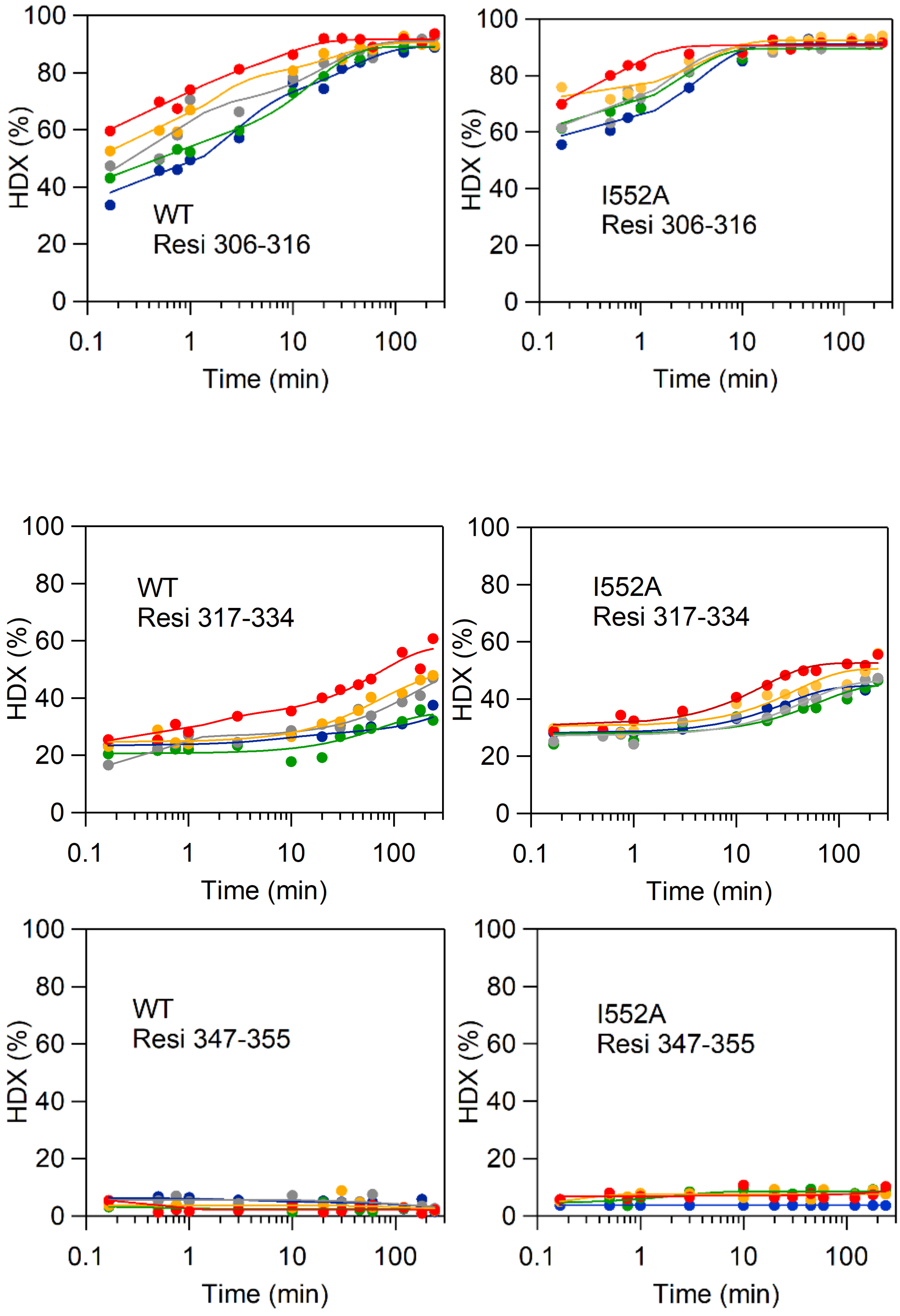
WT
Resi 306-316
I552A
Resi 306-316
WT
Resi 317-334
I552A
Resi 317-334
WT
Resi 347-355
I552A
Resi 347-355
HDX (%)
Time (min)
0
20
40
60
80
100
0.1
1
10
100

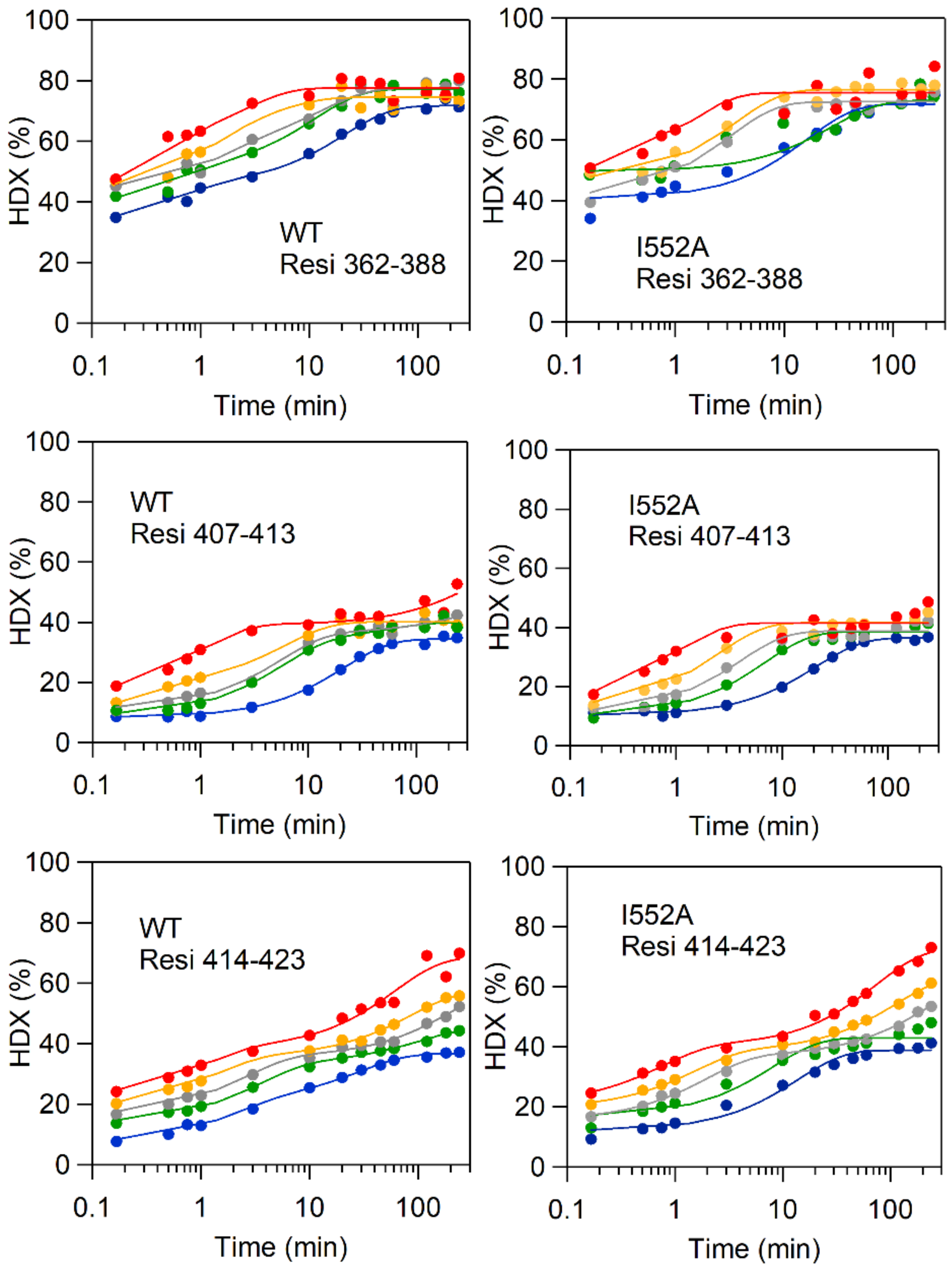
WT
Resi 362-388
I552A
Resi 362-388
WT
Resi 407-413
I552A
Resi 407-413
WT
Resi 414-423
I552A
Resi 414-423
HDX (%)
Time (min)
0
20
40
60
80
100
0.1
1
10
100

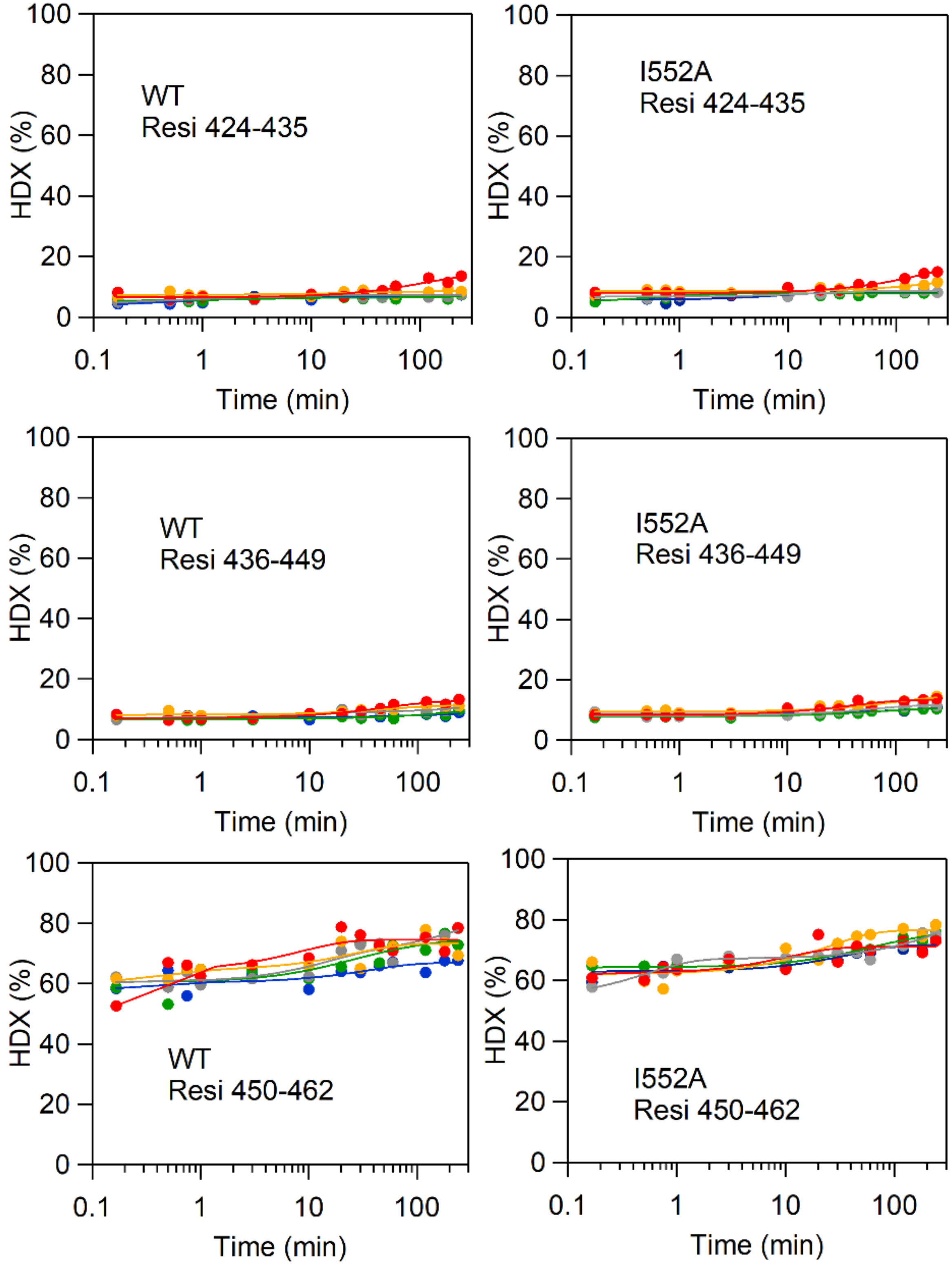
WT
Resi 424-435
I552A
Resi 424-435
WT
Resi 436-449
I552A
Resi 436-449
WT
Resi 450-462
I552A
Resi 450-462
HDX (%)
Time (min)
0
20
40
60
80
100
0.1
1
10
100

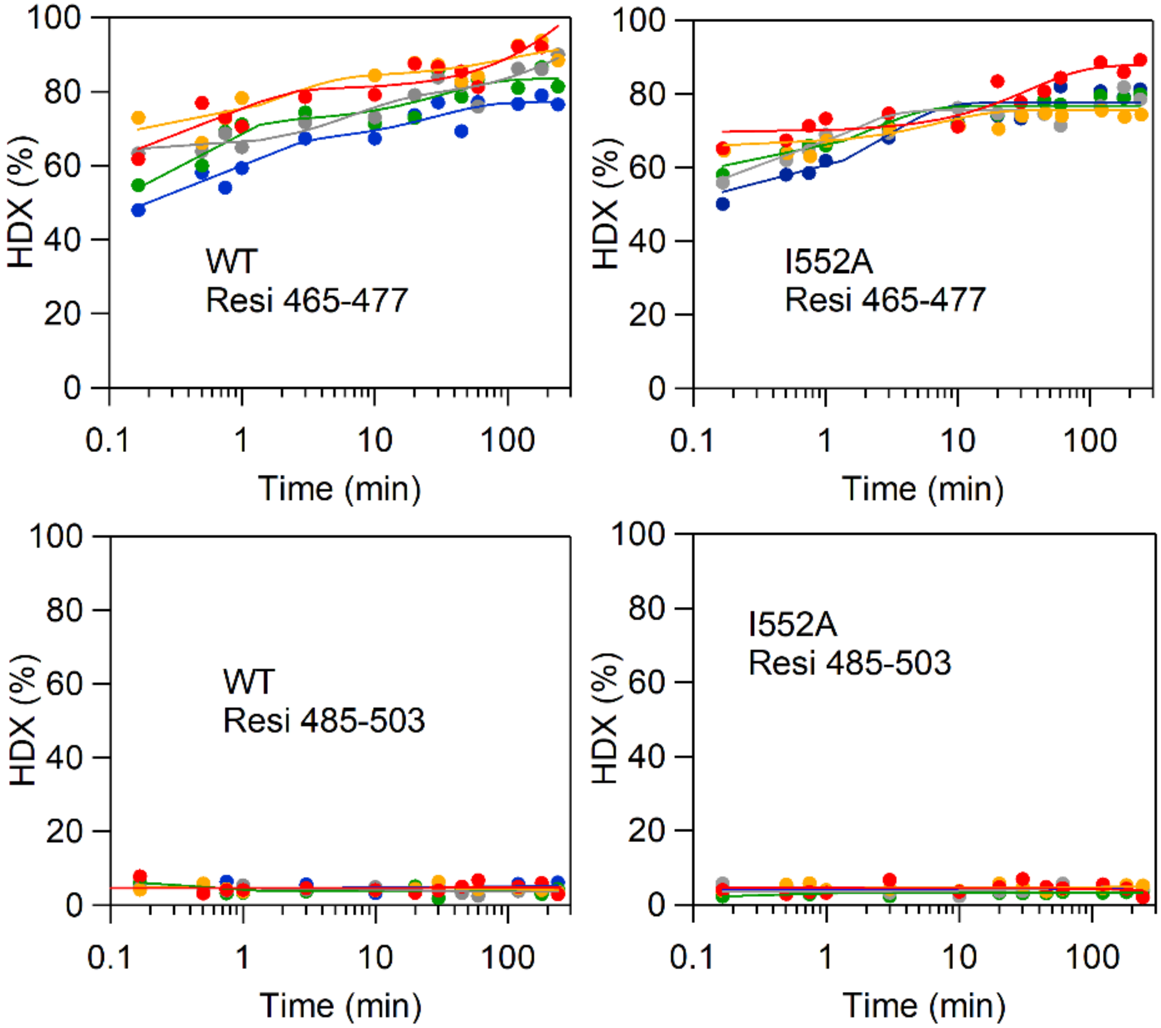
100
80
60
40
20
0
HDX (%)
0.1
1
10
100
Time (min)
WT
Resi 465-477
I552A
Resi 465-477
WT
Resi 485-503
I552A
Resi 485-503

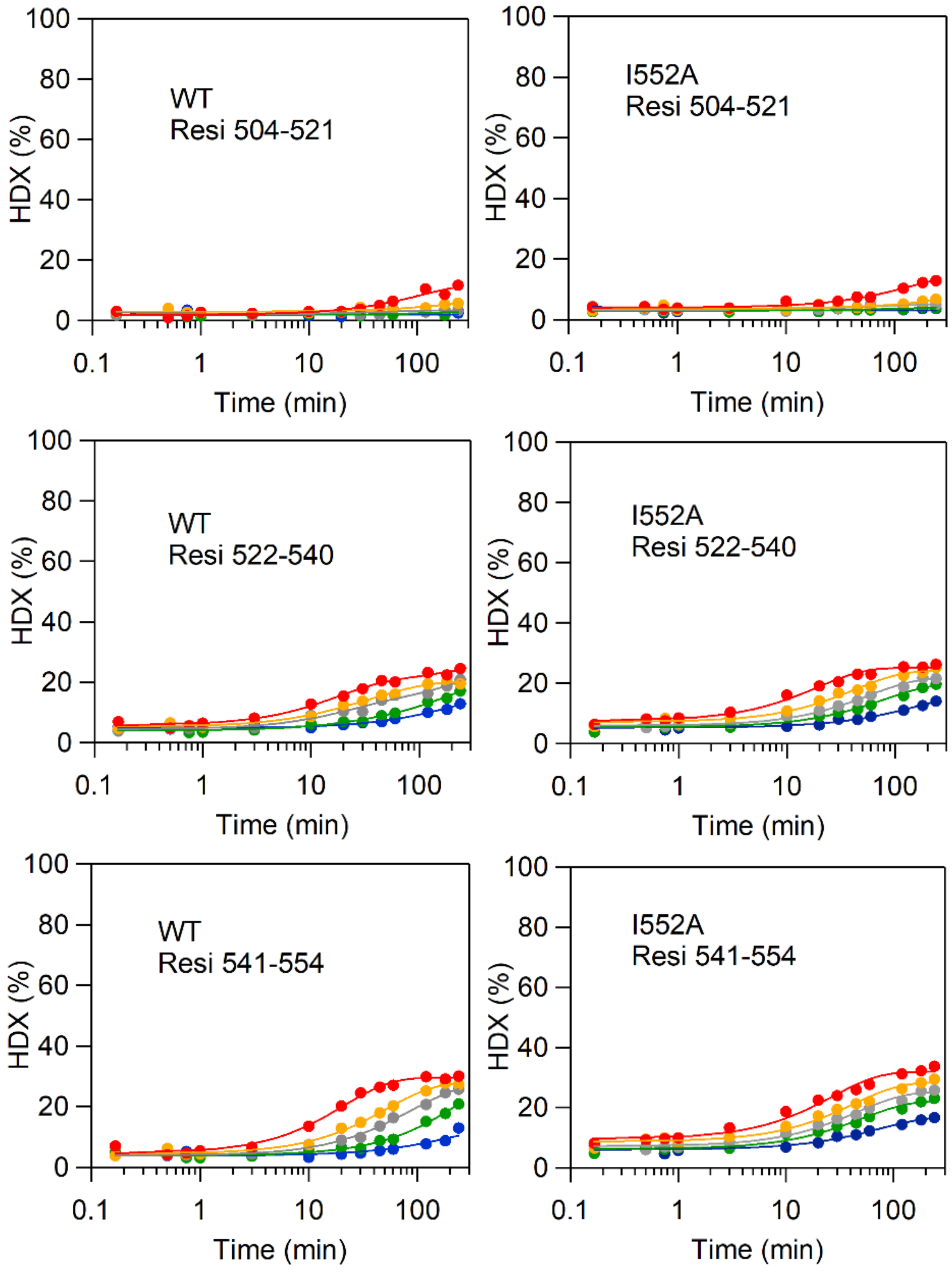
WT
Resi 504-521
I552A
Resi 504-521
WT
Resi 522-540
I552A
Resi 522-540
WT
Resi 541-554
I552A
Resi 541-554
HDX (%)
Time (min)
0
20
40
60
80
100
0.1
1
10
100

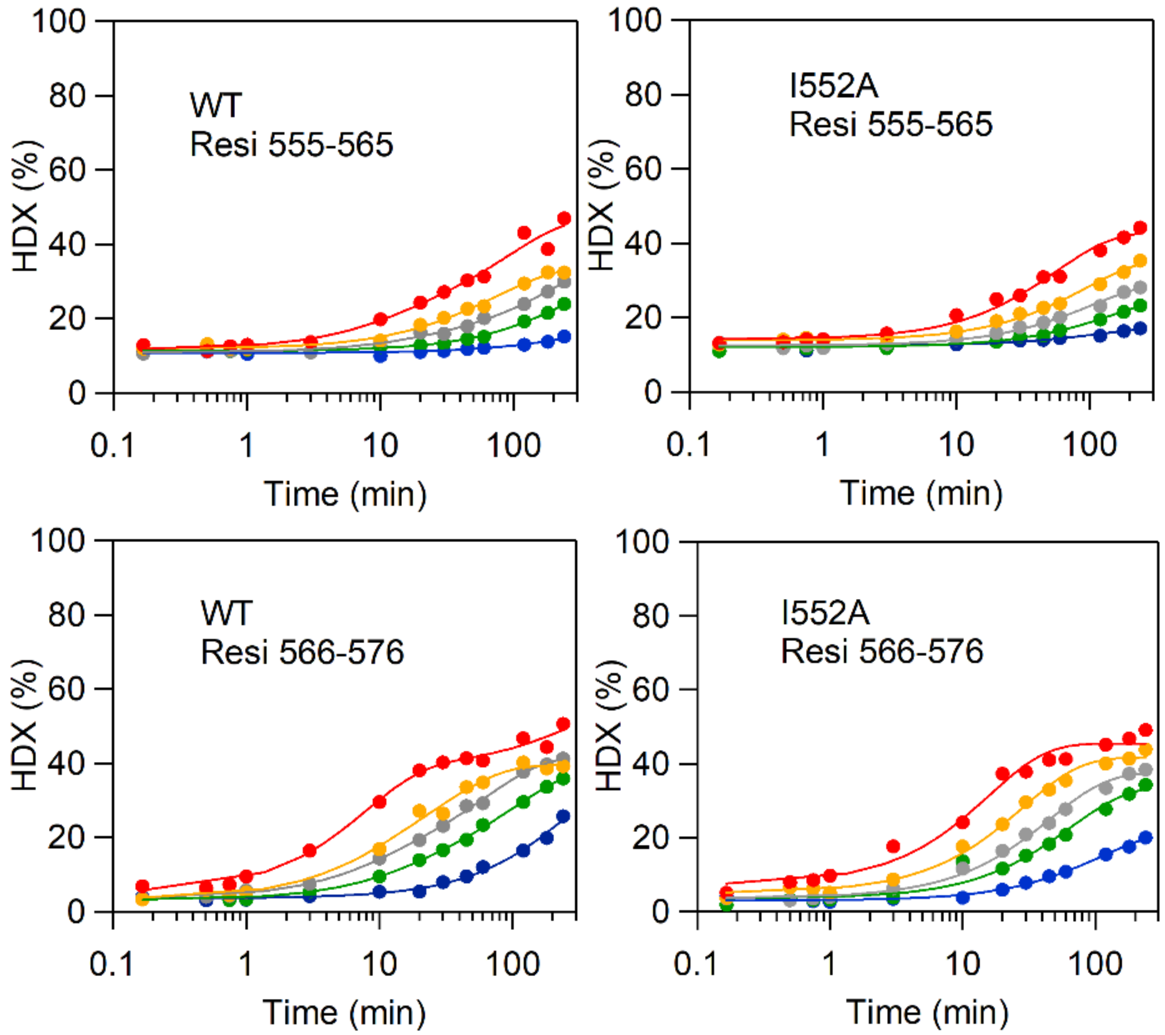
HDX (%)
Time (min)
WT
Resi 555-565
I552A
Resi 555-565
WT
Resi 566-576
I552A
Resi 566-576

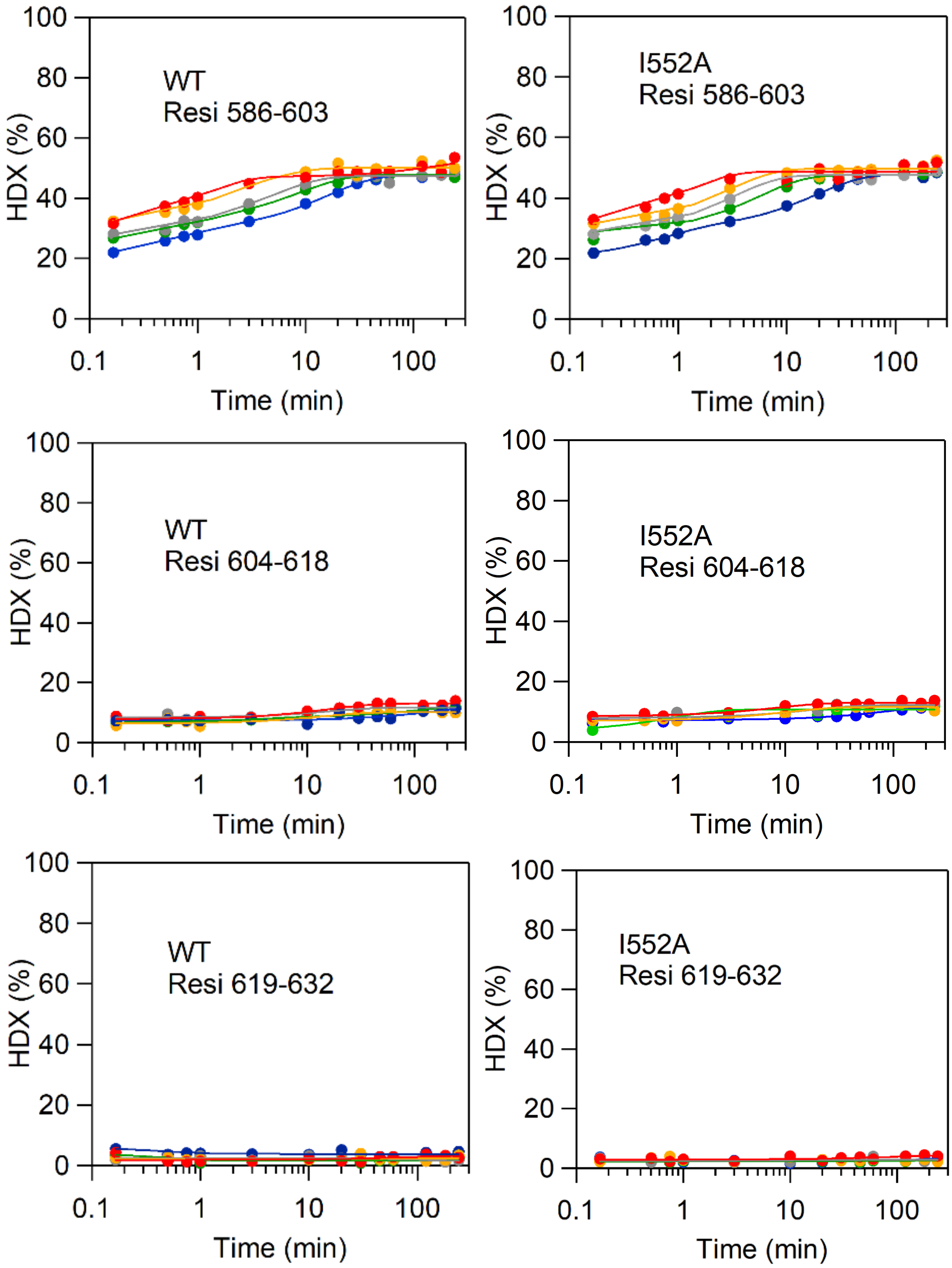
WT
Resi 586-603
I552A
Resi 586-603
WT
Resi 604-618
I552A
Resi 604-618
WT
Resi 619-632
I552A
Resi 619-632
HDX (%)
Time (min)

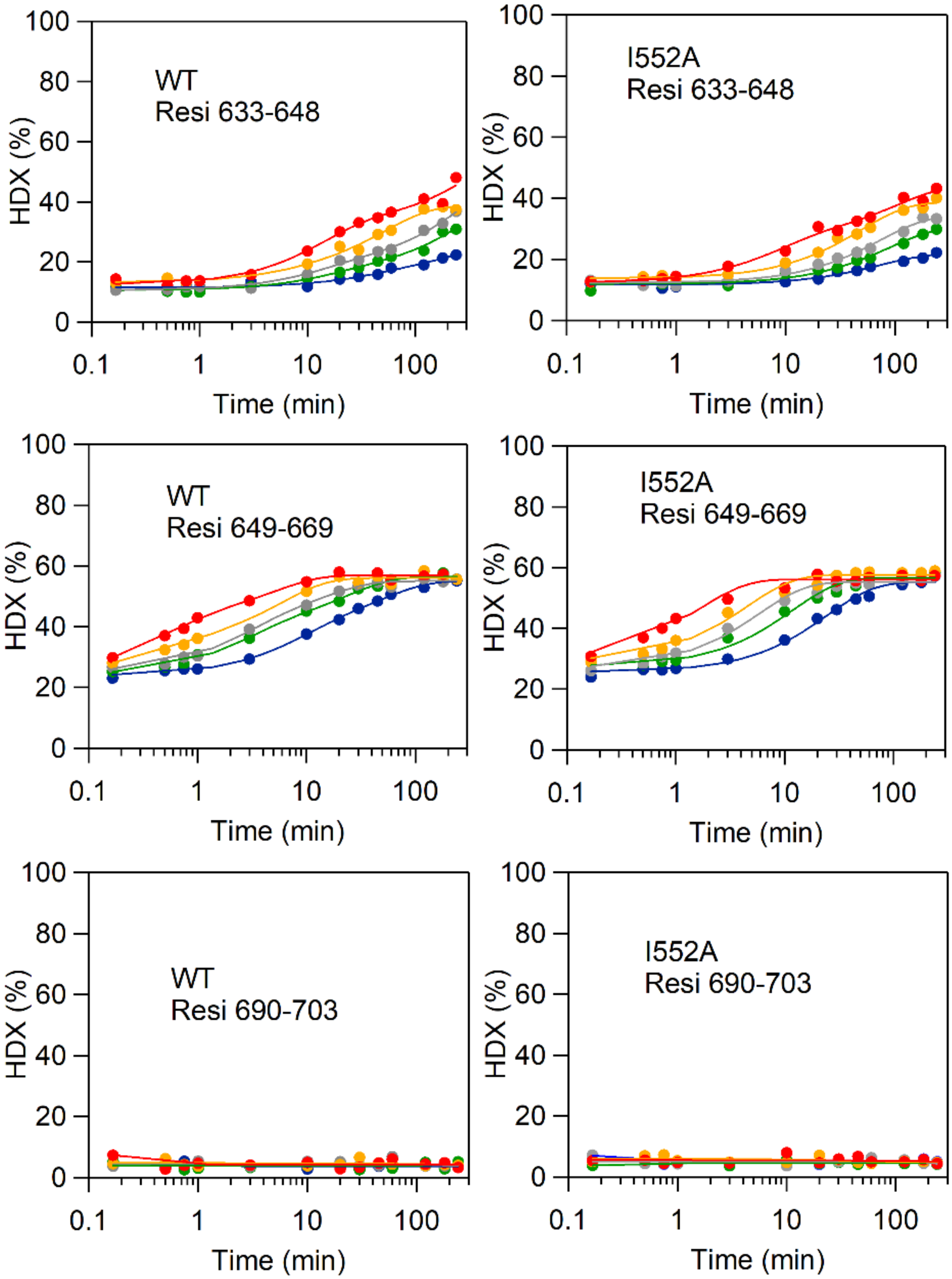
WT
Resi 633-648
I552A
Resi 633-648
WT
Resi 649-669
I552A
Resi 649-669
WT
Resi 690-703
I552A
Resi 690-703
HDX (%)
Time (min)
0
20
40
60
80
100
0.1
1
10
100

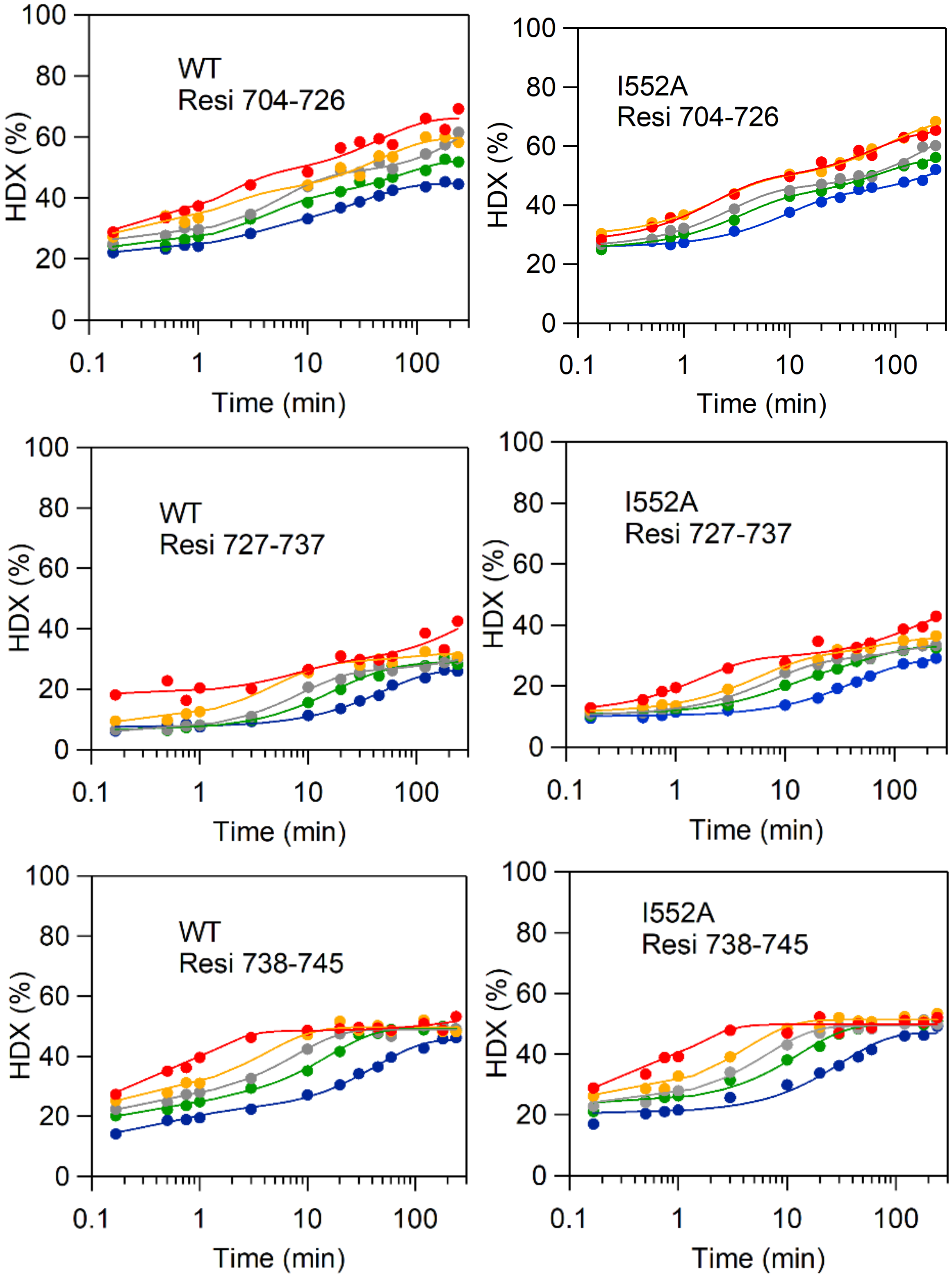
WT
Resi 704-726
I552A
Resi 704-726
WT
Resi 727-737
I552A
Resi 727-737
WT
Resi 738-745
I552A
Resi 738-745
HDX (%)
Time (min)
0
20
40
60
80
100
0.1
1
10
100

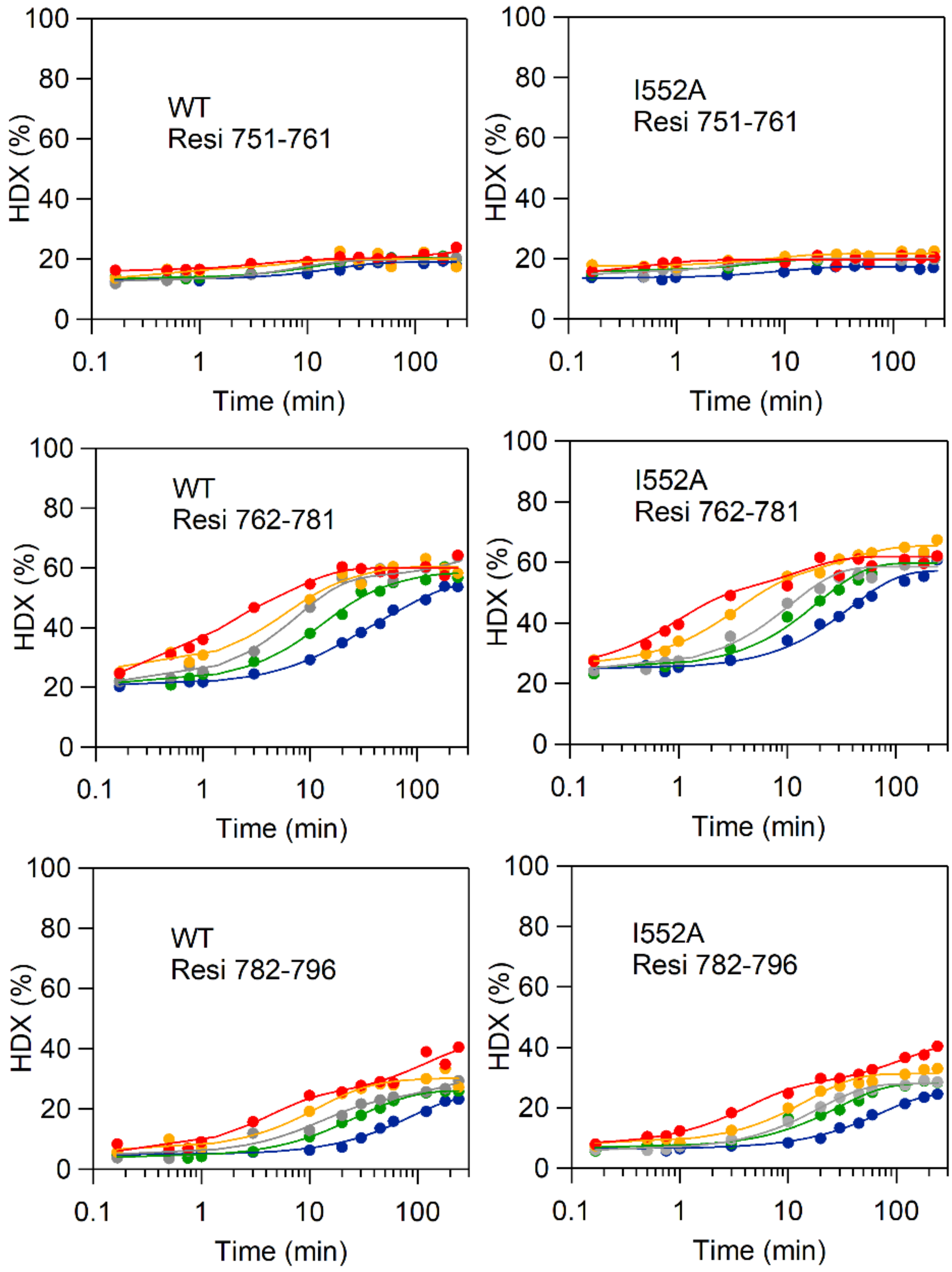
WT
Resi 751-761
I552A
Resi 751-761
WT
Resi 762-781
I552A
Resi 762-781
WT
Resi 782-796
I552A
Resi 782-796
HDX (%)
Time (min)
100
80
60
40
20
0
0.1
1
10
100

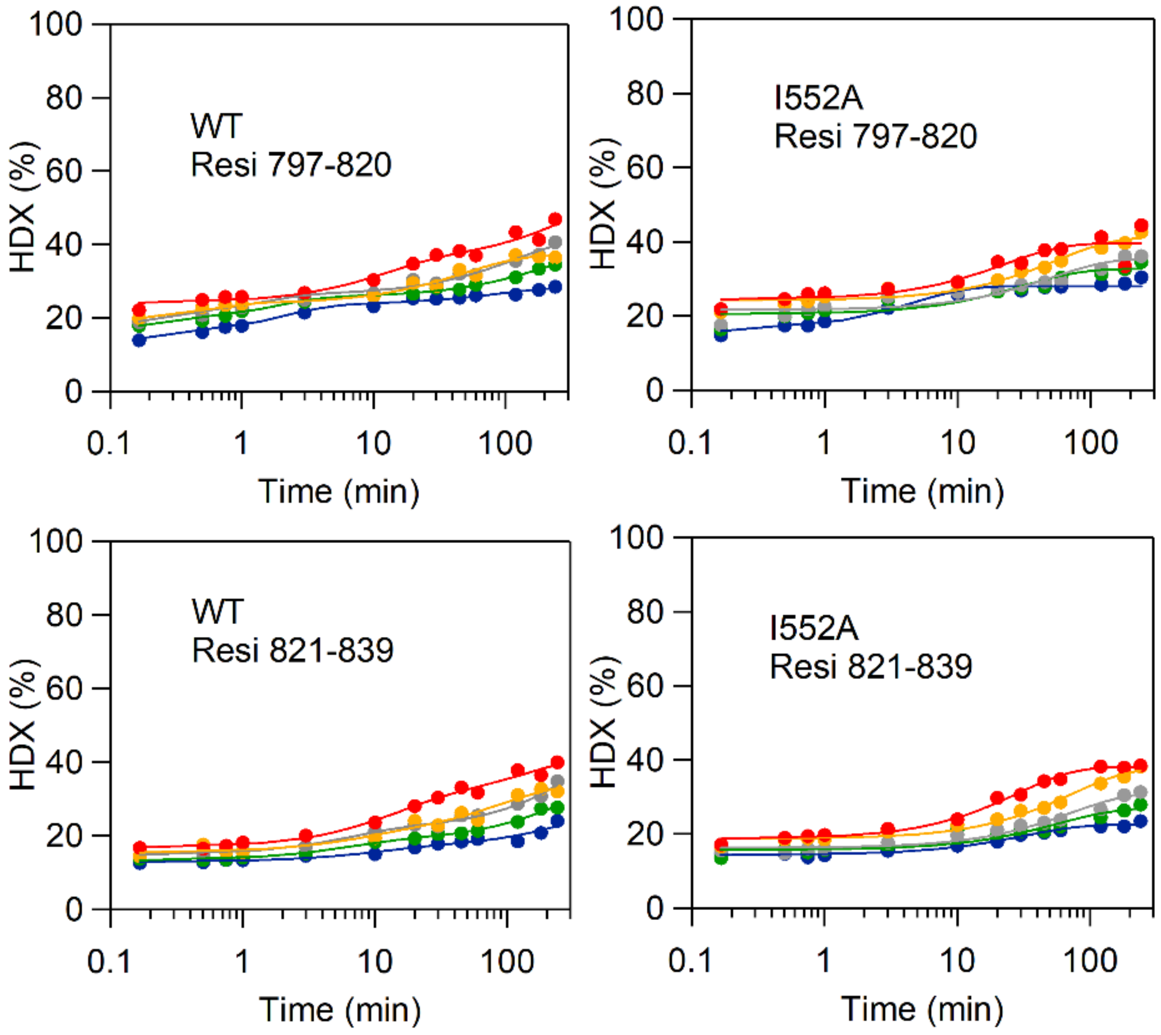
WT
Resi 797-820
I552A
Resi 797-820
WT
Resi 821-839
I552A
Resi 821-839
HDX (%)
Time (min)
0
20
40
60
80
100
0.1
1
10
100
